# Adaptation of a quantitative trait to a changing environment: new analytical insights on the asexual and infinitesimal sexual models

**DOI:** 10.1101/2022.06.22.497192

**Authors:** J. Garnier, O. Cotto, E. Bouin, T. Bourgeron, T. Lepoutre, O. Ronce, V. Calvez

## Abstract

Predicting the adaptation of populations to a changing environment is crucial to assess the impact of human activities on biodiversity. Many theoretical studies have tackled this issue by modeling the evolution of quantitative traits subject to stabilizing selection around an optimal phenotype, whose value is shifted continuously through time. In this context, the population fate results from the equilibrium distribution of the trait, relative to the moving optimum. Such a distribution may vary with the shape of selection, the system of reproduction, the number of loci, the mutation kernel or their interactions. Here, we develop a methodology that provides quantitative measures of population maladaptation and potential of survival directly from the entire profile of the phenotypic distribution, without any a priori on its shape. We investigate two different systems of reproduction (asexual and infinitesimal sexual models of inheritance), with various forms of selection. In particular, we recover that fitness functions such that selection weakens away from the optimum lead to evolutionary tipping points, with an abrupt collapse of the population when the speed of environmental change is too high. Our unified framework allows deciphering the mechanisms that lead to this phenomenon. More generally, it allows discussing similarities and discrepancies between the two systems of reproduction, which are ultimately explained by different constraints on the evolution of the phenotypic variance. We demonstrate that the mean fitness in the population crucially depends on the shape of the selection function in the infinitesimal sexual model, in contrast with the asexual model. In the asexual model, we also investigate the effect of the mutation kernel and we show that kernels with higher kurtosis tend to reduce maladaptation and improve fitness, especially in fast changing environments.

**Highlights:**

- Adaptation to a changing environment may generate non Normal phenotypic distribution.
- The phenotypic variance at equilibrium truly depends on reproduction model;
- Selection shapes mean fitness only in sexual infinitesimal model;
- Weak selection away from the optimum leads to evolutionary tipping points with fast changes;
- Frequent mutations with large effects reduce maladaptation and improve fitness.

## 1. Introduction

Rapid environmental changes resulting from human activities have motivated the development of a theory to understand and predict the corresponding response of populations. Efforts have specially been focused on identifying conditions that allow populations to adapt and survive in changing environments (e.g. Pease et al., 1989; Lynch et al., 1991; Lynch and Lande, 1993; Burger and Lynch, 1995, for pioneering work). To this aim, most theoretical studies have modeled the evolution of polygenic quantitative traits subject to stabilizing selection around some optimal phenotype, whose value is shifted continuously through time (see Kopp and Matuszewski, 2014; Walters et al., 2012; Alexander et al., 2014). A major prediction of these early models is that when the optimal phenotype changes linearly with time, it will be tracked by the mean phenotype in the population with a lag that eventually stabilizes over time. This *evolutionary lag*, which quantifies the maladaptation induced by the environmental change, is predicted to depend on the rate of the change, on the phenotypic variance and on the strength of stabilizing selection on the trait. The maladaptation of the population due to the environmental change decreases the mean fitness of the population, which is commonly defined as the *lag load* or *evolutionary load* (Lynch and Lande, 1993; Lande and Shannon, 1996). Thus, above a critical rate of change of the optimal phenotype with time, the evolutionary lag is so large that the lag-load of the population will rise above the value that allows its persistence and the population will be doomed to extinction.

These predictions have typically been derived under the assumptions of (i) a particular form of selection, (ii) a constant genetic variance for the evolving trait, (iii) a Gaussian distribution of phenotypes and breeding values in the population. The selection function, describing how the Malthusian fitness declines away from the optimum, has typically a quadratic shape in many models (Bürger, 1999; Kopp and Matuszewski, 2014). However, the shape of selection functions is difficult to estimate and some studies suggest that it can strongly deviate from a quadratic shape, for example, in the case of phenological traits involved in climate adaptation (Gauzere et al., 2020). Although a quadratic shape may be an appropriate approximation for the fitness function close to its optimum, it may not be the case for strongly maladapted populations in a changing environment. Recently, (Osmond and Klausmeier, 2017; Klausmeier et al., 2020) have shown that “evolutionary tipping points” occur when the strength of selection weakens away from the optimum. In this situation, the population abruptly collapses when the speed of environmental change is too large. In this paper, we aim to investigate, in a general setting, the effects of the shape of selection functions on the adaptation of the population under environmental changes.

The genetic variance also plays a key role in the adaptation to changing environments and the determination of the critical rate of change. In many quantitative genetic models, this variance is assumed to be constant. Although it is approximately true on a short time scale, over a longer time scale the variance in the population is also subject to evolutionary change. More generally, obtaining mathematical predictions for the dynamics and the equilibrium value of the variance remains a notoriously difficult issue for many theoretical population genetics models (Barton and Turelli, 1989; Bürger, 2000; Barton and Keightley, 2002; Johnson and Barton, 2005; Hill, 2010). How the genetic variance evolves in a changing environment has therefore been explored mostly through simulations (Jones et al., 2012; Bürger, 1999; Waxman and Peck, 1999). In our paper, we overcome this problem by modeling the evolution of the entire phenotype distribution, This allows gaining some insights on the effect of maladaptation, induced by environmental changes, on the evolution of genetic variance.

Many theoretical works assumed that the phenotype distribution is Gaussian (Lynch et al., 1991). In the absence of environmental change, there are indeed many circumstances where the phenotypic distribution in the population is well captured by Gaussian distributions in quantitative genetics models. For example, in asexual populations, the distribution of a polygenic trait is Gaussian at mutation-selection equilibrium, providing that mutation effects are weak and selection is quadratic (Kimura, 1965; Lande, 1975; Fleming, 1979). In the case of sexual reproduction, similar outcomes are expected with the celebrated Fisher infinitesimal model of inheritance introduced by Fisher (1918). In this model, quantitative traits are under the control of many additive loci and each allele has a relatively small contribution on the character (Fisher, 1918). Within this framework, offspring are normally distributed within families around the mean of the two parental trait values, with fixed variance (Turelli and Barton, 1994; Turelli, 2017; Barton et al., 2017, and references therein). As a result, the phenotype distribution of the full population is well approximated by a Gaussian distribution under various assumptions on the selection function. Moreover, the usual Gaussian approximation of phenotypic trait distribution provides remarkably good approximation of the mean and the variance, even if disruptive selection generates strong deviations from normality (see Turelli and Barton 1994 under truncation selection, or see (Raoul, 2021) and (Calvez et al., 2019) for a wider class of selection functions). In the process of adaptation to environmental change, since the mean phenotype is lagging behind the optimum, selection however may induce a skew in the distribution (Jones et al., 2012). The distribution of the mutational effects can have a strong influence on the distribution as well, in particular when the evolutionary lag is large (Waxman and Peck, 1999). The Gaussian approximation of the phenotypic distribution should therefore naturally be questioned for both models of inheritance (asexual and infinitesimal sexual).

The main objective of this work is to derive signatures of maladaptation at equilibrium, *e.g*. the mean phenotype relative to the optimal phenotype, which allows us to quantify the evolutionary lag, the mean fitness and the phenotypic variance, depending on some general shape of selection and various features of trait inheritance. Those three components are linked by two generic identities describing the demographic equilibrium and the phenotypic equilibrium. Would the phenotypic variance be known, it would be possible to identify both the evolutionary lag and the mean fitness (Kopp and Matuszewski, 2014). In the general case, a third relationship is, however, needed. To this aim, we shall compute accurate approximations of the phenotypic distribution. Several methodological alternatives have been developed to unravel the phenotypic distribution, without any *a priori* on its shape. Previous methods attempted to derive the equations describing the dynamics of the mean, the variance and the higher moments of the distribution (Lande, 1975; Barton and Turelli, 1987; Turelli and Barton, 1990; Frank and Slatkin, 1990). Then, in his pioneering work, Burger (1991) derived relationships between the cumulants of the distribution, which are functions of the moments. However this system of equations is not closed, as the cumulants influence each other in cascade. More recently, Martin and Roques (2016) analyzed a large class of integro-differential models where the trait coincides with the fitness, through the partial differential equation (PDE) satisfied by the cumulant generating function (CGF). They applied their approach to the adaptation of asexual populations facing environmental change, using the Fisher Geometric Model for selection and specific assumptions on trait inheritance (diffusion approximation for the mutational effects) (Roques et al., 2020). However, the extension of their method to different models of selection or trait inheritance (general mutational kernel) seems difficult mainly because it relies on specific algebraic identities to reduce the complexity of the problem.

Here, we use deterministic quantitative genetics models based on integro differential equations to handle various shapes of stabilizing selection, and trait inheritance mechanisms. While we deal with a large class of thin-tailed mutational kernels in the asexual model, we restrict to the Fisher infinitesimal model as a mechanism of trait inheritance in sexually reproducing populations. We assume that the environment is changing linearly with time, as in the classical studies reviewed in (Kopp and Matuszewski, 2014). In order to provide quantitative results, we assume that little variance in fitness is generated at each reproduction event, through either mutation or recombination. It allows some flexibility about the trait inheritance process and the shape of the selection function. This assumption, here referred to as the *small variance regime*, enables using a mathematical framework developed in the past two decades in order to derive analytical features in models of quantitative genetics in asexual populations, mostly in a stationary phenotypic environment (Diekmann et al., 2005; Perthame and Barles, 2008; Barles et al., 2009; Lorz et al., 2011; Mirrahimi and Roquejoffre, 2016; Mirrahimi, 2017; Calvez and Lam, 2020), but see (Iglesias and Mirrahimi, 2021) in the case of a changing environment. This asymptotic methodology was first introduced by Diekmann et al. (2005) and Perthame (2007) in the context of evolutionary biology as an alternative formulation of adaptive dynamics, when the phenotypical changes are supposed to be small, but relatively frequent. Recently, this methodology has been also applied to the infinitesimal model for sexual reproduction in a stationary fitness landscape (Calvez et al., 2019; Patout, 2020). In the present paper, we apply this methodology to the case of a moving optimum.

From a mathematical perspective, the regime of small variance is analogous to some asymptotic analysis performed in mathematical physics, such as the approximation of geometrical optics for the wave equation at high frequency (Evans, 2010; Rauch, 2012), semi–classical analysis for the Schrödinger equation in quantum mechanics (Dimassi and Sjostrand, 1999; Zworski, 2012), and also the large deviation principle for stochastic processes (Fleming, 1977; Evans and Ishii, 1985; Freidlin and Wentzell, 1998; Feng and Kurtz, 2006). A common feature of these seemingly different asymptotic theories is to focus on the logarithm of the unknown function, and expand it with respect to a small parameter. We follow this route in the present work, by expanding the logarithm of the phenotypic density with respect to the relatively small variance.

Conversely to previous methods focusing on the moments of the phenotypic distribution, our approach focuses on the entire phenotypic distribution and it provides an accurate approximation of the phenotypic distribution even if it deviates significantly from the Gaussian shape. As a result, our method allows deriving analytical formulas for biologically relevant quantities, such as the relative mean phenotype and the evolutionary lag measuring maladaptation, the phenotypic variance within the population, the lag-load depressing the population mean fitness associated with critical rates of environmental changes, without solving the complete profile of the distribution. We are consequently able to answer the following questions

- What is the effect of the shape of selection on the adaptation of a population to a gradually changing environment?
- How does the distribution of mutational effects affect the adaptation dynamics?
- Does the choice of a particular reproduction model influence predictions about the dynamics of adaptation of a population?

## 2. Models and methodology

First, we describe in detail our general model of mutation-selection under changing environment with two different reproduction models (asexual and infinitesimal sexual) (Section 2.1). Then, we introduce the rescaled model including the relative variance parameter *ε*^2^ (Section 2.2) and we describe our methodology to investigate the regime of small variance (see Figure 1 for a sketch of the methodology). It is based on the asymptotic analysis with respect to this small parameter (Section 2.3). In Section 3, we provide, in the regime of small variance, analytical formula for the different characteristic quantities of the phenotypic distribution at equilibrium — mean fitness, mean relative phenotype and phenotypic variance — for the two different reproduction models: asexual model (Section 3.1) and infinitesimal sexual model (Section 3.2). After scaling back our results in the original units, we can compare the outcomes for the two systems of reproduction, and discuss the effect of a changing environment on the lag (Section 4.1), the mean fitness (Section 4.2) and the phenotypic variance (Section 4.3), respectively. Furthermore, we discuss the conditions for persistence of the population depending on the speed of the changing environment (Section 4.4) and we compare our approximation with numerical simulations of the whole distribution of the population (Section 4.5).

**Figure 1:**
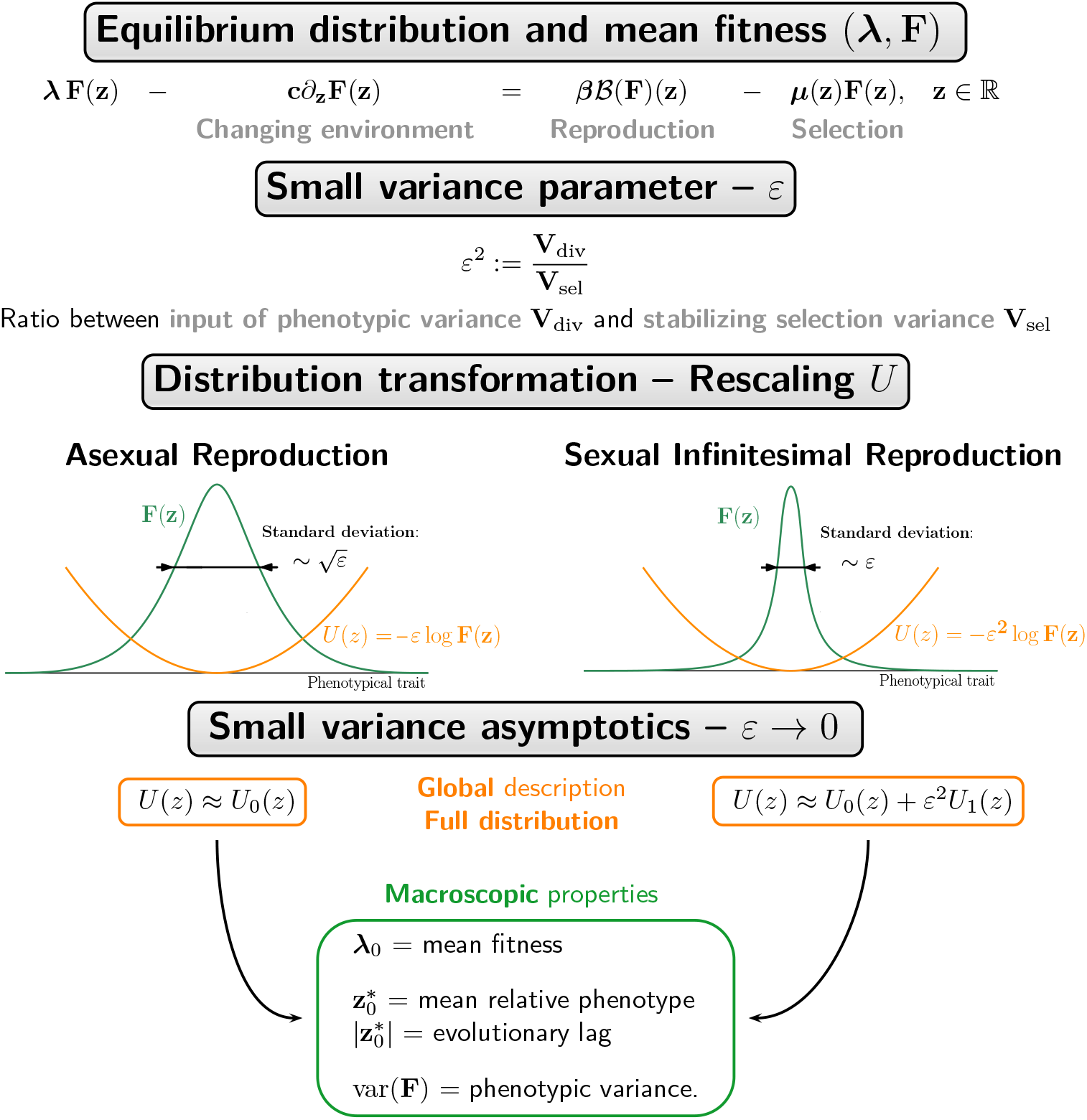
Schematic description of our methodology. To describe the equilibrium *F* we need the following steps: (1) Identify the scaling parameter *ε* and rescale the equation satisfied by the distribution *F* ; (2) Transform the distribution *F* into *U*. The transformed distribution *U* is the logarithmic of the density *F*, normalized by the ratio *ε* in the asexual reproduction case and by *ε*^2^ in the infinitesimal sexual reproduction case; (3) Identify the limit equation for *U* as *ε* → 0 (orange boxes) and deduce macroscopic properties (green box) such as the mean fitness *λ*_0_, the mean relative phenotype 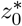 in the population, the evolutionary lag 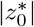 or the phenotypic variance at equilibrium Var(*F*).

### 2.1 The general model under changing environment

We consider a population reproducing in continuous time, subject to selection on the mortality rate, and to density-dependent competition. The population is structured by a one–dimensional phenotypic trait, denoted by **x** ∈ ℝ. The density of individuals with trait **x** is **f** (**t, x**) at time **t** > 0. For the sake of simplicity, the birth rate is assumed to be constant, set to value ***β*** > 0. Selection acts through the intrinsic mortality rate ***μ***(**t, x**), by means of stabilizing selection around some optimal value. In order to capture the dynamics of the population under a gradual environmental change, we assume that the optimal trait is shifted at a constant speed **c** > 0. We define the relative phenotype as the difference between the phenotypic value **x** and the optimal value at time **t**: **z** = **x** − **ct**. It quantifies the maladaptation of an individual of trait **x** in the changing environment. The intrinsic mortality rate ***μ*** is decomposed as follows

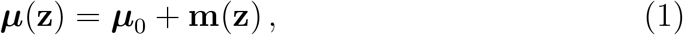

where ***μ***_0_ is the basal mortality rate at the optimum. The function **m**(**z**) = **m**(**x** − **ct**) is the increment of mortality due to maladaptation and its shape encodes the effect of selection. We assume that **m** ⩾ 0 attains its unique minimum value at 0 where **m**(0) = 0, and it is symmetrically increasing: **m** is decreasing on (−∞, 0) and increasing on (0, ∞). We further assume that ***β*** > ***μ***_0_, which ensures a net growth of individuals at the optimal trait.

The dynamics of the density **f** (**t, x**) is given by the following equation:

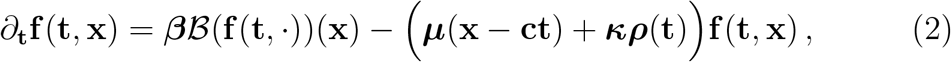

where the term ***ρ***(**t**) = ∫_ℝ_ **f** (**t, x**′)*d***x**′ corresponds to the size of the population, and ***κ*** > 0 is the strength of competition within the population. This nonlinear term introduces density–dependent mortality in the model, as it reduces the population growth rate at high density. Integrating the model (2) over the **x** variable, the population size ***ρ***(**t**) satisfies the following logistic equation

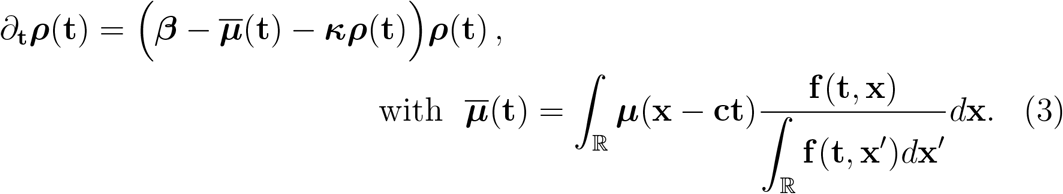

The operator ℬ describes how new individuals with phenotype **x** are generated depending on the whole phenotypic density. For simplicity, we assume no environmental effects on the expression of the phenotype, and phenotypic values equal to breeding values. We consider the two following choices for the reproduction operator ℬ.

#### Asexual model of reproduction with mutations

We first consider the case of asexual reproduction where the phenotype of an offspring **x** is drawn randomly around the phenotype of its single parent **x**′. We restrict to the case where the changes depend only on the trait difference **x**′−**x**, described by the kernel **K**_div_. The reproduction operator has then the following expression:

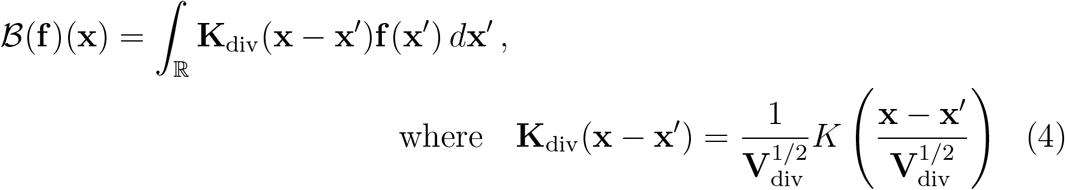

where *K* is a symmetrical probability density function with unit variance. Hence, **V**_div_ is the variance of the phenotypic changes at each reproduction event. We assume that *K* decays faster than some exponential function. This is usually called a *thin–tailed kernel*. This corresponds to the scenario where the mutations with large effect on phenotypic traits are rare.

The extremal case corresponding to accumulation of infinitesimal changes is referred to as the *diffusion approximation*. This translates into the following formula

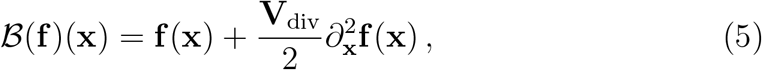

In this case, the shape of the kernel does not matter and only the variance remains.

The general form (4) encompasses the decomposition of the kernel **K**_div_ into **K**_div_ = (1 − *η*)*δ*_0_ + *η***K**_mut_, where *η* ∈ [0, 1] is the probability of a mutation, *δ*_0_ is the Dirac mass at 0 and **K**_mut_ is the probability distribution of mutational effects. In such case, **V**_div_ = *η***V**_mut_, where **V**_mut_ is the variance associated with the mutational effects.

#### Infinitesimal model of sexual reproduction

Secondly, we consider the case where the phenotype of the offspring **x** is drawn randomly around the mean trait of its parents (**x**_1_, **x**_2_), following a Gaussian distribution *G*_LE_. This is known as the Fisher infinitesimal model (Fisher, 1918; Bulmer, 1980; Turelli and Barton, 1994; Tufto, 2000; Barton et al., 2017). The reproduction operator has then the following expression:

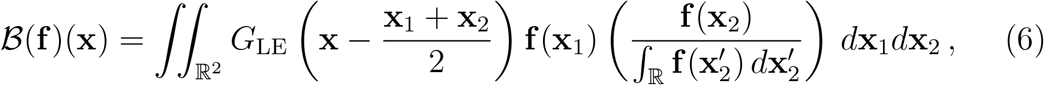

where *G*_LE_ denotes the centered Gaussian distribution with variance **V**_LE_/2. Here the parameter **V**_LE_ corresponds to the genetic variance at linkage equilibrium in the absence of selection (Bulmer, 1971; Lange, 1978; Bulmer, 1974; Santiago, 1998; Turelli and Barton, 1994). For comparison with the asexual case we shall use the same notation **V**_div_ for **V**_LE_, as it scales the input of phenotypic variance in the population for each reproduction event (see discussion below).

#### Input of phenotypic variance through reproduction

In the absence of selection (**m**(**z**) = 0) and random drift, the input of phenotypic variance per reproduction event is scaled in both model by **V**_div_. Indeed, we can show using equation (2) that, in this situation, the dynamics of phenotypic variance are

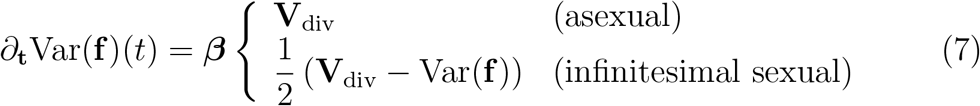

where the variance Var(**f**) and the mean trait 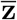 are defined by

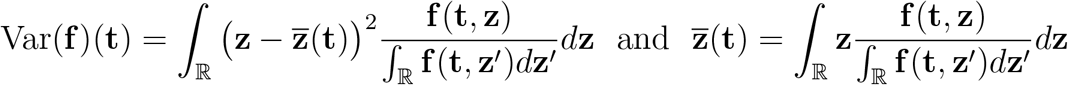

Even if the variance in the offspring distribution in the infinitesimal sexual model and in the asexual model are conceptually different, they both scale with the input of phenotypic variance per reproduction event, that is why we use the same notation **V**_div_.

However, the impact of diversity depends on the model of reproduction. In the asexual case, the variance of the phenotypic distribution increases indefinitely in the absence of selection (7). Thus, the asexual model does not impose any strong constraint on the variance of the phenotypic distribution of the population. Conversely, in the absence of selection, the infinitesimal sexual model generates a finite phenotypic variance at equilibrium, equal to **V**_div_. Thus the dynamics of the phenotypic variance are more constrained in the infinitesimal model than in the asexual model.

#### Equilibrium in a changing environment

In this paper, we focus on the asymptotic behavior of the model, studying whether the population will persist or go extinct in the long term. In order to mathematically address the problem, we seek special solutions of the form **f** (**t, x**) = **F**(**x** − **ct**). These solutions correspond to a situation where the phenotypic distribution **F** has reached an equilibrium, which is shifted at the same speed **c** as the environmental change. This distribution of relative phenotype **z** := **x** − **ct** quantifies *maladaptation* within the population. One can observe from equation (2) that the trivial solution, which corresponds to **F** = 0, always exists. Our aim is first to decipher when non trivial equilibrium **F** exists. Secondly, we characterize in detail the distribution **F** when it exists.

Using the property of invariance by translation verified by the reproduction operator ℬ, we obtain that a non trivial, non-negative, equilibrium **F** solves the following eigenvalue problem,

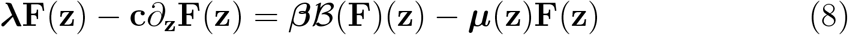

where the eigenvalue ***λ*** is expected to be positive ***λ*** > 0, since it must satisfy

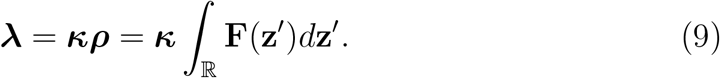

The transport term −**c***∂*_**z**_**F** corresponds to the effect of the moving optimum on the phenotypic distribution **F** at equilibrium. Moreover, since **F** is the density at equilibrium, the population size ***ρ*** is an equilibrium of (3), which provides the following relationship

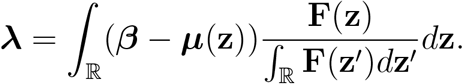

The eigenvalue ***λ*** can thus be interpreted as a measure of the *mean fitness* of the population, or its mean intrinsic rate of increase, where ***β*** − ***μ***(**z**) is the contribution of an individual with relative phenotype **z** to the growth rate of the population at low density. Thus, an analytical description of ***λ*** will provide a formula for the critical speed of environmental change above which extinction is predicted, corresponding to the case where the eigenvalue ***λ*** is negative. The value ***λ*** also informs us on the size of the population at equilibrium in presence of a changing environment, ***ρ*** (see equation (9)).

Our aim is to describe accurately the couple (***λ*, F**) in presence of a moving optimum with constant speed **c** in both reproduction scenarios. To do so, we compute formal asymptotics of (***λ*, F**) at a weak selection or slow evolution limit when little variance in fitness is generated by mutation or sexual reproduction per generation. Note that the shape of **F** is not prescribed *a priori* and the methodology presented here can handle significantly large deviations from Gaussian distributions.

Noteworthy, the equation (8) with asexual reproduction operators defined by (4) or (5) admits solutions under suitable conditions. Cloez and Gabriel (2020) proved that solutions exist for any speed **c** if the mortality function ***μ*** goes to ∞ when |**z**| → ∞. Furthermore, Coville and Hamel (2019) proved that solutions also exist for more general mortality functions ***μ*** as soon as the speed **c** remains below a critical threshold. For the infinitesimal operator (6), Calvez et al. (2019) proved the existence of solutions without changing environment and in the special regime of small variance described below. The existence of a pair (***λ*, F**) for positive speed **c** will be the topic of a future mathematical paper.

### 2.2 Adimensionalization

In order to compute asymptotics of the solution of our model, we first need to rescale the model with dimensionless parameters (see Table 1 for the relationship between original variables and their values after rescaling and Supplementary Information SI B for mathematical details).

**Table 1:**
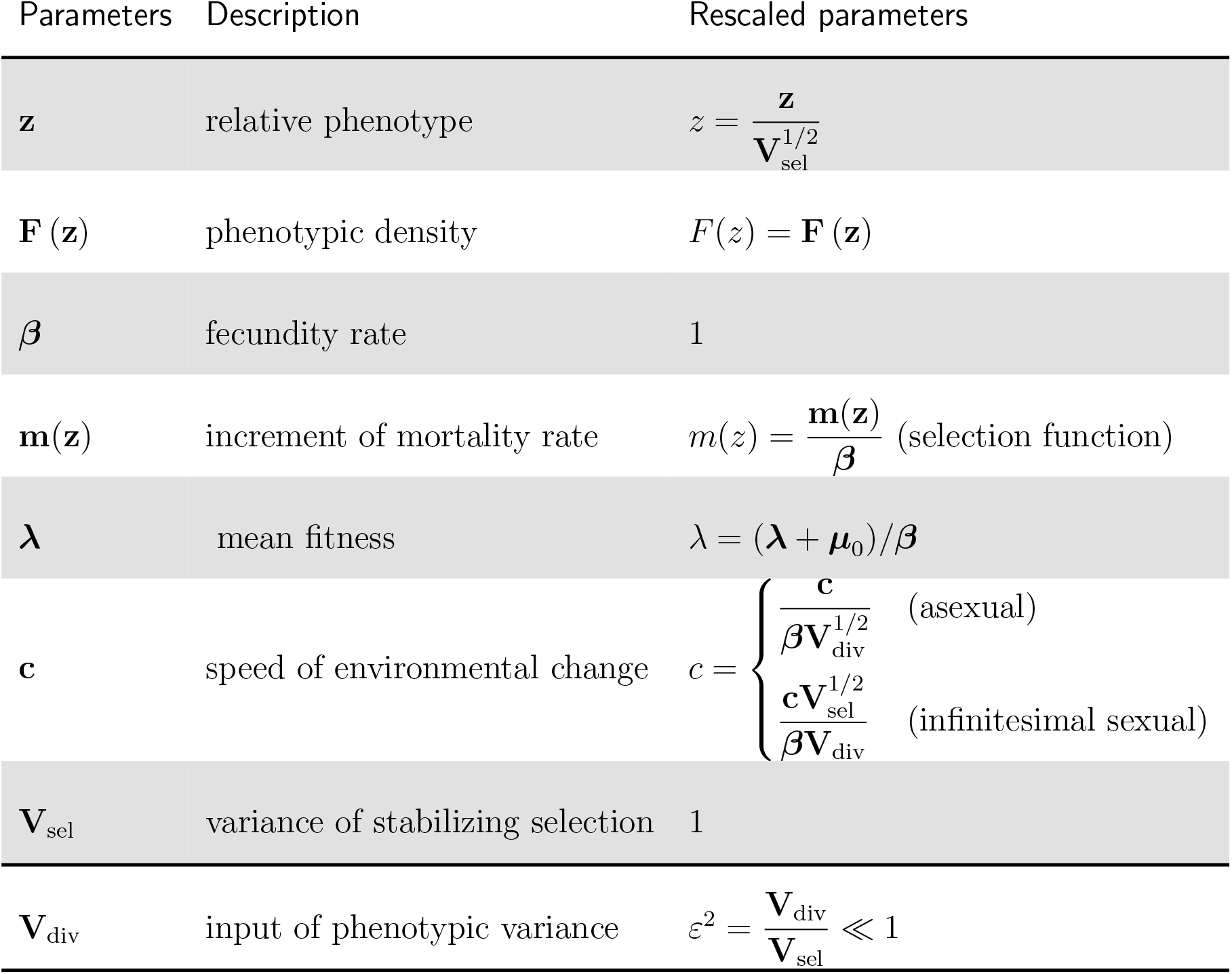
Biological parameters and their formula after rescaling for both the asexual and infinitesimal sexual model. Our methodology relies on the assumption that the dimensionless parameter *ε* is small, *ε* ≪ 1, while the other rescaled parameters are of order 1.

| Parameters | Description | Rescaled parameters |
| --- | --- | --- |
| $\mathbf{z}$ | relative phenotype | $z = \frac{\mathbf{z}}{\mathbf{V}_{\text{sel}}^{1/2}}$ |
| $\mathbf{F}(\mathbf{z})$ | phenotypic density | $F(z) = \mathbf{F}(\mathbf{z})$ |
| $\beta$ | fecundity rate | 1 |
| $\mathbf{m}(\mathbf{z})$ | increment of mortality rate | $m(z) = \frac{\mathbf{m}(\mathbf{z})}{\beta}$ (selection function) |
| $\lambda$ | mean fitness | $\lambda = (\lambda + \mu_0)/\beta$ |
| $\mathbf{c}$ | speed of environmental change | $c = \begin{cases} \frac{\mathbf{c}}{\beta \mathbf{V}_{\text{div}}^{1/2}} & (\text{asexual}) \\ \frac{\mathbf{c} \mathbf{V}_{\text{sel}}^{1/2}}{\beta \mathbf{V}_{\text{div}}} & (\text{infinitesimal sexual}) \end{cases}$ |
| $\mathbf{V}_{\text{sel}}$ | variance of stabilizing selection | 1 |
| $\mathbf{V}_{\text{div}}$ | input of phenotypic variance | $\varepsilon^2 = \frac{\mathbf{V}_{\text{div}}}{\mathbf{V}_{\text{sel}}} \ll 1$ |

#### Time scale

We introduce the relative time coordinate *t* = ***β*t**, to scale the model according to the generation time. Hence, the dimensionless fecundity rate equals one, and the increment of mortality is *m* = **m**/***β***, which corresponds to the *selection function*. The effect of stabilizing selection is captured by **V**_sel_, which is inversely proportional to the strength of selection around the optimum:

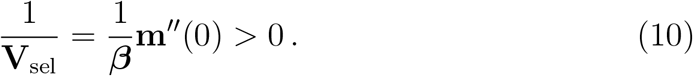

Note that **V**_sel_ scales as a variance parameter.

#### Phenotypic bscale

All measures depending on phenotypic units are expressed in unit 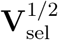, and we change variables accordingly, 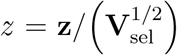. As such, the strength of selection in the rescaled system is equal to unity:

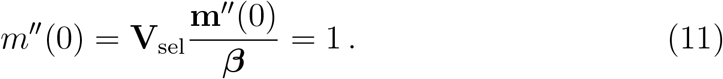

#### Phenotypic variance parameter

Similarly, in both asexual and infinitesimal sexual models, the dimensionless parameter describing how much phenotypic variance is introduced in the population at each generation is 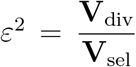. Accordingly, we have the following expression in dimensionless variables,

##### Asexual reproduction

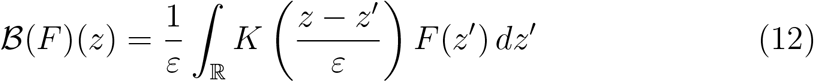

##### Infinitesimal sexual reproduction

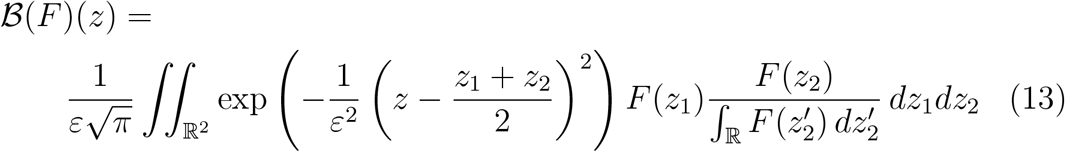

#### Speed of environmental change

The rescaling of the speed of environmental change differs in the asexual and infinitesimal sexual versions of our model. In both models, the ability to evolve fast enough to track the moving optimum depends critically on the input of phenotypic variation fueling evolutionary change. Since this input is of a different nature between the models, we have to adjust the scale of the speed differently in each context to observe non-trivial behaviours. We define accordingly the speed of change 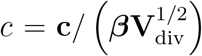 in the asexual case, but 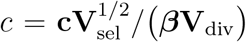 in the case of the infinitesimal model (see Table 1).

As a consequence, the transport term −**c***∂*_**z**_**F** which carries the effect of environmental change in (8), inherits respectively a factor *ε* (asexual) and *ε*^2^ (infinitesimal), see SI SI B.3. A mismatch in this expression (e.g. involving any other power of *ε*) would result in a severe unbalance between the various contributions in the models, leading either to a dramatic collapse of the population if the effective speed is too large, or to no significant effect of the change if the effective speed is too small.

In addition, the discrepancy between these scaling formula reveals a strong difference on the effect of the selection between the models. Indeed, the strength of selection is involved in the infinitesimal sexual model, whereas it does not appear in the asexual model. Our analysis is aimed to enlighten and explain those differences (see section 4).

#### Rescaled model

Using these rescaled variables, we obtain the following equations:

##### Asexual reproduction

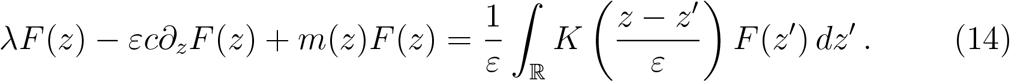

##### Infinitesimal sexual reproduction

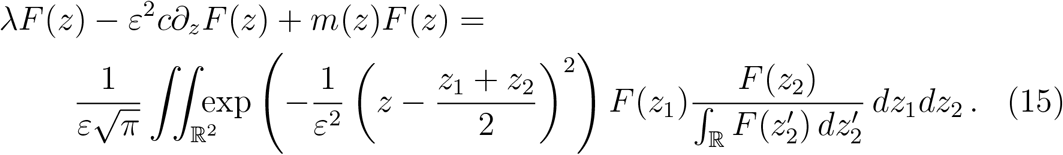

### 2.3. Small variance asymptotics

In the following, we further assume that the parameter *ε* is small, which means that little variance in fitness is introduced in the population through either mutation or recombination during reproduction. This is what we call *the small variance regime*. This situation may happen either when the input of phenotypic variation is small or because stabilizing selection is weak.

In the asexual model, we also assume that mutations are reasonably frequent, that is the probability of mutation *η* is of order one. Under the small variance regime (*ε* ≪ 1), this assumption of frequent mutations prevents the mutation kernel *K* to degenerate in our scaling regime, which is a key assumption in the mathematical framework introduced by Diekmann et al. (2005) (see SI D.4 for mathematical details).

Moreover, our regime of small variance (*ε* ≪ 1 ∼ *η*) is usually referred to as the strong mutation and weak selection regime, which is linked to the Gaussian approximation regime when **V**_mut_ ≪ *η***V**_sel_ (Kimura, 1965; Lande, 1975; Fleming, 1979; Bürger, 2000). This regime of frequent mutations contrasts with the House-of-Cards (HC) regime where mutations are rare with large effects when *η***V**_sel_ ≪ **V**_mut_ (Turelli, 1984; Turelli and Barton, 1990; Bürger, 2000). In the HC regime, the mutation rate *η* is smaller than our *ε* parameter (*η* ≪ *ε* ≪ 1). Our analysis would thus fail in this regime, because the asymptotic limits are conceptually different.

In the small variance regime (*ε* ≪ 1), we expect the equilibrium *F* to be concentrated around a mean value *z*\* of the relative phenotype, that we name the *mean relative phenotype*, see Fig. 1. The evolutionary lag |*z*\*| is defined here as the distance between the mean phenotypic trait in the population and the optimal trait. Note that, in previous literature the evolutionary lag is sometimes defined as we do here (e.g. Gomulkiewicz and Houle, 2009), sometimes as the difference between the mean trait value and the optimum, referred as the mean relative phenotype here (e.g. Burger and Lynch, 1995) or the opposite difference (e.g. Lande and Shannon, 1996).

The core of our approach consists in the accurate description of the phenotypic distribution *F* when *ε* ≪ 1. This is made possible after a suitable transformation of the phenotypic distribution *F*. The Cole-Hopf transformation is an appropriate mathematical tool to provide approximations of singular distributions with respect to a small parameter, for instance the wavelength in wave propagation (geometric optics) or the Planck constant in quantum mechanics (semi–classical analysis), and also the phenotypic variance in our theoretical biology setting. It is defined as the logarithm of the density *F*, multiplied by a small parameter related to the order of magnitude of the phenotypic variance. In our problem, we need to introduce different quantities depending on the modeling choice:

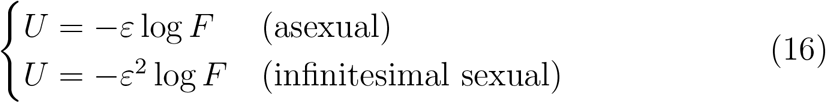

We emphasize that the discrepancy between the two scenarios is an outcome of our analysis. The scaling has been carefully tuned to induce a non trivial limit in the regime *ε* ≪ 1. We discuss this scaling in details in the Discussion section. In order to describe *U* asymptotically, we expand it with respect to *ε* as follows:

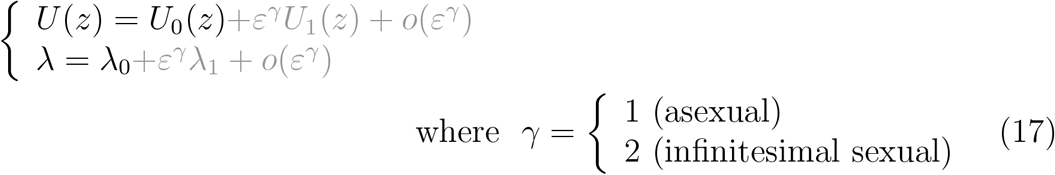

and (*λ*_0_, *U*_0_) is the limit shape as *ε* → 0, and (*λ*_1_, *U*_1_) is the correction for small *ε* > 0. In the next sections 3.1 and 3.2, we show, by formal arguments, that the function *U* and the mean fitness *λ* converge towards some non trivial function *U*_0_ and some value *λ*_0_ as *ε* → 0.

In the following section Results, we compute relevant quantitative features, such as the mean fitness *λ*_0_, the mean relative phenotype 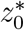, and the phenotypic variance Var(*F*). The latter is related to *U*_0_ by the following formula (derived in SI C):

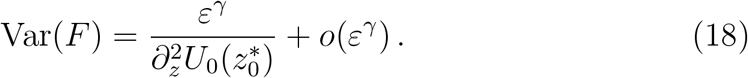

Remarkably, our methodology is able to compute those quantities directly, bypassing the resolution of the limit equation solved by (*λ*_0_, *U*_0_) (which may have non-explicit solutions).

## 3. Results in the regime of small variance

### 3.1. The asexual model

Using the logarithmic transformation (16) to reformulate our problem (14) and the Taylor expansion of the pair (*λ, U*) with *γ* = 1, we show that the limit shape (*λ*_0_, *U*_0_) satisfies the following problem (see SI D.1):

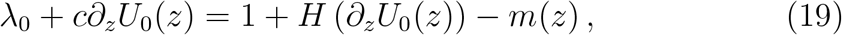

where the Hamiltonian function *H* is the two-sided Laplace transform of the mutation kernel *K* up to a unit constant:

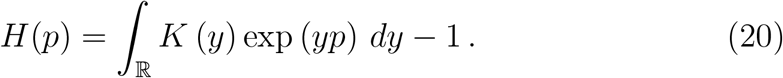

It is a convex function that satisfies *H*(0) = *H*′(0) = 0, and *H*′′(0) = 1 from hypothesis (4) on the mutation kernel *K*. Moreover, thanks to our assumption on the mutation probability *η*, the function *H* is not singular (see SI D.4 for more details).

We can remark that the shape of the equation (19) also contains *the diffusion approximation model* where the reproduction operator is approximated by a diffusion operator (5). For the diffusion approximation, we find that the Hamiltonian function is given by *H*(*p*) = *p*^2^/2 (see SI D.1.1)

#### Computation of the mean fitness λ_0_

We find that (see SI D.1.3 for details)

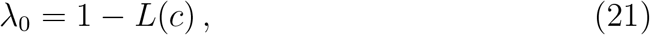

where the Lagrangian function *L*, also known as the Legendre transform of the Hamiltonian function *H*, is defined as:

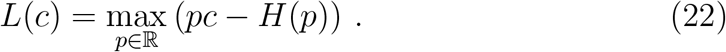

It is a convex function satisfying *L*(0) = *L*′(0) = 0, and *L*′′(0) = 1. Moreover, we always have *L*(*c*) ⩽ |*c*|^2^/2 where *L*(*c*) = |*c*|^2^/2 corresponds to the *diffusion approximation* case.

Since the mean fitness is *λ*_0_ = 1 in the absence of environmental change, the quantity *L*(*c*) represents the *lag-load* in the rescaled units, which is induced by the moving optimum (Lynch and Lande, 1993; Lande and Shannon, 1996). Moreover, if we push the expansion to the higher order we are able to compute the following mean fitness (see SI D.6 for mathematical details)

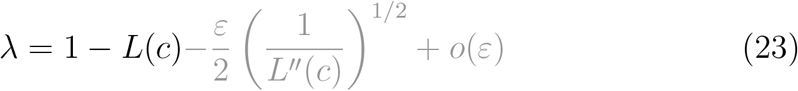

The new term of order *ε* can be seen as the *standing load*, i.e. a reduction in mean fitness due to segregating variance for the trait in the population (Lynch and Lande, 1993; Burger and Lynch, 1995; Kopp and Matuszewski, 2014).

#### Computation of the mean relative phenotype 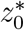

We obtain from the main equation (19), evaluated at 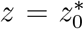, that 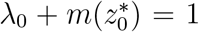. Thus, combining with equation (21), we deduce that 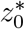 is a root of

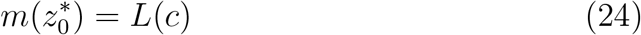

with the appropriate sign, that is *m*′(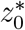) and *c* have opposite signs: 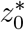 < 0 if *c* > 0 and vice-versa.

#### Computation of the phenotypic variance

From equation (18), we need to compute the second derivative of *U*_0_ at the mean relative phenotype 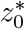. We can derive it from the differentiation of equation (19) evaluated at 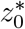 (recall that *H*′(0) = 0 by symmetry of the mutation kernel *K*):

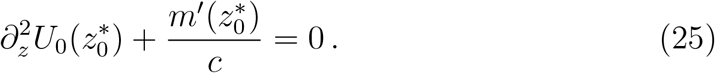

We deduce the following first order approximation of the phenotypic variance:

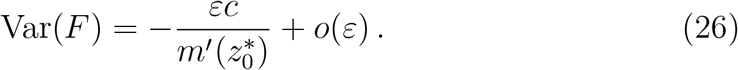

##### Remark 1

*The expressions obtained in this section are still valid when c* = 0. *A direct evaluation gives that λ*_0_ = 1 *and* 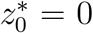. *Moreover, we show in Supplementary Information SI D.5 that in the limit c* → 0, *the previous formula* (25) *becomes*

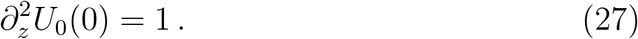

We will discuss the biological implications of these predictions after expressing them in the original units in the section 4.

### 3.2. The infinitesimal model of sexual reproduction in the regime of small variance

#### The limiting problem formulation

Remarkably enough, a similar mathematical analysis can be performed when the convolution operator (12) is replaced with the infinitesimal model for reproduction (13). However, the calculations are slightly more involved than the former case, but the final result is somewhat simpler. Here, the suitable logarithmic transformation of the phenotypic distribution *F* is *U* = −*ε*^2^ log(*F*). The equation for the new unknown function *U* is:

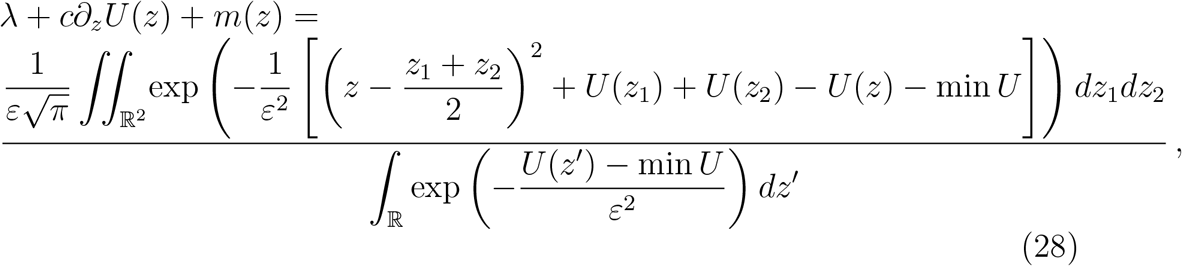

where min *U* has been subtracted both in the numerator and the denominator. The specific form of the right-hand-side characterizes the shape of *U*. Indeed, the quantity between brackets must remain non negative, unless the integral takes arbitrarily large values as *ε* → 0. Moreover, its minimum value over (*z*_1_, *z*_2_) ∈ ℝ^2^ must be zero, unless the integral vanishes. As a consequence, the function *U* must be a quadratic function of the form 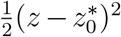 where the mean relative phenotype of the distribution, 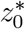, can be determined aside (see SI F.1 for details). To describe 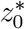, we expand the pair (*λ, U*), in a power series with respect to *ε*^2^:

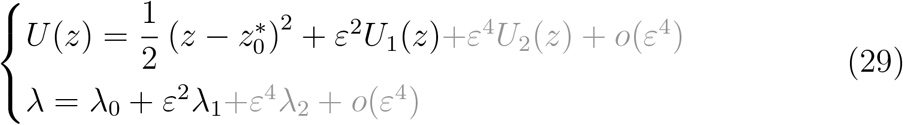

Plugging this expansion into (28), we obtain the following equation on the corrector *U*_1_:

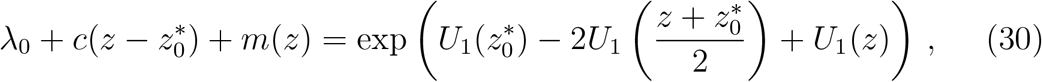

which contains as a by–product the value of some quantities of interest, such as the mean fitness *λ*_0_, and the mean relative phenotype 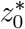. Moreover, we can solve this equation if, and only if *λ*_0_ and 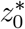 take specific values that we identify below.

The mid-point 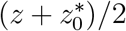 which appears in the right-hand-side of (30) has a direct interpretation in terms of the conditional distribution of parental traits. It means that an individual of trait *z* is very likely to be issued from a pair of parents having both traits close to the mid-value between *z* and the mean phenotype 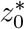 (and equal to 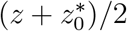 in the limit *ε* → 0). This is the result of the following trade-off: on the one hand parents with traits close to the mean trait value 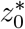 are frequent but the chance of producing offspring with relative phenotype 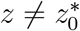 is too small; on the other hand, parents with traits evenly distributed around *z* would likely produce offspring with relative phenotype *z*, but they are not frequent enough. As a compromise, the most likely configuration is when both parents have their relative traits close to 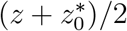, see Figure S2 and SI F.2.1.

#### Computation of macroscopic quantities

Let us first observe that equation (30) is equivalent to the following one:

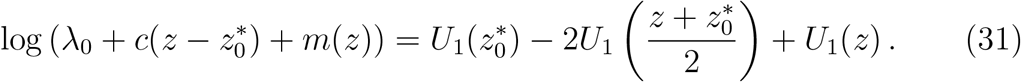

The key observation is that the expression on the right hand side vanishes at 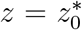, and so does its first derivative with respect to *z* at 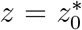. This provides two equations for the two unknowns *λ*_0_, 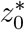, without computing the exact form of *U*_1_:

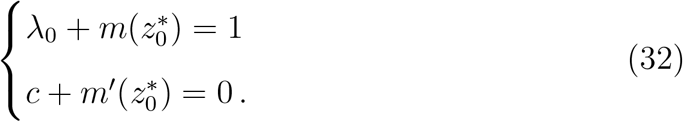

These two relationships are necessary and sufficient conditions, meaning that they guarantee that equation (30) admits at least one solution *U*_1_ (see SI F.1 for mathematical details, and (Calvez et al., 2019)). In addition, we can push the expansion further and we can gain access to the higher order of approximation for the quantities of interest (see SI F.2).

##### Mean relative phenotype

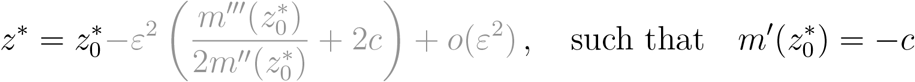

##### Mean fitness

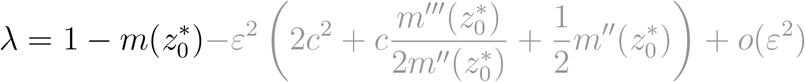

##### Phenotypic variance

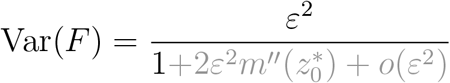

## 4. Comparison of predictions of the asexual and infinitesimal models

To discuss our mathematical results from a biological perspective, we need to scale back the results in the original units (see Table 1 for the link between the scaled parameters and the parameters in the original units). Our general predictions for macroscopic quantities in the original units are shown in Table 2. For ease of comparison with previous literature, which has generally assumed a quadratic form for the selection function, we present our predictions in Table 3 under this special assumption and with the diffusion approximation.

**Table 2:**
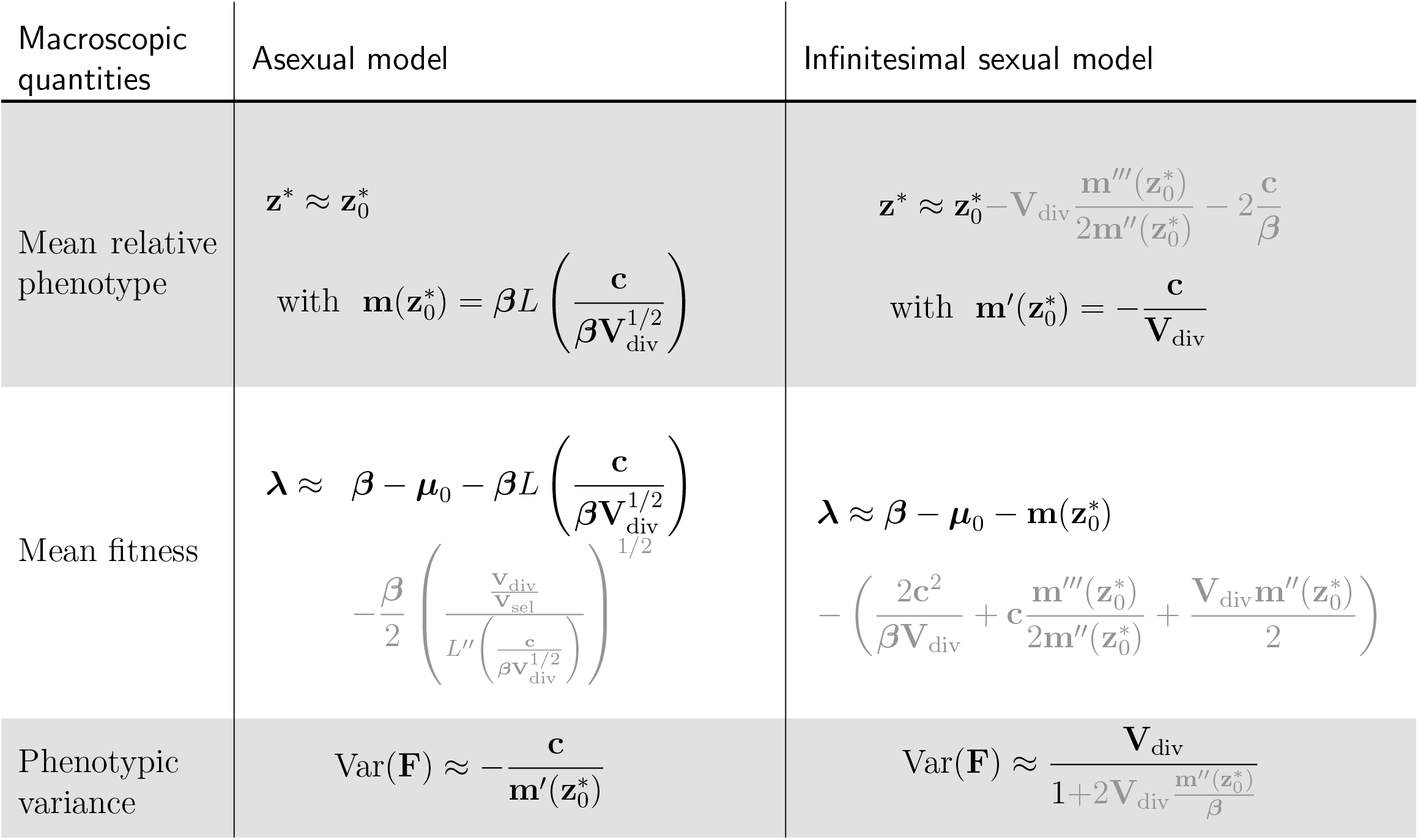
Analytical predictions for the mean relative phenotype **z**\*, the mean fitness ***λ*** and the phenotypic variance Var(**F**) for both the asexual and infinitesimal sexual model in the original variables. Black terms correspond to first order approximations, while gray terms correspond to second order approximations. In the asexual model, *L* is the Lagrangian defined by (22) and it is associated to the mutation kernel *K* by a two-sided Laplace transformation.

**Table 3:** Analytical predictions for the mean relative phenotype **z**\*, the mean fitness ***λ*** and the phenotypic variance Var(**F**) for both the asexual and infinitesimal sexual model in the original variable when assuming a quadratic form of selection *m*(*z*) = *z*^2^/2 (corresponding to **m**(**z**)/***β*** = **z**^2^/(2**V**_sel_) in original units). In the asexual model, we are under the diffusion approximation: *L*(*v*) = *v*^2^/2. Black terms correspond to first order approximations, while gray terms correspond to second order approximations.

| Macroscopic quantities | Asexual model<br>(quadratic selection / diffusive approx) | Infinitesimal sexual model<br>(quadratic selection) |
| --- | --- | --- |
| Mean relative phenotype | $\mathbf{z}^* = -\frac{\mathbf{c}}{\beta} \left( \frac{\mathbf{V}_{\text{sel}}}{\mathbf{V}_{\text{div}}} \right)^{1/2}$ | $\mathbf{z}^* \approx -\frac{\mathbf{c}}{\beta} \left( \frac{\mathbf{V}_{\text{sel}}}{\mathbf{V}_{\text{div}}} \right) - 2\frac{\mathbf{c}}{\beta}$ |
| Mean fitness | $\lambda = \beta - \mu_0 - \frac{\mathbf{c}^2}{2\beta\mathbf{V}_{\text{div}}} - \frac{\beta}{2} \left( \frac{\mathbf{V}_{\text{div}}}{\mathbf{V}_{\text{sel}}} \right)^{1/2}$ | $\lambda \approx \beta - \mu_0 - \frac{\mathbf{c}^2\mathbf{V}_{\text{sel}}}{2\beta\mathbf{V}_{\text{div}}^2} - \left( \frac{2\mathbf{c}^2}{\beta\mathbf{V}_{\text{div}}} + \frac{\beta\mathbf{V}_{\text{div}}}{2\mathbf{V}_{\text{sel}}} \right)$ |
| Phenotypic variance | $\text{Var}(\mathbf{F}) = (\mathbf{V}_{\text{div}}\mathbf{V}_{\text{sel}})^{1/2}$ | $\text{Var}(\mathbf{F}) \approx \frac{\mathbf{V}_{\text{div}}}{1 + 2\frac{\mathbf{V}_{\text{div}}}{\mathbf{V}_{\text{sel}}}}$ |

### Numerical simulations

To illustrate our discussion, we also perform numerical simulations. The simulated stationary distribution is obtained through long time simulations of a suitable numerical scheme for (2) (details in SI SI G). Using this numerical expression, we compute the lag, the mean fitness and the phenotypic variance of the distribution. In the asexual model, the function *U*_0_ is obtained from the direct resolution of the ordinary differential equation (19) using classical integration methods – see SI D.7. In the infinitesimal model, the correction *U*_1_ is computed directly from its analytical expression given in SI F.2.4. The macroscopic quantities in the regime of small variance are directly computed from their analytical expressions given in the Table 2 and 3. We also compare our analytical expression with the outcome of a stochastic model, which considers an evolving population with a finite number of individuals (see SI H).

### 4.1. Mean relative phenotype and evolutionary lag

The mean relative phenotype **z**\* is here defined as the difference between the mean phenotypic trait in the population **x**\* and the optimal trait **ct**. In our study, for numerical illustration, we used **c** > 0. So a negative mean relative phenotype **z**\* indicates that the distribution of phenotype lags behind the optimal trait. The maladaptation of the population is generally measured by the *evolutionary lag*, which is defined as the distance between the mean phenotypic trait and the optimal trait. In our study, the evolutionary lag corresponds to the absolute value of the mean relative phenotype, |**z**\*| = |**x**\* − **ct**|, which is positive.

#### The lag increases with the speed of environmental change

In both the asexual model and infinitesimal model, we recover the classic result that the lag 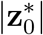 is an increasing function of **c** (as illustrated by Fig. 2 and Fig. 3).

**Figure 2:**
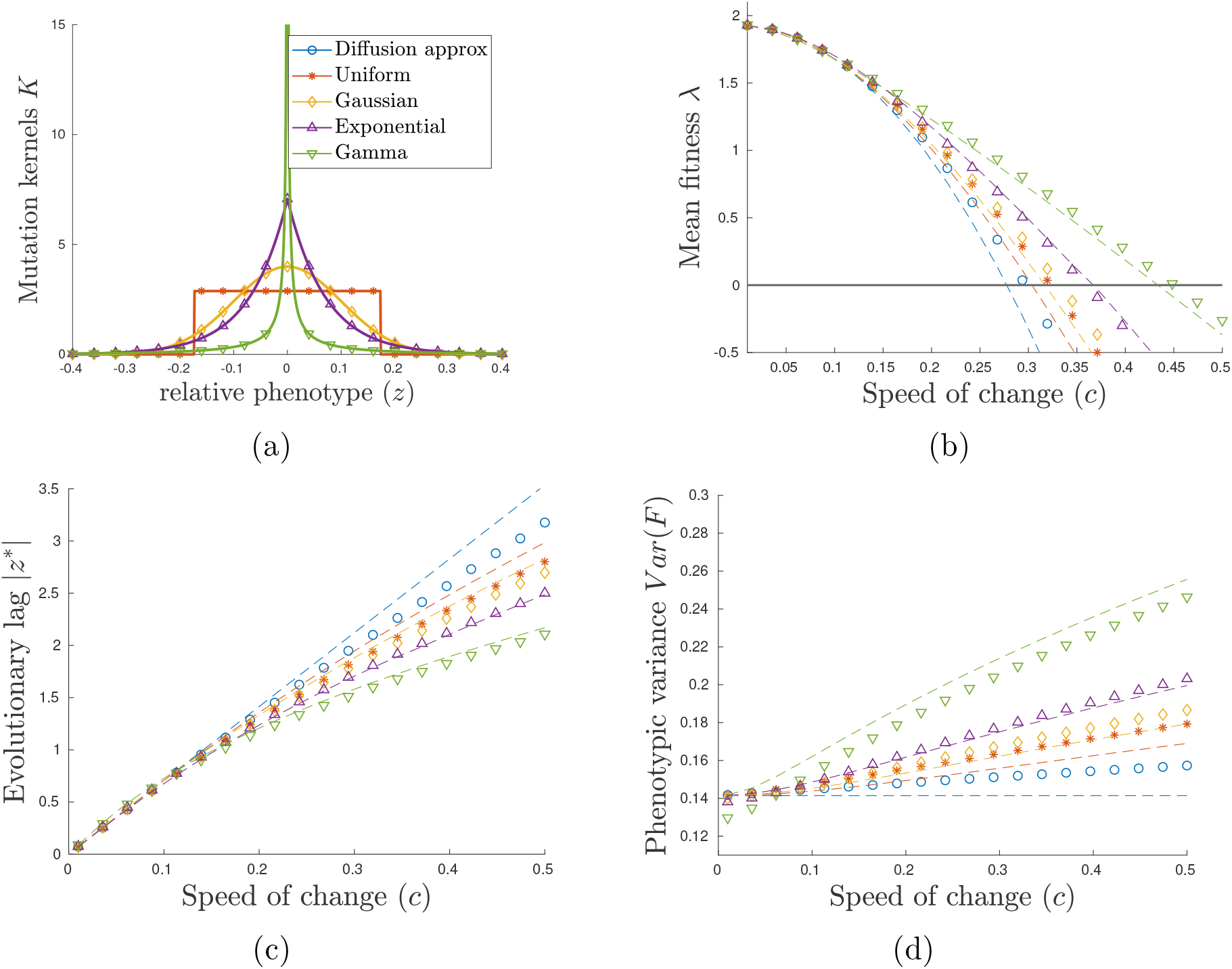
Influence of the mutational kernel *K*, described in panel (a), on (b) the mean fitness ***λ***, (c) the evolutionary lag |**z**\*| and (d) the phenotypic variance Var(**F**) at equilibrium in an environment changing at rate **c** ranging in (0, 0.5) for the asexual model with quadratic selection function *m*(*z*) = *z*^2^/2. We compare the diffusion approximation (blue curves) with four different mutation kernels with the same variance **V**_div_ = 0.01, while the variance of selection is **V**_sel_ = 1: the Uniform distribution (red curves), the Gaussian distribution (orange curves), Exponential distribution (purple curves) and Gamma distribution (green curves). For each case we compare our analytical results (dashed lines) with the simulation results (marked symbol).

**Figure 3:**
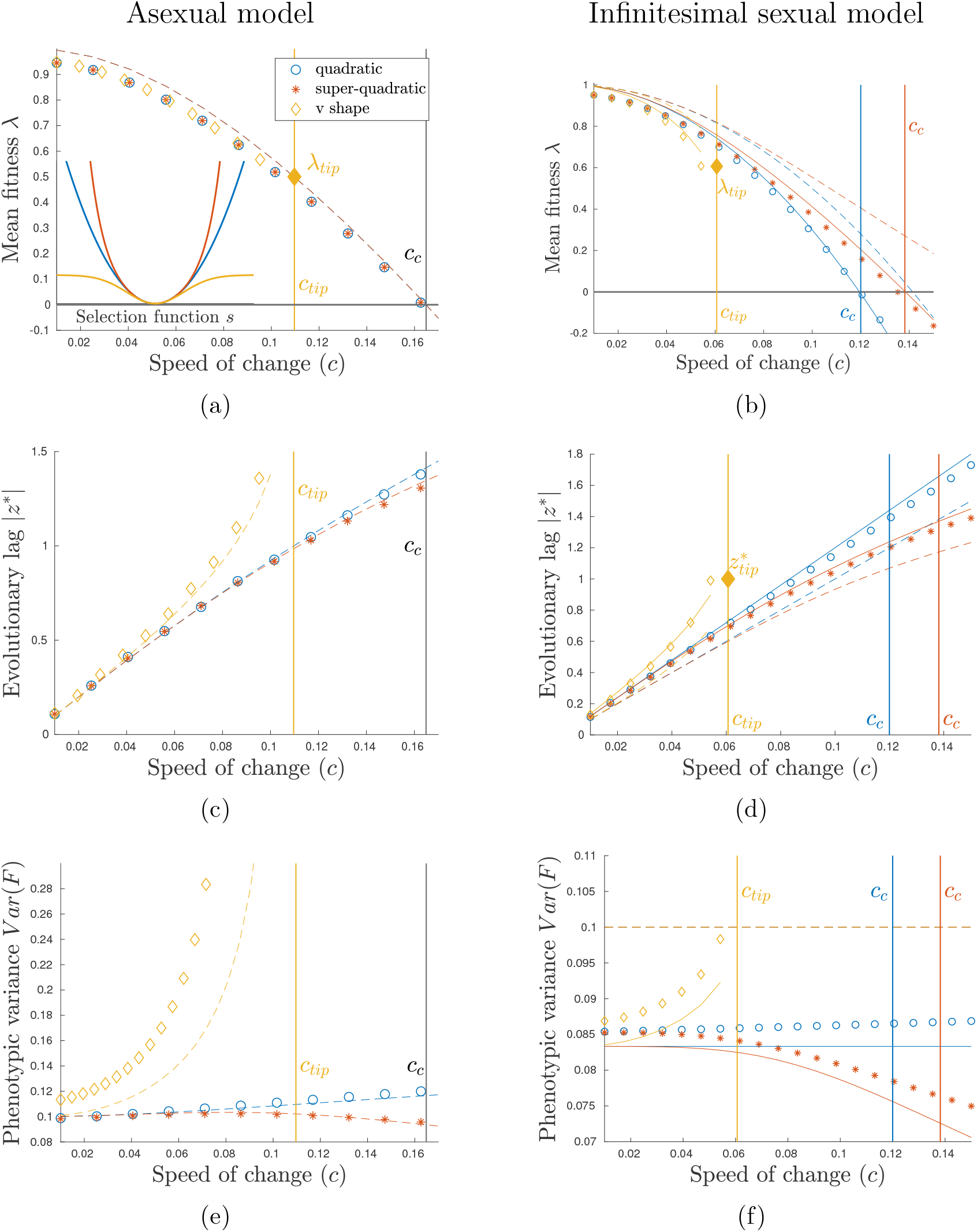
Influence of the speed of environmental change **c** for three different shapes of the selection function: quadratic function *m*(*z*) = *z*^2^/2 (blue curves), super–quadratic function *m*(*z*) = *z*^2^/2 + *z*^6^/64 (red curves) or bounded function *m*(*z*) = *m*_∞_(1 − exp(−*z*^2^/(2*m*_∞_)) (orange curves). Other parameters are: ***β*** = 1, **V**_sel_ = 1 and **V**_div_ = 0.01 and *m*_∞_ = 0.5 in the asexual model and **V**_div_ = 0.1 and *m*_∞_ = 1 in the infinitesimal sexual model. In the asexual model, the mutation kernel is Gaussian. We compare our analytical results (first approximation dashed lines and second approximation plain lines) with the numerical simulations of the stationary distribution of (8) (marked symbols) for both asexual and sexual infinitesimal model. The vertical lines correspond to the critical speeds for persistence **c**_*c*_, defined by (37) and the critical speed of tipping point **c**_tip_, defined by (35).

In the asexual model, the evolutionary lag at equilibrium is such that the mortality rate equals 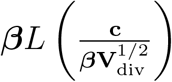 (see Table 2). The latter quantity increases with the rate of environmental change. As the mortality rate **m** increases when we move away from the optimal trait, the lag 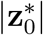 must also increase with respect to **c**.

In the infinitesimal model of sexual reproduction, the evolutionary lag at equilibrium is found where the gradient of selection (**m**′) equals 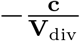, which increases in absolute value with the rate of environmental change **c** (see Table 2). In the convex neighborhood of the optimal trait, the gradient of selection (**m**′) is increasing with deviation from the optimum, hence the lag 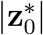 is increasing with respect to **c**. However, if the fitness function has both a convex and a concave part (as in the yellow curves in Fig. 3), there may be multiple equilibria fulfilling the condition in Table 2 (see Fig. 4(b)). In the concave part of the fitness function, the selection gradient is decreasing when **c** increases, and so would the lag (see dashed curve in Fig. 5(b)). However, heuristic argument and numerical simulations suggest that equilibrium points in the concave part of the fitness function are unstable (see Fig. 5(b) and more detailed discussion of this scenario below).

**Figure 4:**
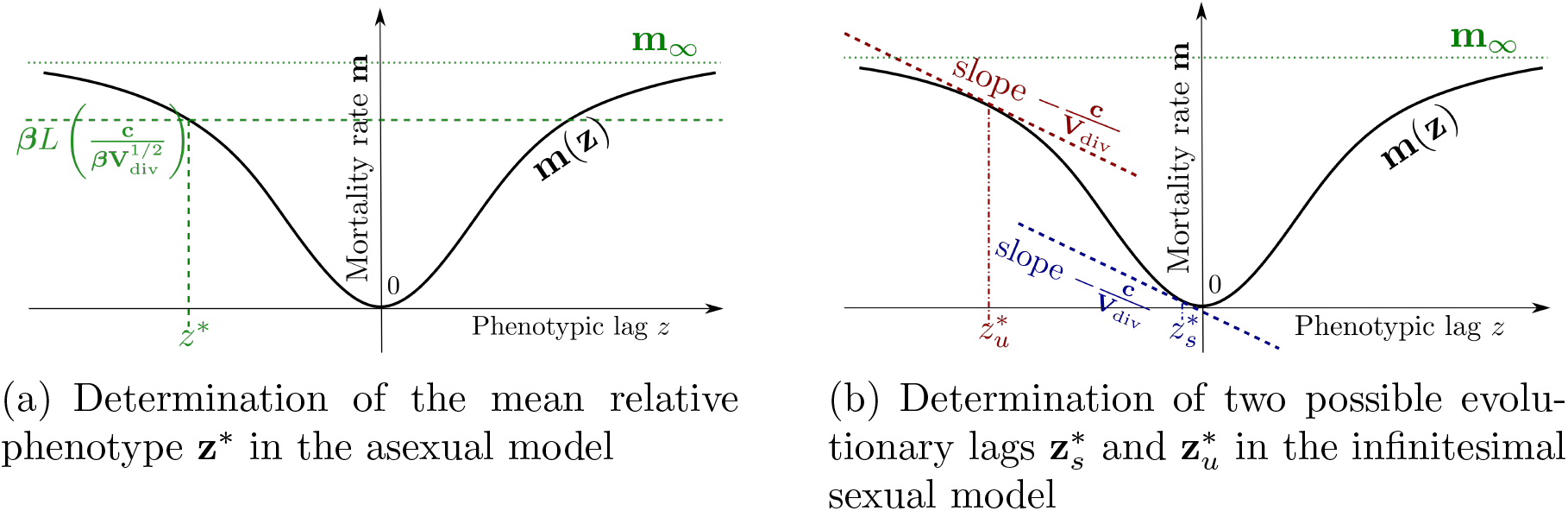
Graphical illustration of the two ways to characterize the mean relative phenotype **z**\* (in original units). (a) In the asexual model, the mean relative phenotype is found where the mortality rate **m** equals a specific value 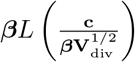. In this case we only have one possible lag **z**\* because **m**′(**z**\*) and **c** should have opposite signs. (b) In the sexual infinitesimal model, the mean relative phenotype **z**\* is found where the selection gradient **m**′ equals a specific value 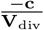. In this case, we may obtain two possible values, a stable point 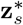 in the convex part of **m** and an unstable point 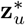 in its concave part.

**Figure 5:**
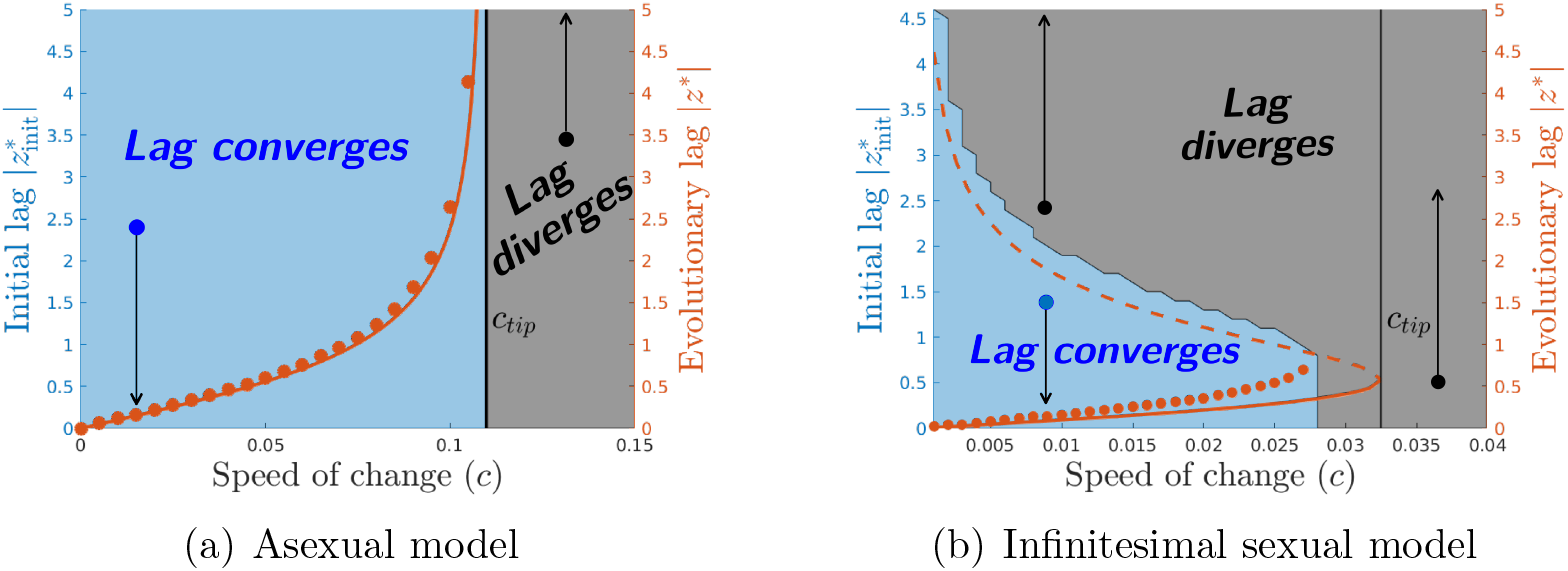
Effect of the initial lag on the persistence of the population with various rates of environmental change **c**. We compute numerically, the solutions of the time–dependent problem (2) with Gaussian initial conditions centered on various mean relative phenotypes 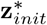 inducing an initial evolutionary lag 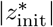 (left axes). We repeated this exploration for various speeds **c** ranging in (0, 1.5 **c**_tip_). For each case, we plot the evolutionary lag |*z*\*| at the final time of computations (red circles on right axes). We also compare with the analytical evolutionary lags given by the first line of Table 2 (red lines right axes): the plain lines corresponds to the stable trait (|*z*\*| in asexual model and 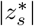 in infinitesimal sexual model) while the dashed lines corresponds to the unstable trait 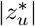 occurring in the infinitesimal sexual model. The grey region corresponds to initial data such that the final evolutionary lag diverges (black dots), while the blue region corresponds to initial data such that the final evolutionary lag converges (blue dots). In the asexual simulations, the mutation kernel is Gaussian.

#### The lag increases faster or slower than the speed of environmental change

Our analytical predictions suggest that a linear relationship between the rate of environmental change and the evolutionary lag is expected only under special circumstances. We indeed show that the rate of increase of the lag according to the speed of change **c** crucially depends on the shape of the selection in both the infinitesimal and asexual models (Fig. 3).

In addition, in the asexual model, this rate of increase will depend on the shape of the mutation kernel through the Lagrangian function *L*. Indeed, we can show from our formula in Table 2 that the lag increases linearly with the speed of change as soon as the function *c* ↦ *m*^−1^(*L*(*c*)) is linear. Thus, both the shape of selection and that of the mutation kernel interact to determine how the evolutionary lag responds to faster environmental change. If the selection function is quadratic (i.e. *m*(*z*) = *z*^2^/2), we can show from the convexity of the Lagrangian function *L* that the lag increases linearly with the speed only in the diffusion approximation *L*(*c*) = *c*^2^/2 (see Table 3 and blue curve in Fig. 2), while it increases sub–linearly for any other mutation kernels (see red, orange, purple and green curves in Fig. 2). We can further show that the lag in this scenario increases more slowly with the speed of environmental change when the kurtosis of the mutation kernel is higher (see SI D.4 for mathematical details). In Fig. 2, we compare four different mutation kernels with increasing kurtosis: uniform distribution kernel (red), Gaussian kernel (orange), double exponential kernel (purple) and Gamma kernel (green). In the asexual model, a fat tail of the mutation kernel thus tends to reduce the lag, even though this effect is most visible when the environment changes fast (Fig. 2).

To examine the effect of the shape of the selection function on how the evolutionary lag increases in faster changing environment, we now focus on the case of diffusion approximation in the asexual model (*L*(*c*) = *c*^2^/2), for the sake of simplicity, and compare it to the results in the infinitesimal model. In both cases, we can exhibit a simple criteria to decipher the nature of this increase. Let us first observe that, in those cases, the lag increases *linearly* with the speed if the selection function is quadratic (see Table 3 and the blue curves in Fig. 3). The lag however accelerates with the speed if *m* is *sub-quadratic* in the following senses (see orange curves in Fig. 3):

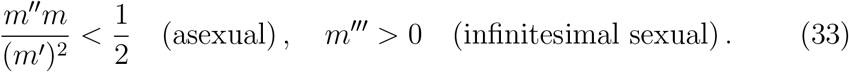

Conversely, the lag decelerates with the speed if *m* is *super-quadratic* in the following senses (see red curves in Fig. 3):

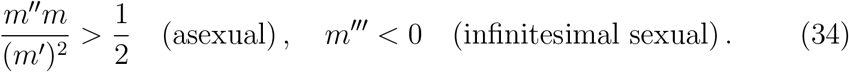

The criteria are of different nature depending on the model of reproduction (asexual versus infinitesimal). However, they coincide in the case of a homogeneous selection function *m*(*z*) = |*z*| ^*p*^ (*p* > 1). Indeed, selection is super-quadratic in both cases if and only if *p* > 2. More generally, the lag is reduced when the selection function has a stronger convexity in the sense of (34). This behavior is illustrated in Fig. 3.

#### The lag can diverge for a fast speed of environmental change

As observed by Osmond and Klausmeier (2017), we also find that the lag may diverge, i.e. grow infinite, when the selection function is too weak away from the optimum, and when the speed of environmental change exceeds some critical threshold. Interestingly, both the infinitesimal model and the asexual model exhibit such “evolutionary tipping point”, corresponding to *a critical level of an external condition where a system shifts to an alternative state* (van Nes et al., 2016). The underlying mechanisms are however qualitatively different in the two models, as explained below.

In order to illustrate this phenomenon, we consider a bounded selection function depicted in Fig. 3 (orange curve). We restrict to the diffusion approximation in the asexual case for the sake of simplicity. We find the following critical speed **c**_tip_,

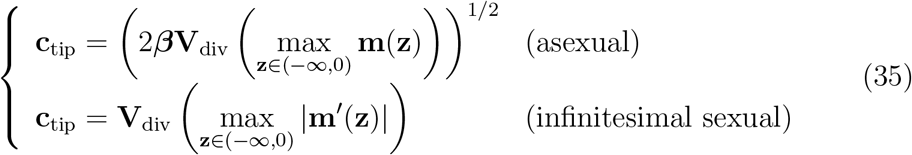

so that the lag is finite if and only if **c** < **c**_tip_, while the lag diverges if **c** > **c**_tip_ and the population cannot keep pace with the environmental change. The difference between the two formulas can be understood through graphical arguments (see Fig. 4). In the asexual model, the lag at equilibrium is found where the mortality rate equals a specific value, which increases with the speed of change **c**. This point is found where the selection function intersects an horizontal line, of higher elevation as **c** increases in Fig. 4. With a bounded mortality function, there is thus a finite value of **c** for which this critical quantity equals the maximal mortality rate, the latter being reached for an infinitely large lag. In the infinitesimal model, the evolutionary lag is found where the selection gradient equals a specific value increasing with **c**, see the graphical construction in (Osmond and Klausmeier, 2017, Fig.1B). With a bounded mortality function such as in Fig. 4, there are in general two equilibrium points characterized by such local slope, one stable in the convex part and one unstable in the concave part. As the speed of environmental change increases, so does the local slope at the two equilibria, which gradually converge towards the inflection point of the mortality function with the maximal slope. This point characterizes the maximal speed of environmental change for which there is a finite evolutionary lag. Above that critical speed of change, the lag grows without limit. We illustrate this phenomenon of severe maladaptation in Fig. 3 (see the orange curves).

Despite the existence of tipping points in both cases, the transition from moderate (**c** < **c**_tip_) to severe maladaptation (**c** > **c**_tip_) have different bifurcation signatures depending on the reproduction model. In the asexual model, the lag becomes arbitrarily large as the speed **c** becomes close to the maximal sustainable speed **c**_tip_. At the transition, the stable equilibrium state reaches infinity, which corresponds to a peculiar state where all individuals have the same fitness, and selection is not effective, reminiscent of a transcritical bifurcation. In contrast, in the infinitesimal model, the lag remains uniformly bounded up to **c**_tip_. At the transition, the stable equilibrium state merges with the unstable equilibrium state, through a saddle-node bifurcation.

We can also see a major difference between the two reproduction models when we look at the time dynamics (Fig. 5). We run simulations of equation (2) starting from various initial data centered at different traits (see crosses in Fig. 5). In the infinitesimal model, when the initial lag is beyond the unstable point 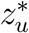, defined in Fig. 4(b), the lag diverges, whereas it converges to the stable point 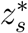, also defined in Fig. 4(b), if the lag is initially moderate. We see that the long term adaptation of the population to a changing environment does not only depend on the speed of change, but also on the initial state of the population. In the asexual model, the initial configuration of the population does not play a significant role in the long term dynamics of adaptation: we observe that the population can adapt whatever the initial lag is, if the speed of change is below **c**_tip_ (see Fig. 5). We can expect such difference because the lag at equilibrium is uniquely defined in the asexual model while it can take multiple values in the infinitesimal model if the function has an inflection point, a signature of bistability (see Fig 4).

### 4.2. The mean fitness

We now investigate the effect of the changing environment on the mean fitness of the population.

#### The mean fitness decreases with increasing speed of environmental change

In both scenarios the *lag load* △***λ***, defined as the difference between the mean fitness in a constant environment (***λ*** = ***β*** − ***μ***_0_, for a perfectly adapted population without standing load and **c** = 0) and the mean fitness under changing environment, is (unsurprisingly) given by the increment of mortality at the mean relative phenotype 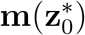

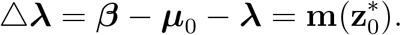

Since **m** is symmetrically increasing and the lag 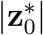 is increasing with respect to **c**, we deduce that the mean fitness decreases with respect to **c**. It is illustrated in Fig. 3 for different selection functions.

In the asexual model, the lag-load takes the following form

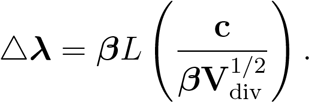

which is exactly the expression (22) in the original units with a speed **c**. Since *L* increases with the kurtosis of the mutation kernel, we deduce that higher kurtosis of the mutation kernel increases the mean fitness (see Fig. 2 and SI D.4). Thus the lag-load is maximal for the diffusion approximation.

#### The shape of selection affects the lag load in the infinitesimal model, but not in the asexual model

In the asexual model, the lag load only depends, at the leading order, on the speed of environmental change and the mutation kernel through the Lagrangian function *L* (20)-(22) and the variance **V**_div_ (see Table 2). It does not depend on the selection, as illustrated in Fig. 6(a) (dashed line). At the next order of approximation, the mean fitness however depends on the local shape of the selection function around the optimal trait through **V**_sel_ (10). The mean fitness is then predicted to decline as the strength of stabilizing selection around the optimum 1/**V**_sel_ increases, due to increasing standing load. These predictions are confirmed by our numerical simulations see Fig. 3(a) and 6(a).

**Figure 6:**
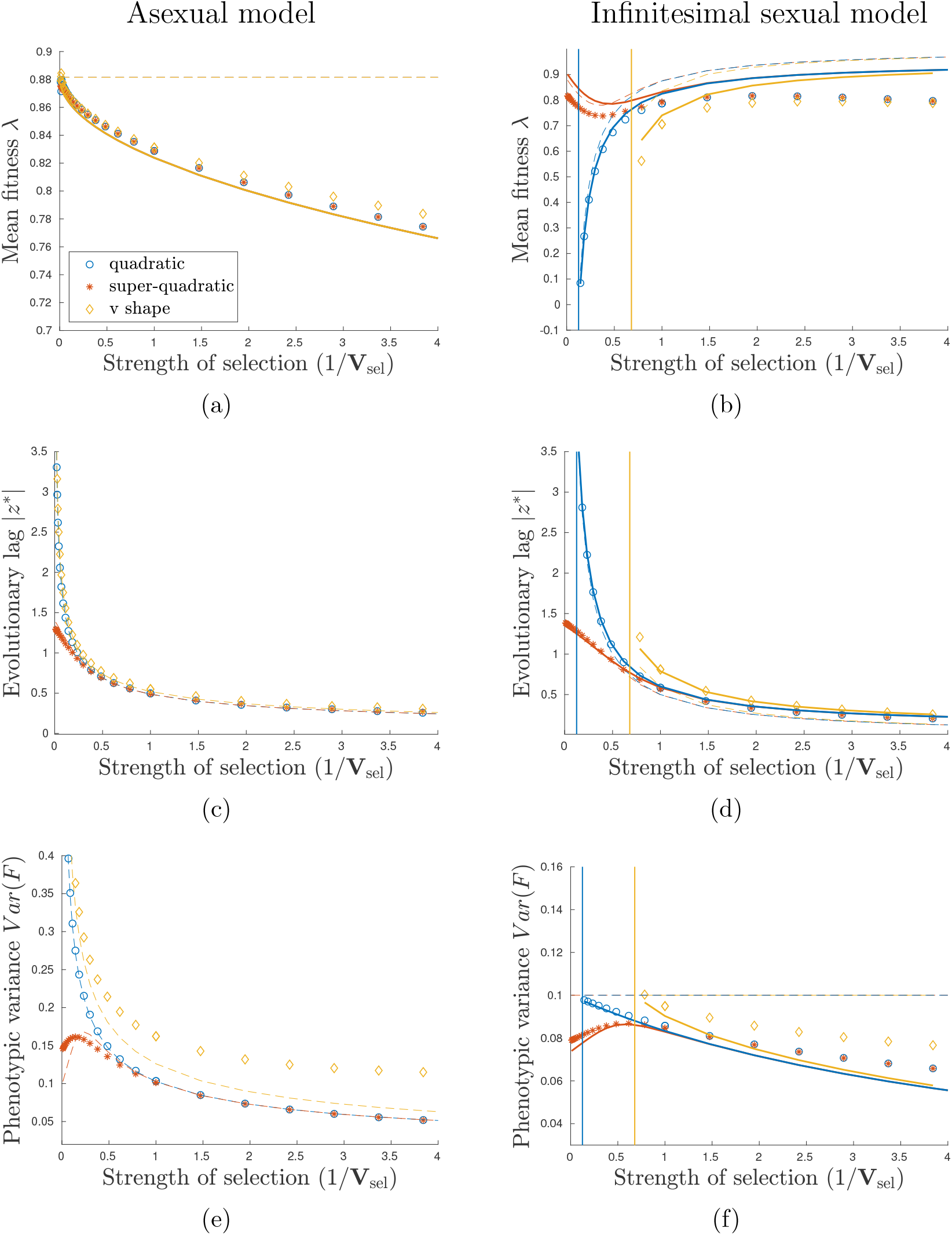
Influence of the strength of selection 1/**V**_sel_ on the mean fitness ***λ***, the evolutionary lag |**z**\*| and the phenotypic variance Var(**F**) at equilibrium in an environment changing at rate **c** = 0.05 and with three different selection patterns: quadratic (blue curves), super–quadratic (red curves) or bounded (orange curves). Other parameters are: ***β*** = 1, **V**_div_ = 0.01 for the asexual case and **V**_div_ = 0.1 for the sexual infinitesimal case and the intensity of selection 1/**V**_sel_ ranges from 10^−2^ to 4. We compare our analytical results (first approximation dashed lines and second approximation plain lines) with the numerical simulations of the stationary distribution of (2) (marked symbol) for both asexual and sexual infinitesimal model. In the asexual model, we only consider a Gaussian mutation kernel.

In contrast, the influence of the selection pattern is more intricate in the case of the infinitesimal model of reproduction. The lag load depends strongly on the global shape of **m** (see Fig. 3(b) and 6(b)). In particular, we see that for low strength of selection 1/**V**_sel_, the mean fitness crucially depends on the shape of selection. Mean fitness is higher in the scenario with super– quadratic selection than with quadratic selection, and lowest when selection is sub-quadratic in Fig. 3(b) and 6(b)). Moreover, the mean fitness increases with increasing strength of selection in the quadratic case, while it initially decreases for the super–quadratic case. However, for stronger strength of selection, the shape of selection has less importance. Our approximation allows us to capture those differences. For instance, in the quadratic case (blue curves in Fig. 6 and 3), we can see from Table 3 that the mean fitness increases with the strength of selection at the leading order, which corresponds to large value of **V**_sel_. However, when the strength of selection becomes stronger, antagonistic effects occur at the next order so that the fitness may decrease due to standing load, defined in (23) (Lynch and Lande, 1993; Lande and Shannon, 1996; Kopp and Matuszewski, 2014). This effect is illustrated in Fig. 6(b).

### 4.3. The phenotypic variance

In both asexual diffusion approximation and the infinitesimal model, the phenotypic variance does not depend on the speed of change **c** when the selection function is quadratic (see blue curves in Fig. 2(d) for asexual model and Fig. 3(f) for infinitesimal model). The phenotypic variance however increases with **c** if the selection function is *sub-quadratic* in the sense of (33) (see orange curves in Fig. 3). Conversely, the phenotypic variance decreases with **c** if the selection function is *super-quadratic* in the sense of (34) (see red curves in Fig. 3) – see details in SI E.

The phenotypic variance is less variable in the infinitesimal model than in the asexual model. It was expected from our analysis (see formula of Table 2) because the infinitesimal model tends to constrain the variance of the phenotypic distribution. Indeed, we know from previous analysis (Mirrahimi and Raoul, 2013; Barton et al., 2017), that in the absence of selection, the infinitesimal model generates a Gaussian equilibrium distribution with variance **V**_div_. Our analysis shows that under the small variance assumption, the phenotypic variance is close to this variance **V**_div_ and our numerical analysis shows that phenotypic variance slowly deviates from the genetic variance without selection **V**_div_, when either the speed of change increases or the strength of selection increases. This pattern is observed whatever the shape of selection. We can thus conclude that for the infinitesimal model under the small variance hypothesis, the phenotypic variance is not very sensitive to either selection (strength of selection or shape of selection) or the speed of environmental change.

Conversely, in the asexual model, the phenotypic variance is quite sensitive to the selection function. This is emphasized in the case of a bounded selection function. The phenotypic variance dramatically increases as the speed of change becomes close to the critical speed **c**_tip_ because the selection gradient becomes flat (see Table 2).

In the asexual model, the phenotypic variance is moreover sensitive to the shape of the mutation kernel. We see from Fig. 2(c) that the phenotypic variance generally increases with a fatter tail of the mutation kernel. There are however exceptions to this pattern (see for instance the Gamma mutation kernel at low speed of environmental change, green curves in Fig. 2). This situation, unexpected by our approximation, might be due to the fact that when the speed of change is low, the mutations with large effects are quickly eliminated by selection, which in turn reduces the phenotypic variance. This detrimental effect of large mutations when the environmental change rate is low has been also observed by Kopp and Hermisson (2009) and Collins et al. (2007).

### 4.4. Persistence of the population: the critical speed c_c_

The final outcome of our analysis is the computation of the speed **c**_*c*_ beyond which the population cannot keep pace with the environmental change (***λ*** < 0). In the general case, we can obtain the following approximation formula:

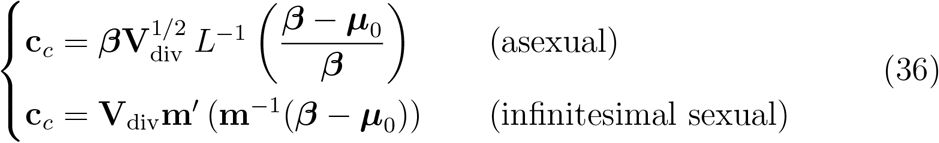

We can first observe that, in the small variance regime, the critical speed in the asexual model does not depend on the shape of the selection **m**, but on the mutation kernel through the Lagrangian *L* and the variance **V**_div_. Thus, for any selection function, the critical speed is the same (see Fig. 7(a)). Conversely, for the infinitesimal model, the critical speed crucially depends on the shape of the selection (see Fig. 7(b)). Moreover, we can mention that the discussion of the dependency of ***λ*** with respect to various parameters also holds naturally for **c**_*c*_.

**Figure 7:**
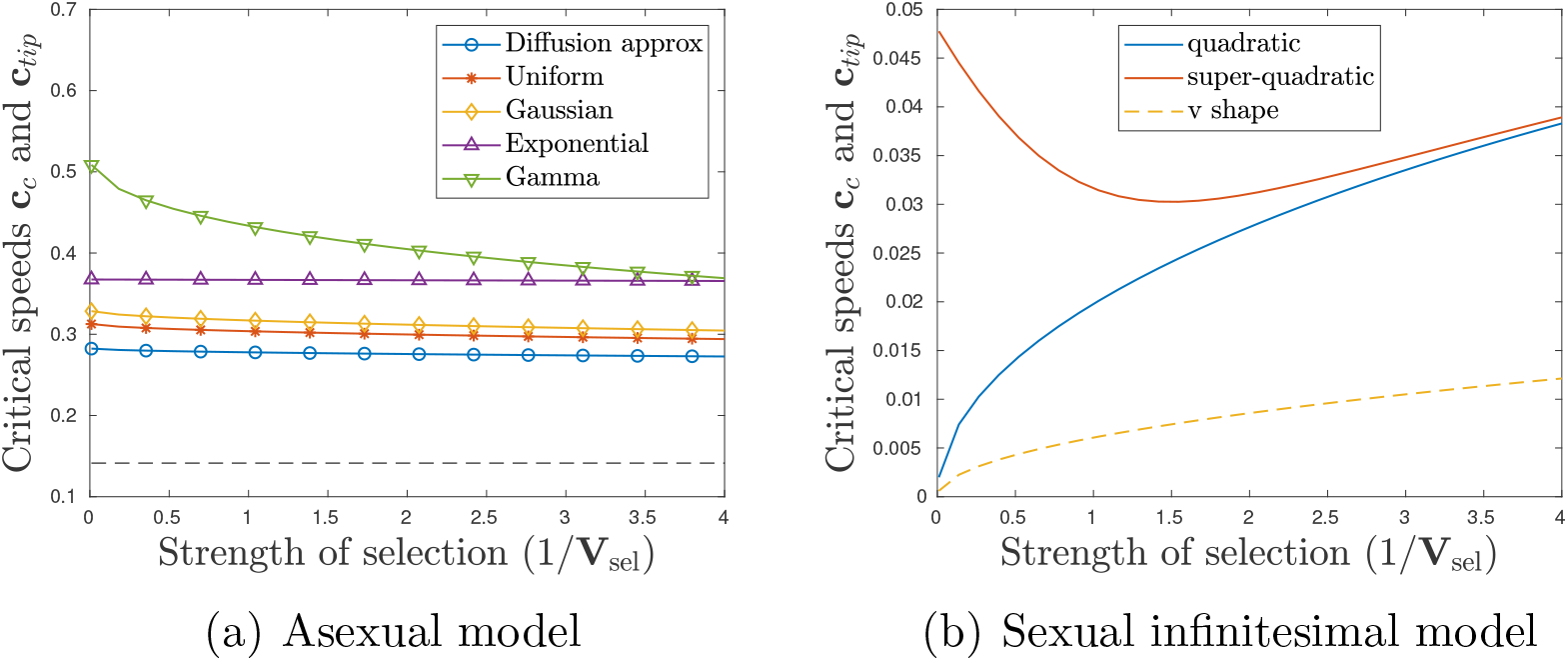
Critical speeds **c**_*c*_ and **c**_tip_ as the function of the selection strength 1/**V**_sel_ for: (a) the asexual model and (b) the sexual infinitesimal model. In panel (a), the plain curve corresponds to the critical speed **c**_*c*_ with different mutation kernel: Diffusion approximation (blue), Uniform distribution (red), Gaussian distribution (orange), Exponential distribution (purple curve) and Gamma distribution (green). The dashed line is the critical speed **c**_tip_. In panel (b), the curves correspond to different selection functions: quadratic (blue), super-quadratic (red) and v-shape (orange). The plain curves corresponds to the critical speed **c**_*c*_ while the dashed curve to the critical speed **c**_tip_.

When we consider the diffusion approximation for the asexual model (*L*(*v*) = *v*^2^/2) and the quadratic selection function **m**(**z**) = **z**^2^/(2**V**_sel_), we obtain the following formula, including the next order term:

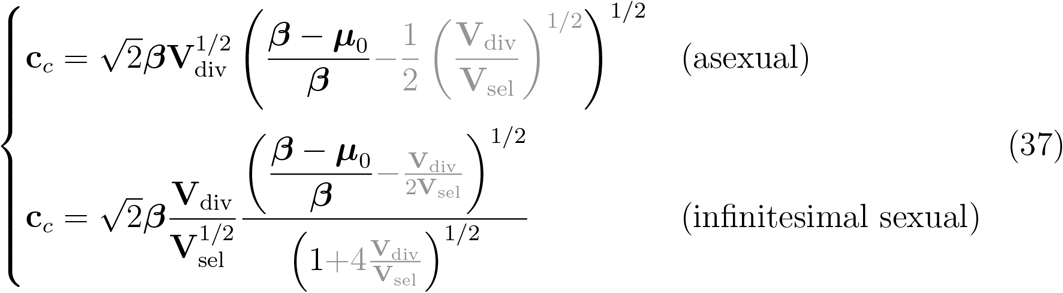

In the asexual case, the formula (37) is in agreement with previous results where it was assumed that the relative phenotype **z** is normally distributed in the population, which corresponds in our framework to assuming that the equilibrium distribution **F** is Gaussian (see Lynch et al., 1991; Lynch and Lande, 1993).

Moreover, in the asexual case with ***μ***_0_ = 0, our formula (37) is consistent with the classical formula given with the phenotypic variance as a parameter:

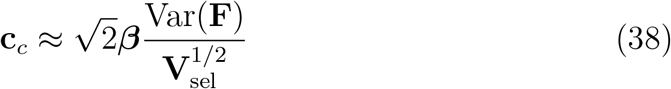

see for instance Eq. [A6] in (Kopp and Matuszewski, 2014). This approximation (38) also holds true in the sexual infinitesimal case. Although this simple formula is a good approximation in a general setting, it might be misleading, as it omits some possible compensation, such as the selection strength 1/**V**_sel_, which disappears in the case of asexual reproduction because it also affects Var(**F**).

#### Numerical approximations for finite population

Here, we compare our approximation formula described in Table 2, with the outcomes of the stochastic model, defined in SI H, when the number of individuals **N** is small (**N** is equal to 10^2^ or 10^3^) and the selection scenario varies as in Fig. 3.

When the speed of change is slow compared to the critical speeds, our approximations seem accurate in the sense that the approximation error usually falls on our confidence intervals (see Fig. S4-S6). In the infinitesimal sexual model, our approximations also do well when the speed is close to the critical threshold. In this model, we know that the population adapt thanks to the bulk of the population, which moves forward. Thus, even if the size of the population decreases, many individuals remains at the dominant trait. The size of the population does not have a critical influence on the adaptation response.

However for the asexual model, when the speed of change increases, our approximations become less accurate. In this model, only the individuals near the optimal trait help the population to adapt. Thus when the speed increases, the proportion of individuals near the optimal trait decreases because the lag increases. Moreover, when the population size decreases, the actual number of individuals at the optimal trait may be zero, which may lead to an additional burden, and possibly the extinction of the population before the critical value **c**_*c*_ is reached (Calvez et al., 2023). In particular, we see in Figures S4-S6 (a) that the mean fitness of the population drops below 0 for fifty percent of the simulations when the speed is close to the critical speed. Thus the effect of the population size is stronger for the asexual model than for the infinitesimal sexual model.

### 4.5. Numerical predictions for the whole distribution of phenotypes Quality of the approximation

For the asexual model, we only compare the simulation results with our first order approximation stated in Table 2 (black colored), except for the variation of the mean fitness with respect to the strength of selection, where it is preferable to take into account the standing load that appears at the second order of approximation (see gray colored formula in Table 2). We can first observe from Fig. 6 that our approximations are accurate when *ε* = (**V**_div_/**V**_sel_)^1/2^ is small (see value of 1/**V**_sel_ < 0.5 in Fig. 6). The scale of Fig. 6(a) is of order *ε*, which is why the first order approximation seems less accurate than the second order approximation. This was expected since the standing load, which increases with the strength of selection, occurs at the second order of the approximation. The approximations of **z**\* and ***λ*** remain efficient even when *ε* increases (see Fig. 2 and 3 for small value of **c**). However, we see that the approximations deviate from the simulations when the speed of change increases and reaches the critical value **c**_*c*_ (see Fig. 2 and 3) or when the mutation kernel becomes leptokurtic (see green curves of Fig. 2). The approximation of the phenotypic variance is more sensitive to the parameter *ε*. When **c** and *ε* are small it is accurate (see Fig. 2). However, when the speed increases, the approximation diverges from the simulations even if *ε* is small (see Fig. 2 and 3).

For the infinitesimal model, we have compared our simulations to our first order approximation, as well as the second order approximation stated in Table 2 (first order approximation is black colored and second order approximation is gray colored). The first order approximation of **z**\* and ***λ*** are efficient only when *ε* is really small, while the first order approximation of the phenotypic variance may deviate from the simulation value even for small *ε* (see red curve Fig. 6(f)). However, the second approximations are really precise for small value of *ε* (see Fig. 6) and they remain accurate when *ε* increases and **c** increases (see Fig. 6 and 3).

#### Comparing simulations to the approximation for the entire distribution

We compare the simulated equilibrium distribution **F** with our analytical approximations (Fig. 8): the first order approximation corresponds to **F**_0_ = exp(−*U*_0_*/ε*^*γ*^), where *U*_0_ satisfies respectively the differential equation (19) (asexual model) or 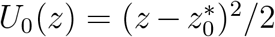 (infinitesimal sexual model), and *γ* is respectively equal to 1 in the asexual model and 2 in the infinitesimal case; and the second order approximation **F**_1_ = exp(−*U*_0_*/ε*^*γ*^ −*U*_1_), where *U*_1_ satisfies respectively equation (D.12) (asexual model) or the non–local functional equation (30) (infinitesimal model). Our simulations are performed with an *ε*^*γ*^ = 0.1, which is not that small.

**Figure 8:**
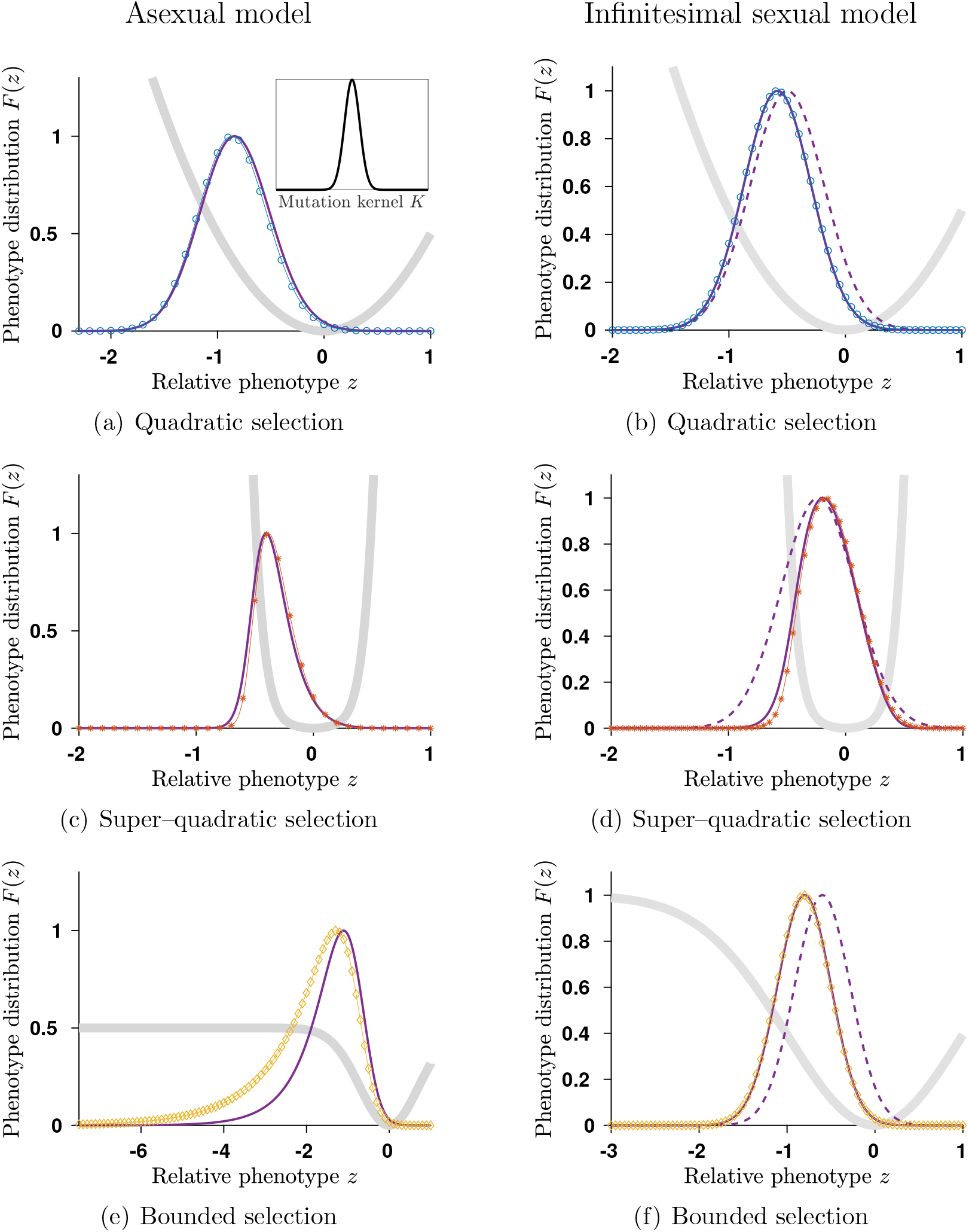
Mutation-selection equilibria **F** in changing environment with three different shapes of selection: (a)-(b) quadratic function *m*(*z*) = *z*^2^/2 (blue circled marked curves); (c)-(d) super-quadratic function *m*(*z*) = *z*^2^/2 + *z*^6^/64 (blue star marked curves); (e)-(f) bounded function *m*(*z*) = *m*_∞_(1 − exp(−*z*^2^/(2*m*_∞_)) (orange diamond marked curves). The speed of environment change is **c** = 0.09 in the asexual model while it is **c** = 0.05 in the infinitesimal sexual model so that itremains below the critical speeds **c**_*c*_ and **c**_tip_ and the distribution deviates significantly from the Gaussian distribution approximation. Other parameters are: ***β*** = 1, **V**_sel_ = 1 and **V**_div_ = 0.01 and *m*_∞_ = 0.5 in the asexual model and **V**_div_ = 0.1 and *m*_∞_ = 1 in the infinitesimal sexual model. We compare simulated equilibria distribution **F** (marked curves) with our analytical results (first order results dashed curves and second order results plain curves). For the asexual scenario, we used the Gaussian kernel.

In the asexual model, we can observe that the first order analytical approximation is really efficient at tracking the shape of the entire distribution for both super-quadratic and quadratic selection, even if *ε* is not so small (Fig. 8). For the bounded selection, our first order approximation fails to fit the left tail of the distribution, mainly because the speed of environmental change is close to the critical speed.

In the infinitesimal model, we can observe that the first order Gaussian approximation is not precise enough to track the entire distribution (Fig. 8). We need the second order approximation to fit the distribution. This is a direct consequence of our analysis, where we observe that we need the second order approximation to define the first order approximation of the lag **z**\* and the mean fitness ***λ***.

We also compare our approximations of the phenotypic distribution with the empirical distribution of the IBM model, described in SI H, for the scenarios described in Fig. 8. When the size of the population is large (of order **N** = 10^4^), our approximations are accurate and fit with the empirical distribution of the stochastic model (see Fig. S3).

#### The skewness and the kurtosis of the phenotypic distribution

To go further in understanding the effect of a changing environment, we looked at the skewness and the kurtosis of the distributions. Those two indicators allow us to test whether the distribution **F** can be well approximated by the Gaussian distribution.

In the asexual model, with a Gaussian kernel *K*, we can observe from Fig. 9 that, even for quadratic selection, the distributions differ from a Gaussian distribution: they are skewed and leptokurtic, which means that their kurtosis are higher than the kurtosis of the Gaussian distribution with same mean and variance. So the Gaussian distribution fails to track the exact distribution of the trait around the mean trait of the population in a changing environment. This phenomenon is enhanced when the selection function differs from the quadratic function (see Fig. 9 diamond curves and Fig. 8). In addition, we see that, when the selection function is super–quadratic, the distribution has a positive skew, while, for a bounded selection function, it has a negative skew.

**Figure 9:**
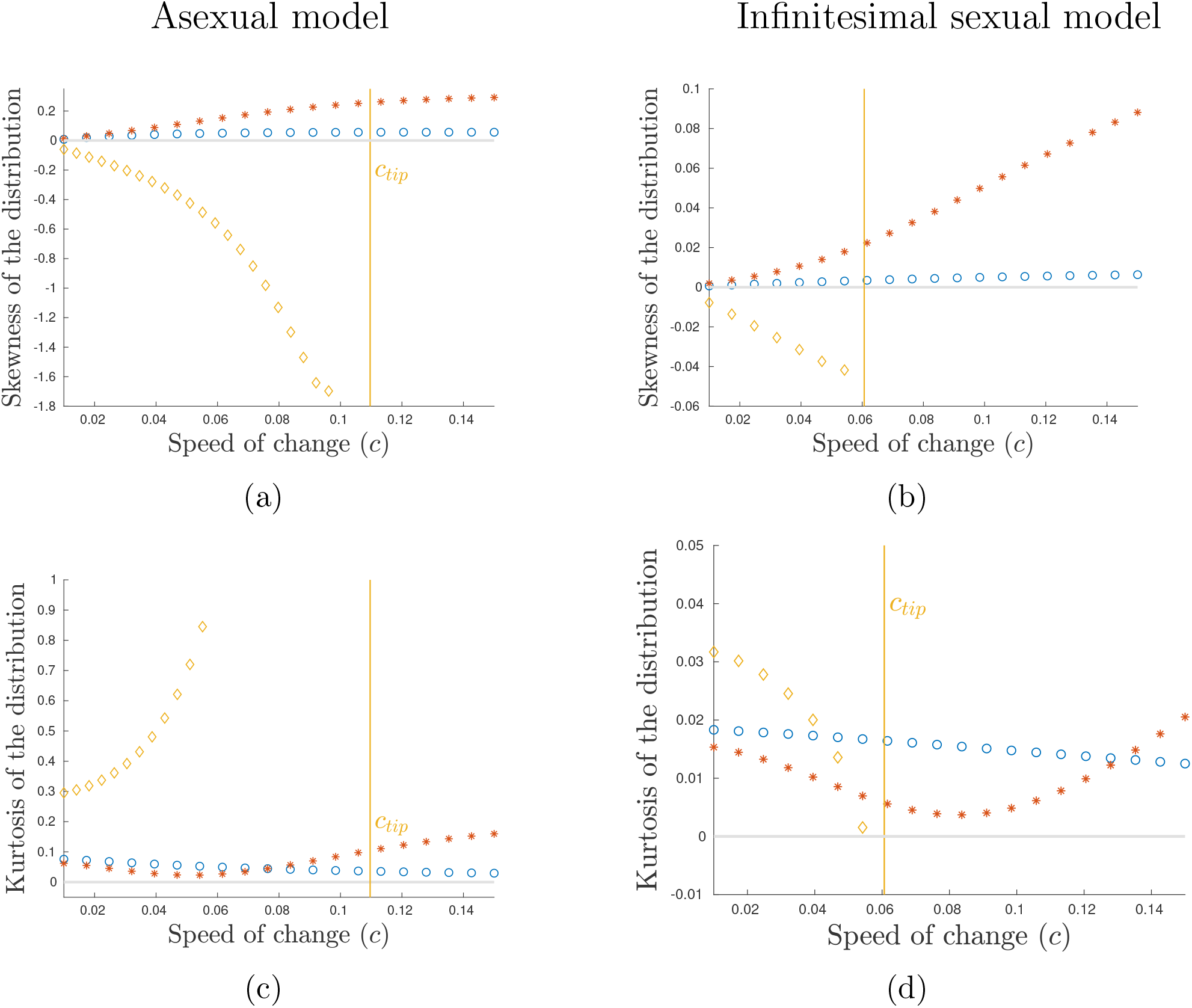
Influence of the speed of environmental change **c** on the skewness and the kurtosis of the distribution **F** for three different shapes of selection: quadratic function *m*(*z*) = *z*^2^/2 (blue circles), super-quadratic function *m*(*z*) = *z*^2^/2 + *z*^6^/64 (red stars) or bounded function *m*(*z*) = *m*_∞_(1−exp(−*z*^2^/(2*m*_∞_)) (orange diamonds). Other parameters are: ***β*** = 1, **V**_sel_ = 1 and **V**_div_ = 0.01 and *m*_∞_ = 0.5 in the asexual model and **V**_div_ = 0.1 and *m*_∞_ = 1 in the infinitesimal sexual model. In the asexual model, the mutation kernel is Gaussian.

Conversely, in the infinitesimal case, the Gaussian distribution well approximates the equilibrium distribution in general. This was already described by our approximation formula (29) in Section 3.2. We can see that the kurtosis of the equilibrium distribution remains close to zero for any speeds of change and any selection functions. However, when the selection function is either super-quadratic or bounded, we can observe from Fig. 8 and 9 that the distribution of phenotypes in the infinitesimal model also becomes skewed as the speed increases. The skew of the distribution corresponds to regions where the gradient of selection is low, with the same pattern as in the asexual model.

## 5. Discussion

We have pushed further a recent methodology aimed at describing the dynamics of quantitative genetics models in the regime of small variance, without any *a priori* knowledge on the shape of the phenotype distribution. This methodology combines an appropriate rescaling of the equation with Taylor expansions on the logarithmic distribution.

### Small variance asymptotics

Our approach differs from the previous studies based on the cumulant generating function (CGF), which is the logarithm of the Laplace transform of the trait distribution, here *C*(*t, p*) = log (∫*e*^*pz*^*f*(*t, z*) *dz*). In his pioneering work, Burger (1991) derived equations for the so-called cumulants, which are the coefficients of the Taylor series of the CGF *C*(*t, p*) at *p* = 0. However this system of equations is not closed, as the cumulants influence each other in cascade. This analysis was revisited in (Martin and Roques, 2016) in the asexual model, using PDE methods. They derived an analytical formula for the CGF itself, but restricted it to a directional selection, when the trait represents the fitness itself. This was further extended to a moving optimum in (Roques et al., 2020). However, they made the crucial assumption of the Fisher Geometric Model for selection, which is analogous to our quadratic case, and diffusion for mutations, for which it is known that Gaussian distributions are particular solutions. The common feature with our present methodology is the PDE framework. Nevertheless, we focus our analysis on the logarithm of the trait distribution itself, as it is commonly done in theoretical physics to reformulate the wave-function in terms of its action (see SI D.3 for heuristics on this approach). This strategy is well-suited to provide precise approximations with respect to a small parameter, for instance the wavelength in wave propagation (geometric optics) and the Planck constant in quantum mechanics (semi-classical analysis), and the phenotypic variance in our theoretical biology setting.

Here, the small variance regime corresponds to relatively small effect of mutations compared to the strength of stabilizing selection. Under this regime, little variance in fitness is introduced in the population through either mutation or recombination events during reproduction. And the strength of selection acting on mutations arising in well-adapted population is weak. This regime is usually referred to as the “weak selection – strong mutation” regime. However, population suffering from maladaptation due for instance to a changing environment, can experience strong effect of selection that may drive the population to extinction.

Under the small variance regime, we could describe analytically the phenotype distribution (see Fig. 8), and assess the possible deviation from the Gaussian shape. We further gave analytical approximations of the three main descriptors of the steady state: the mean relative phenotype, the mean fitness, and the phenotypic variance (see Table 2). We also compared our deterministic approximations with the outcomes of stochastic simulations with a finite number of individuals (see SI H). Stochastic simulations are in good agreement when the number of individuals is large enough, or when the speed of change is not too close to the critical speed **c**_*c*_ in the asexual case. Furthermore, in the infinitesimal sexual model, our approximations seems really precise even when the size of the population shrinks as the speed of change increases. In this case, the variance is constrained to remain nearly constant, which forces the bulk of the population to adapt and prevent random drift to drive the population towards extinction. However, in the asexual model, we observe large discrepancies when the speed of change approaches the critical threshold. More precisely, in the asexual model, the dynamics of adaptation relies upon those individuals which are the fittest, as can be illustrated by the ancestral lineages (Patout et al., 2020; Calvez et al., 2022b). In an infinite population, the fittest individuals are certainly at the optimal trait. This actually explains why the lag load does not depend on selection at the leading order. However, in finite populations, when the speed of change increases, the lag increases, thus reducing the chance to find individuals with an optimal trait. This sampling effect induces an additional burden to the population, resulting in an increase of maladaptation, which may lead to extinction of the population, not predicted by the deterministic model of infinite population size (Calvez et al., 2023). This negative feedback between maladaptation and phenotypic variance, called “mutational meltdown” by Lynch and Gabriel (1990) in the context of the evolution of small population by mutation-selection, has already been observed numerically for small sexual populations subject to fast environmental change by Burger and Lynch (1995).

Noticeably, the two different models of reproduction, assuming either asexual reproduction, or infinitesimal sexual reproduction with an infinite number of freely recombining loci (the infinitesimal model), could be handled in a unified framework. This allows discussing similarities and differences between the two models, which are frequently used in analytical models of adaptation to changing and/or heterogeneous environments. However, the two models are subject to different scaling regimes, as exemplified by the differences in the phenotypic variance at equilibrium (see Tables 2 and 3), or by the different formulas of the critical speed (36). This discrepancy is an outcome of our mathematical analysis, which aims to capture the shape of the equilibrium phenotypic density in the regime of small variance *ε* ≪ 1. However, it is interesting to note that this result is consistent with previous predictions of the literature about the phenotypic variance at equilibrium in a constant environment in the asexual and infinitesimal model (Bürger, 2000; Barton et al., 2017). Those predictions were derived under the assumption that the phenotypic distribution is well approximated by a Gaussian distribution, an assumption that we here have relaxed. More precisely, under mutation-selection balance, the phenotypic variance of an haploid asexual population is well approximated under the Gaussian regime by Var(**F**) = (**V**_div_**V**_sel_)^1/2^ = *ε***V**_sel_ (see Bürger, 2000, and Table 3). While the phenotypic variance of a population following the infinitesimal model is approximately Var(**F**) = **V**_div_ = *ε*^2^**V**_sel_ (Barton et al., 2017). As a result, we see that the phenotypic variance in the asexual model is of order *ε*, while it is of order *ε*^2^ for the infinitesimal sexual model. However, we here deal with changing environment with general selection functions and various mutation kernels in the asexual case, for which analytical approximations are unknown, up to our knowledge. Without *a priori* estimates to justify our scaling, we looked for characteristic scales from a mathematical analysis (see section 2.2 and SI SI B). Noteworthy, our analysis shows that, despite the fact that the phenotypic distribution can deviate significantly from a Gaussian shape (see Fig. 8), the phenotypic variance scale remains of the same *ε* order in our analysis, in line with (Diekmann et al., 2005; Barles et al., 2009; Lorz et al., 2011) for the asexual case, and (Calvez et al., 2019) for the infinitesimal sexual case.

### Relaxing the Gaussian distribution assumption

Our analytical framework allows us to relax the assumption of a Gaussian distribution of phenotypic values, commonly made by several quantitative genetics models of adaptation to a changing environment with a moving optimum, both in the case of sexually (e.g. Burger and Lynch, 1995; Osmond and Klausmeier, 2017) and asexually reproducing organisms (e.g. Lynch et al., 1991). Consistently with previous simulations and analytical results (Turelli and Barton, 1994; Bürger, 1999; Jones et al., 2012), our results show that we expect stronger deviations from a Gaussian distribution of phenotypes if the selection function departs from a quadratic shape, if the mutation model departs from a simple diffusion, if reproduction is asexual rather than well described by the infinitesimal model, and/or if the environment changes relatively fast. We in particular recover the observation made by Jones et al. (2012) in their simulations that the skew of the phenotypic distribution is greater in absolute value in faster changing environments, but we further predict that the sign of this skew critically depends on the shape of the selection function away from the optimum, an observation that could not be made by their simulations that only considered quadratic selection.

### Universal relationships

Interestingly, despite deviations from the Gaussian distribution, our predictions in the regime of small variance for the mean relative phenotype, or the critical rate of environmental change, are consistent with predictions of past quantitative genetics models that have assumed a constant phenotypic variance and a Gaussian distribution of phenotypes. We discuss below the links between the present results and those past predictions and how they provide new insights. As a direct consequence of the small variance assumption, the two following relationships, linking the three main descriptors of the population (the mean relative phenotype, mean fitness and phenotypic variance), hold true, whatever the model of reproduction (either asexual or infinitesimal):

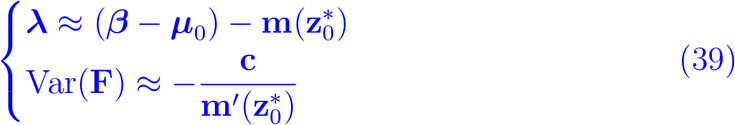

The first relationship corresponds to the demographic equilibrium, when the mean fitness is the balance between (constant) fecundity and mortality at the mean relative phenotype. The second one corresponds to the evolutionary equilibrium, when the speed of evolutionary change (as predicted by the product of phenotypic variance and the selection gradient) equals the speed of change in the environment. Note that our model assumes for simplicity that the phenotypic variance is fully heritable. Those relationships can be deduced directly from equations in dimensionless units (14)-(15)

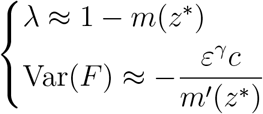

Although the reproduction model does not affect the demographic relationship, it influences the evolutionary relationship through the scaling exponent *γ*(*γ* = 1 for asexual reproduction and *γ* = 2 for infinitesimal sexual reproduction). Similar equations appear in quantitative genetics models assuming a Gaussian phenotypic distribution and a constant phenotypic variance. In particular, with quadratic selection, the second relationship allows us to recover the following results of Burger and Lynch (1995) and Kopp and Matuszewski (2014):

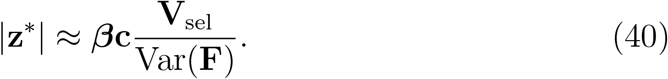

However, the two relationships (39) are not enough to compute the three descriptors, if one does not consider the phenotypic variance Var(*F*) as a fixed parameter, as previous studies often did. Our small variance approximations allows us to predict the value of the phenotypic variance in a changing environment in the two models, where previous studies have generally used simulations (e.g. Bürger, 1999) to examine how the evolution of the phenotypic variance affects the adaptation of sexual and asexual organisms in a changing environment. Many of our results are ultimately explained by the fact that the evolution of the phenotypic variance is under very different constraints under the asexual model and the infinitesimal model.

In the asexual model, the evolution of the phenotypic variance is not strongly constrained and has in particular no upper bound. The mean fitness *λ* does not depend on the shape of the selection function at the leading order (see (21) and Table 2), but only on the speed of environmental change and on the mutation kernel. Once the mean fitness is determined, the mean relative phenotype *z*\* and the phenotypic variance Var(*F*) are deduced from respectively the first and the second relationship in (39). The phenotypic variance then strongly depends on the shape of the selection in the asexual model. In contrast, in the sexual infinitesimal model, we found that the phenotypic variance Var(*F*) does not depend on the shape of the selection function at the leading order (see (29) and Table 2). The mechanism of inheritance in the infinitesimal model indeed constrains the value of the phenotypic variance at equilibrium. Then, the mean relative phenotype *z*\* and the mean fitness *λ* are deduced from respectively the second and the first relationship (39). Most of our predictions (discussed below) are a consequence of this core discrepancy between the two models.

### Mean fitness weakly depends on selection in the asexual model, but not in the infinitesimal model

In the asexual model, ***λ*** depends on **m** only at the second order through the strength of selection around the optimal trait 1/**V**_sel_ = **m**′′(0) (10). Hence, up to a reasonable accuracy, the mean fitness depends (weakly) on the local shape of the selection pattern around the optimal trait, even if the population can be localized around a mean relative phenotype far from the optimal trait. This happens because, in a gradually moving environment, the asexual population is constantly regenerated by the fittest individuals. This phenomena is apparent when tracing back lineages in the population at steady state: it was proven independently by Patout et al. (2020) and Calvez et al. (2022b) that the typical trajectories of ancestors of individuals sampled uniformly in the population converge to the optimal trait backward in time. In contrast, the mean fitness strongly depends on the shape of the selection function in the infinitesimal sexual model. It appears clearly in the quadratic case where **V**_sel_ enters into the formula for the mean fitness at the leading order (Table 3). In particular, we recover the previous finding that weak selection represents a “slippery slope” in a changing environment, leading to a lower mean fitness, when effects of selection on the evolution of phenotypic variance are neglected (Kopp and Matuszewski, 2014). Again, it is interesting to link this finding to the behavior of the typical trajectories of the ancestors in the infinitesimal model, which converge to the mean relative phenotype backward in time (Patout, 2019, Chapter 5).

### The shape of selection has strong effects on the evolution of the mean relative phenotype and phenotypic variance under both the asexual and infinitesimal models

In both models, however, the exact shape of the selection function away from the optimum has noticeable consequences for the evolution of the lag between the mean phenotype in the population and the moving optimum, and for the evolution of the phenotypic variance, especially in fast changing environments. There is unfortunately very scarce empirical evidence about the exact shape of fitness landscapes and how much they deviate from a quadratic, due to the difficulty to estimate precisely the shape of such fitness functions. However, some empirical studies reviewed in (Agrawal and Whitlock, 2010) suggest strong deviations from a quadratic shape. For instance, selection can become weaker with increasing maladaptation, due to lower bound on fitness (Agrawal and Whitlock, 2010). The effect of such selection can only be observed when the population is really maladapted, which might be the case for populations facing rapid environmental change. Under this scenario, Osmond and Klausmeier (2017) have shown that selection can constrain evolution, by limiting the ability of population to evolve and persist in a directional environmental change. Most models however assume, for mathematical convenience and in the absence of strong empirical support for an alternative, a quadratic selection function. Our analysis allows considering a broad diversity of selection functions and also to draw general conclusions about how their shape may affect the evolution of the phenotypic distribution. In both asexual and infinitesimal models, we found, consistently with previous predictions (reviewed in Kopp and Matuszewski, 2014), that the lag increases with the speed of environmental change: however there is a linear relationship between the two only when assuming a quadratic selection function. When the selection function is super-quadratic (and selection much stronger away from the optimum), this puts a brake on maladaptation and the evolutionary lag does not increase as fast when the environment changes more rapidly. For the same reason, the phenotypic variance then declines when the environment changes faster in the super-quadratic selection scenarios. Conversely, with a sub-quadratic selection function, the weakening of selection away from the optimum results in larger lags, accelerating maladaptation with increasing speed of environmental change and increasing phenotypic variance. There has been little discussion yet in the theoretical literature of the consequences of the exact shape of selection in changing environments (see however (Osmond and Klausmeier, 2017; Klausmeier et al., 2020) and discussion of tipping-points below). In a constant or stationary environment with weak fluctuations, the mean phenotype value is never very far from the optimum and the quadratic selection is an adequate approximation. However, the present results suggest that further empirical investigation of the shape of the fitness landscape far from the optimum is critically needed to understand how much populations may depart from the optimal phenotypic value.

### Evolutionary tipping points

The case of sub-quadratic selection functions has recently attracted some interest, since it was discovered that the weakening of selection away from the optimum could lead to evolutionary tipping points: above some critical speed of environmental change, the evolutionary lag grows without limit and the population abruptly collapses without much warning signal (Osmond and Klausmeier, 2017; Klausmeier et al., 2020). This behaviour is very different from the dynamics of the lag under classic models of quadratic selection on moving optimum. Osmond and Klausmeier (2017) assumed a Gaussian distribution of phenotypes and a constant phenotypic variance and compared their analytical results to simulations of a sexually reproducting population. Klausmeier et al. (2020) went on to show that non quadratic fitness function with inflection points, leading to such tipping points, could emerge from various realistic ecological feedbacks involving density-dependence or interactions with other species. Our analytical results allow us to predict the critical speed at which the evolutionary tipping points occur. In particular, we show that the infinitesimal tipping points occurs at the maximal rate of evolution, which corresponds to the product of the phenotypic variance and the maximal selection gradient. This relationship was already derived in a particular case by Osmond and Klausmeier (2017). We furthermore show that evolutionary tipping points also emerge in the asexual model, but with a different signature. In the asexual model, there is only one possible equilibrium for each value of the speed of environmental change. Again, ultimately, this unique equilibrium is due to the fact that the variance evolves more freely in the asexual model, which allows any variant close to the optimal trait to become dominant in the population (Patout et al., 2020). As the speed increases towards the critical value *c*_tip_, the lag diverges (Figure 4(a)-5(a)). As a result, the variance gets arbitrarily large and the skewness becomes negative, which shows that more individuals lag behind the mean relative phenotype. Conversely, in the infinitesimal model, the variance is constrained to remain nearly constant, which forces the bulk of the population to adapt. As a result, multiple equilibria exist, which determine several basins of stability, up to the critical value *c*_tip_. The lag remains bounded in the vicinity of the tipping point, determining a characteristic range for the basin of attraction of the origin (Figure 4(b)-5(b)). The lag can diverge, even if *c* < *c*_tip_, for maladapted initial distributions concentrated far from the origin. This corresponds to a population that cannot keep pace with the environmental change because they are initially maladapted, possibly due to some transient change in the environment of major effect.

### Effect of the mutation kernel

In the asexual model, our results also give analytical insights on the effect of the shape of the mutation kernel on the adaptation to a changing environment. Empirical data on the exact distribution of mutational effects on phenotypic traits are hard to get (even though there is more data on the fitness effects of mutations) (see e.g. Halligan and Keightley, 2009; Nei, 2014). Most models therefore assume for mathematical convenience a Gaussian distribution of mutational effects. A few simulation studies have however explored marginally the consequences of a different, leptokurtic, mutation kernel (Keightley and Hill, 1988; Bürger, 1999; Waxman and Peck, 1999) : they found that a fatter tail for the distribution of mutational effects led to higher phenotypic variance, smaller evolutionary lag and greater fitness. The present analytical results are consistent with these past simulation results and show that we may expect in general distributions of mutations with higher kurtosis to reduce maladaptation and improve fitness, especially in fast changing environments.

### The advantage of sex in changing environments

Previous studies (Charlesworth, 1993; Bürger, 1999; Waxman and Peck, 1999) have used the Gaussian assumption and/or simulations to compare the dynamics of adaptation to a changing environment in sexual and asexual organisms. They all reached the conclusion that sex should provide a net advantage in a directionally changing environment, with a lower lag and greater fitness, which was ultimately due to the greater phenotypic variance evolving in a sexually reproducing populations. More precisely, Bürger (1999) and Waxman and Peck (1999) found that the phenotypic variance in sexual organisms would increase significantly with the speed of environmental change, while it would have only moderate effects on the variance in the asexual population. These findings seem to contrast with our comparison of the asexual model and sexual infinitesimal model, with more constraints on the evolution of the phenotypic variance for the latter. However, we would warn against interpreting our comparison of the infinitesimal and asexual model as informing about the advantage of sex in a changing environment. We rather see this comparison as informing us about the consequences of some modeling choices, with various constraints on the evolution of the phenotypic variance. First, for the ease of comparison between models, we used the same notation **V**_div_ to determine the amount of new variation introduced through reproduction in the progeny of parents in both models: in the asexual model it describes the amount of variance introduced by mutation, while it describes variation due to segregation in the infinitesimal model. It is unclear whether these quantities would be comparable with an explicit genetic model, including mutation and segregation at a finite set of loci. Second, we note that both Bürger (1999) and Waxman and Peck (1999) used in their simulations parameter values for mutation and selection corresponding well to the regime of the House-of-Cards approximation (Turelli, 1984; Turelli and Barton, 1990; Bürger, 2000), with rare mutations of large effects on fitness. Our study focused on a different regime of frequent mutations with small effects. Even if the equilibrium variance is small in both cases, the effect of a changing environment is different.

### Conclusions and perspectives

One of the main conclusion of our study is that the phenotypic variance at equilibrium truly depends on the modelling choice of the mode of reproduction. To understand this relationship, the approximation of the phenotype distribution appeared necessary. This approach is indeed robust, as shown by several studies following the same methodology in spatial structured population models: discrete patches ((Mirrahimi, 2017) with an asexual model and (Dekens, 2020) with the infinitesimal sexual model); dispersal evolution ((Perthame and Souganidis, 2016; Lam and Lou, 2017; Lam, 2017; W Hao, 2021; Calvez et al., 2022a; Lam et al., 2022) in the asexual case and (Dekens and Lavigne, 2021) in the infinitesimal sexual case). Moreover, this methodology is expected to be efficient to investigate other structured population models. Our next step will be to study the adaptation of an age–structured population to a changing environment, following (Cotto and Ronce, 2014). Other modes of reproduction with a more complicated genetic underlying architecture are also under investigation, (see for instance Dekens and Mirrahimi, 2021; Dekens et al., 2021).

## Supplementary Information

The following subsections gather mathematical analysis supporting the dimensionless scaling, numerical methods, Taylor expansions and formula derived in the main text. Although some parts are standard methods (rescaling, numerics), some parts are original contributions (dedicated Taylor expansions and formula involving the Lagrangian function), extending the literature in multiple ways. Hence, this supplementary material can be read as the companion mathematical paper of the main text.

Before we enter into the technical details, let us highlight some important observations about the Taylor expansions:

- These expansions are more than moment closure methods, where one usually tries to guess the higher moments of the distribution in order to derive a close system of equations on some scalar quantities (first moments of the distribution, *e.g*. population size, mean relative phenotype value, etc). Here, the *whole distribution* is approximated, then scalar quantities are deduced without any *a priori* assumptions on the shape of the distribution.
- In contrast to classical expansions of the distribution *F* which are *linear, e.g. F* = *F*_0_ + *εF*_1_ + …, we perform here a *multiplicative* Taylor expansion, meaning a linear expansion of the logarithm of the density: *U* = *U*_0_ + *εU*_1_ + …. We claim this is the natural expansion in the regime of small variance in order to discard the variance from the asymptotic calculations. Nonetheless, intermediate computations may appear heavy because of the nonlinear nature of the multiplicative expansion.
- We believe all these approximations can be theoretically justified, and error terms can be controlled quantitatively up to some extent. Results in the literature so far cover the case without environmental change (c = 0), see (Perthame and Barles, 2008; Barles et al., 2009; Mirrahimi and Raoul, 2013) for the asexual model, and the more recent (Calvez et al., 2019; Patout, 2020) for the infinitesimal sexual model.

### SI A. Derivation of generic formula (39)

Let us consider the equilibrium of our model:

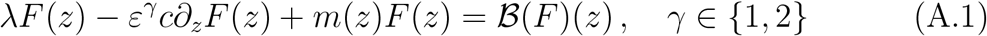

By integration over ℝ, we find:

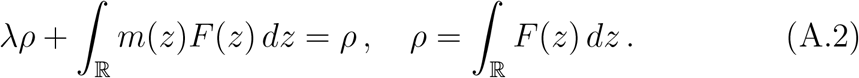

In the regime of small variance, we expect *F* to concentrate around the mean relative phenotype *z*\*, so as to get the following relationship

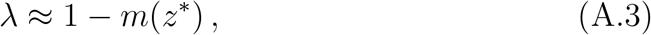

which corresponds to the demographic equilibrium. Next, we multiply by (*z* − *z*\*), where *z*\* is the mean value of the distribution *F*. Then, we integrate over ℝ to find:

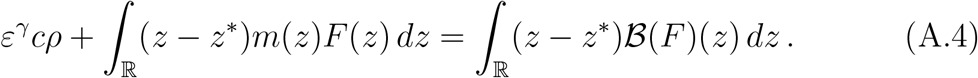

For any operator ℬ defined by (12) or (13), we find that the right-handside vanishes by definition of *z*\*. The concentration of the distribution *F* motivates the Taylor expansion of the selection function: *m*(*z*) ≈ *m*(*z*\*) + (*z* − *z*\*)*m*′(*z*\*) which implies the following:

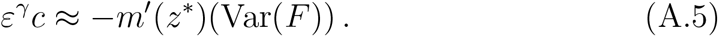

### SI B. Dimensionless scaling

We present in this section the details of the scaling procedure which leads to equations (14) and (15) in dimensionless form. By convention, the variables and parameters in original units are written in bold, whereas dimensionless quantity are in normal font.

The stationary state (***λ*, F**) satisfies

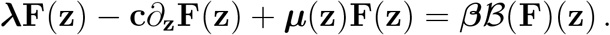

Dividing by the fecundity rate ***β***, (trait-independent) it becomes

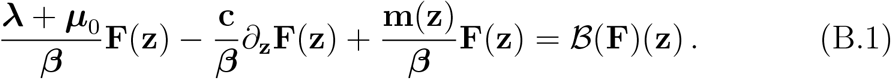

Around the optimum trait **z** = 0, the mortality per individual per generation **m**/***β*** is equivalent to

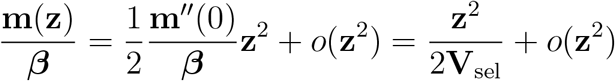

This, it is natural to measure traits at the selection scale:

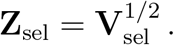

The mean fitness and the phenotypic distribution becomes in the scaled trait variable *z* = **z**/**Z**_sel_:

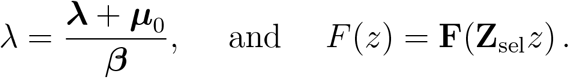

The mortality rate per individual becomes

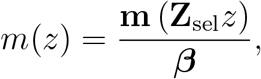

so that the selection strength around the optimum is scaled to a unit value:

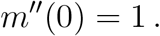

Our main assumption is that there is a small variability with respect to the selection scale **Z**_sel_. Denoting by **Z**_div_ the standard deviation of offspring traits from the parental traits, 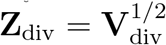, we define *ε* the scaling ratio:

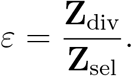

Then, our main assumption can be summarized as *ε* ≪ 1, paving the way to suitable Taylor expansions. Both models share the same notation for the standard deviation 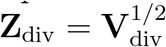 in the original units. However, we emphasize that it corresponds to mechanisms of variability associated with very different genetical background.

The reproduction operators ℬ are transformed as follows:

#### SI B.1. Asexual reproduction operator in scaled variables

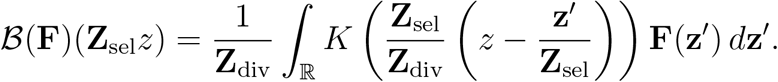

Using the change of variable *z*′ = **z**′/**Z**_sel_ in the integral and the definition of *ε* = **Z**_div_/**Z**_sel_, we obtain

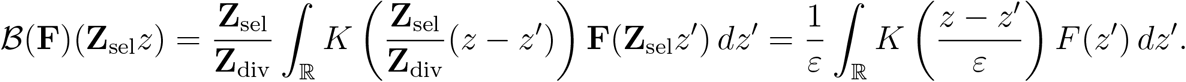

#### SI B.2. Sexual reproduction operator in scaled trait

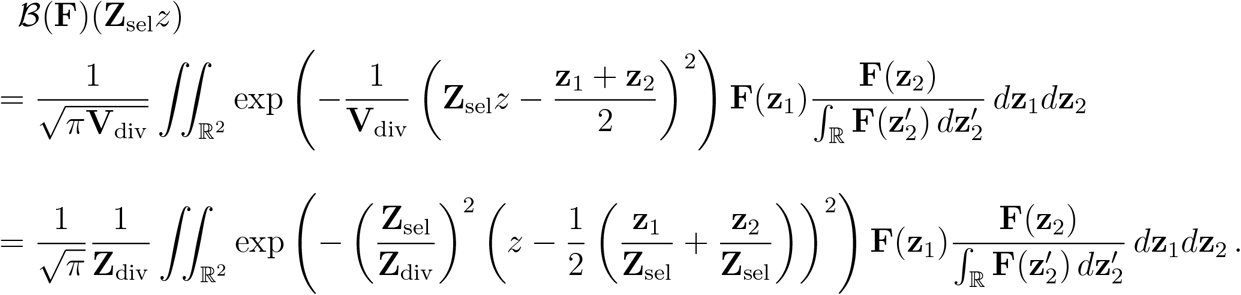

Using the change of variable *z*_1_ = **z**_1_/**Z**_sel_, *z*_2_ = **z**_2_/**Z**_sel_, and 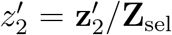, inthe integrals and the definition of *ε* = **Z**_div_/**Z**_sel_, we obtain

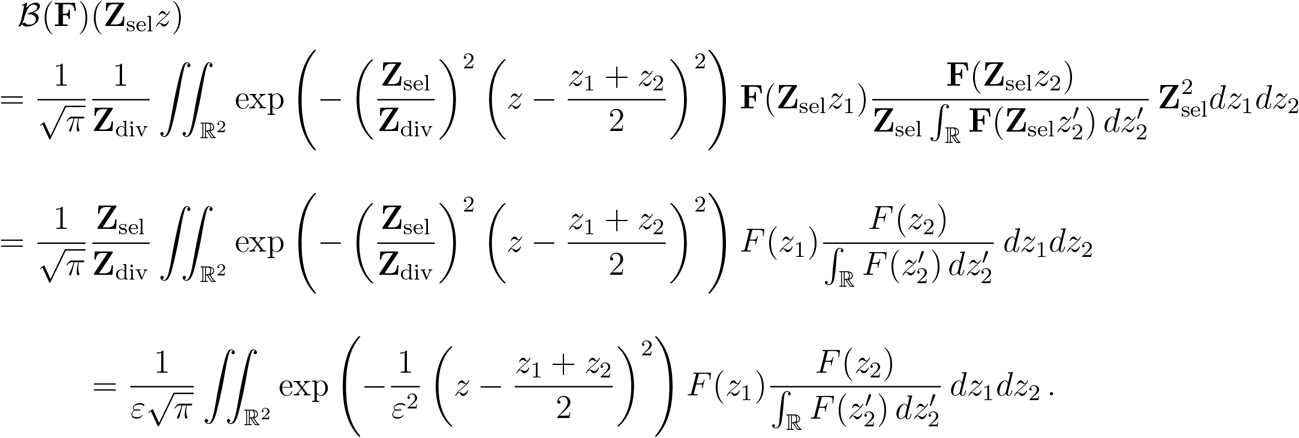

#### SI B.3. The dimensionless speed

It remains to express the dimensionless speed *c* = **c**/**C** with different choices of the typical speed **C**. This choice depends on the mode of reproduction as follows:

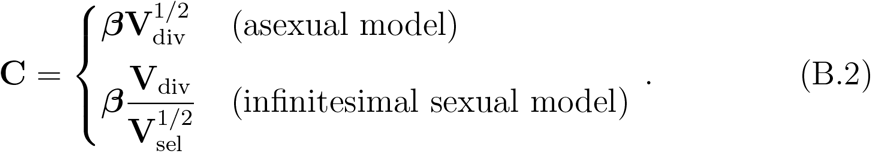

We thus deduce the dimensionless expression of the advection term:

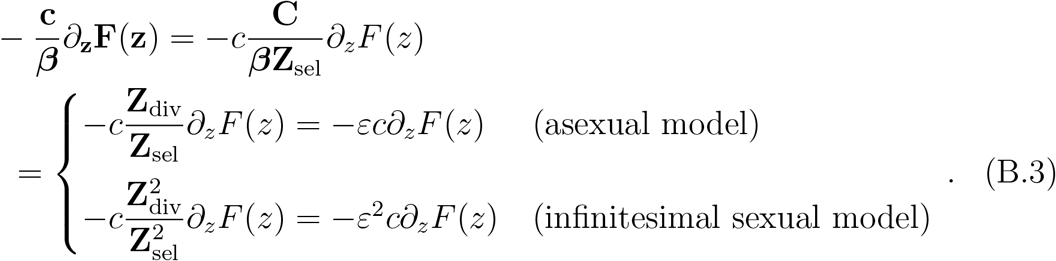

We obtain eventually the two rescaled problems as shown in (14) and (15). To conclude, let us mention that the discrepancy between the two values of **C** (B.2) is due to the very last step (B.3), where the dimensionless speed must be of order *ε* in the asexual model, resp. of order *ε*^2^ in the infinitesimal sexual model, in order to balance the other contributions. A mismatch at this step (*e.g*. any other power of *ε*) would result in a severe unbalance between the contributions, namely dramatic collapse of the population if the effective speed is too large, or no clear effect of the change if the effective speed is too small.

### SI C. Derivation of the variance

We compute below the formula of the phenotypic variance Var(*F*) in terms of *U* = −*ε*^*γ*^ log *F*,

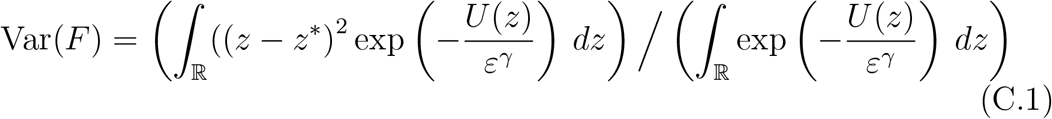

We assume that U reaches a non-degenerate minimum point at a unique z*, such that 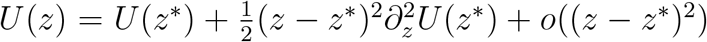 as *z* → *z*\*. The denominator is equivalent to

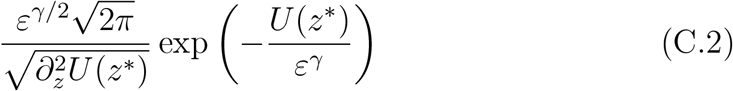

whereas the numerator is equivalent to

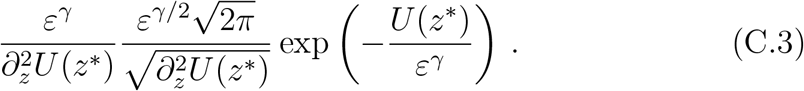

Thus, the ratio is equivalent to (18):

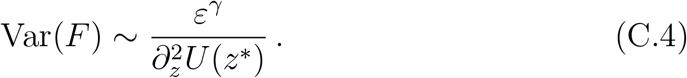

### SI D. Asexual type of reproduction (Details of Section 3.1)

This long section is devoted to the details of the Taylor expansion of *U* defined by (16). The equations verified by the successive terms *U*_0_ and *U*_1_ are derived. The meaningful formula are computed.

We can formally expand the pair (*λ, U*) with respect to *ε* as follows,

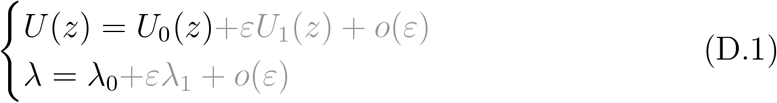

where (*λ*_0_, *U*_0_) gives the limit shape as *ε* →0, and (*λ*_1_, *U*_1_) is the correction for small *ε* > 0. We focus on the leading order contribution in this work. The corrector is required to refine our approximation in some part of the discussion.

#### SI D.1. Equations for (λ, U), (λ_0_, U_0_) and (λ_1_, U_1_)

We begin with the diffusion approximation for the sake of simplicity. This enables to present the main ingredient, namely *the completion of the square* in the equation, that will be generalized next for a general mutation kernel.

##### SI D.1.1. The diffusion approximation

The equation for *F* (14), together with the logarithmic transformation *F*(*z*) = exp(−*U*(*z*)*/ε*), is equivalent to the following one:

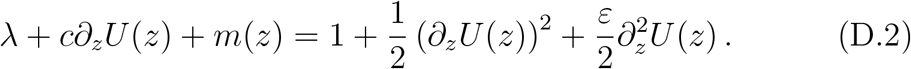

Clearly, the limiting problem for (*λ*_0_, *U*_0_) is

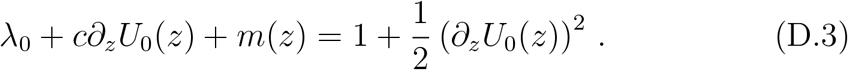

It is instructive to gather all the *∂*_*z*_*U*_0_ in the right hand side, then to complete the square:

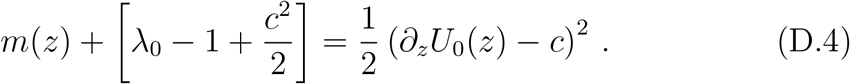

The key point is that there exist admissible solutions of this ODE if, and only if, the value between brackets vanishes, *i.e*. 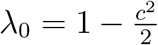. The argument is as follows.

###### Completion of the square

On the one hand, evaluating (D.4) at *z* = 0, we find that 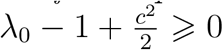 since *m*(0) = 0. On the other hand, if 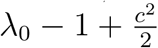 is positive, then *∂*_*z*_*U*_0_ − *c* does not change sign. Assuming without loss of generality that it is everywhere positive, we find that *U*_0_(*z*) ⩾ *cz* + *U*_0_(0) for *z* ⩾ 0 and *U*_0_(*z*) ⩽ *cz* + *U*_0_(0) for *z* ⩽ 0. In particular, we have *U*_0_(*z*) → −∞ as *z* → −∞, and *U*_0_(*z*) → +∞ as *z* → +∞, which is clearly not admissible because *F* is a population density. Therefore, 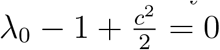.

Next, we can deduce the lag by evaluating (D.3) at 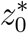 such that 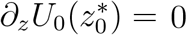,

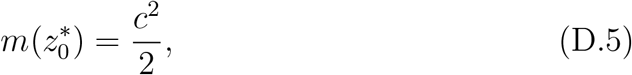

and also the value of the second derivative by differentiating once and evaluating at 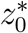:

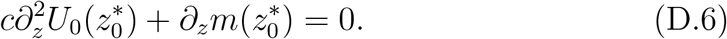

Finally, we deduce the variance from (18)

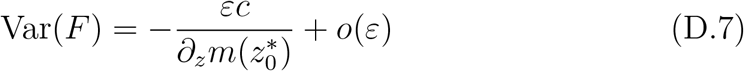

consistently with (39).

We can even provide a formula for the profile *U*_0_ by solving the ODE (D.4):

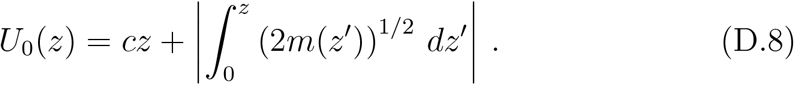

Notice that the environmental change acts here as a linear correction of the equilibrium profile obtained in the case *c* = 0. However, this is a peculiarity of the diffusion approximation.

It is another peculiarity that a quadratic selection function 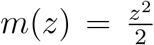 results in a quadratic profile 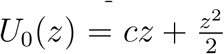 (D.8), which corresponds to a Gaussian distribution function *F* with variance *ε*.

##### SI D.1.2. The case of a mutation kernel

Again, we can reformulate the problem (14) in an equivalent form:

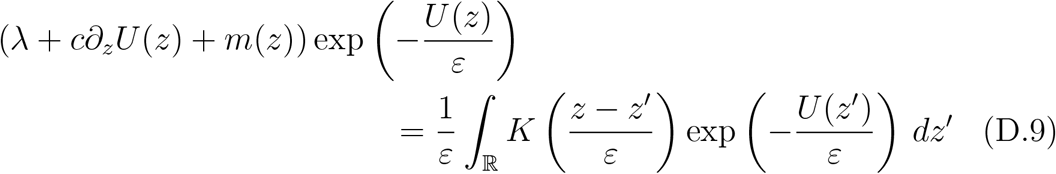

After the change of variables *z*′ = *z* − *εy* in the integral term, we obtain:

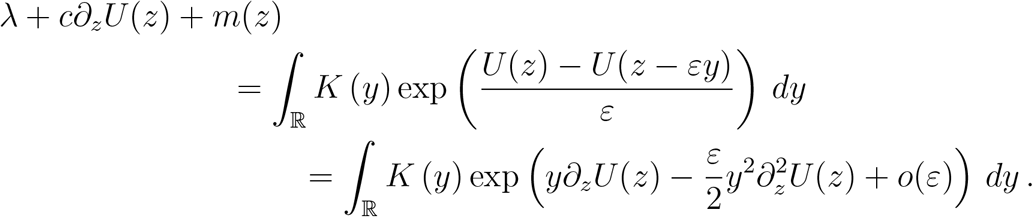

Injecting (D.1) into (14), but dropping terms of order higher than *ε*, we get

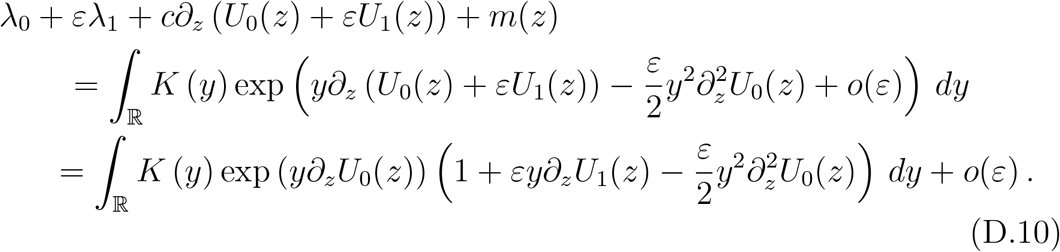

By identification of the contributions having the same order in *ε* in equation (D.10), we obtain the following equations for the pairs (*λ*_0_, *U*_0_) and (*λ*_1_, *U*_1_)

###### Limit problem

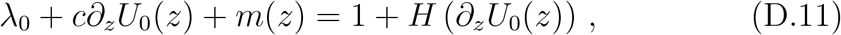

###### First correction problem

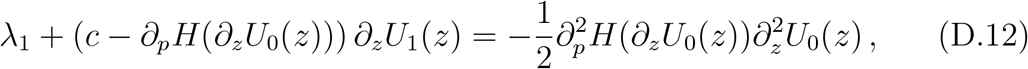

where the Hamiltonian function *H* is the two-sided Laplace transform of *K* up to an additive constant:

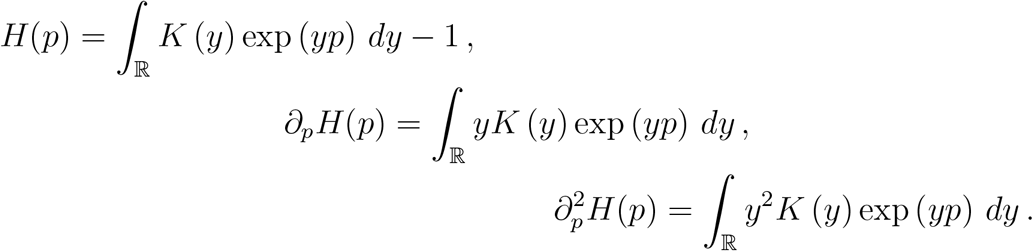

##### SI D.1.3. Computation of the mean fitness

The argument of Section SI D.1.1 for computing *λ*_0_ can be extended to the general case. Quadratic functions are replaced by convex ones, but the argument is essentially the same.

Again, let us reorganize (19) as follows, gathering the *∂*_*z*_*U*_0_ in the right hand side,

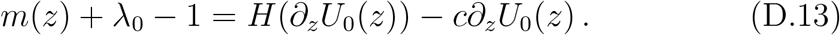

The function *p* ↦*cp* − *H*(*p*) reaches a maximum value, denoted as *L*(*c*) by definition (22). Adding this value on each side, we find

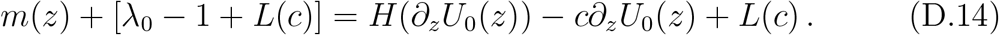

###### Completion of the generalized square

As in (D.4), the function *p* ↦*H*(*p*) − *cp* + *L*(*c*) in the right-hand-side is convex, nonnegative and touches zero. This is the analogous computation of *the completion of the square* by means of adding *L*(*c*). The same reasoning as above implies that the constant between brackets must vanish, *i.e λ*_0_ = 1 − *L*(*c*). Otherwise, the quantity *H*(*∂*_*z*_*U*_0_(*z*)) − *c∂*_*z*_*U*_0_(*z*) + *L*(*c*) would take positive values for *z* ∈ ℝ, hence the function *∂*_*z*_*U*_0_(*z*) could take values only on one of the two branches of the function *p* ↦ *H*(*p*) − *cp* + *L*(*c*), as depicted in Fig S1. As the function *p* ↦ *H*(*p*)−*cp*+*L*(*c*) is invertible on each separate branch, we could determine unambiguously the value of *∂*_*z*_*U*_0_(*z*) for *z* ∈ ℝ. In particular, it would have the same limiting value (possibly infinite) as *z* → −∞ and *z* → +∞ since *s*(−∞) = *s*(+∞). This would preclude the asymptotic behavior *U*_0_(*±*∞) = +∞ which is equivalent to vanishing population density at infinity. Hence, *λ*_0_ = 1 − *L*(*c*) is the only possible value.

#### SI D.2. Summary

So far we have obtained an analytical formula for the mean fitness,

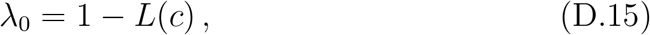

by means of the Lagrangian function which is the Legendre transform of the Hamiltonian function,

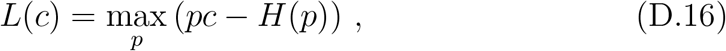

where *H* is the Laplace transform of the mutation kernel *K*.

**Figure S1:**
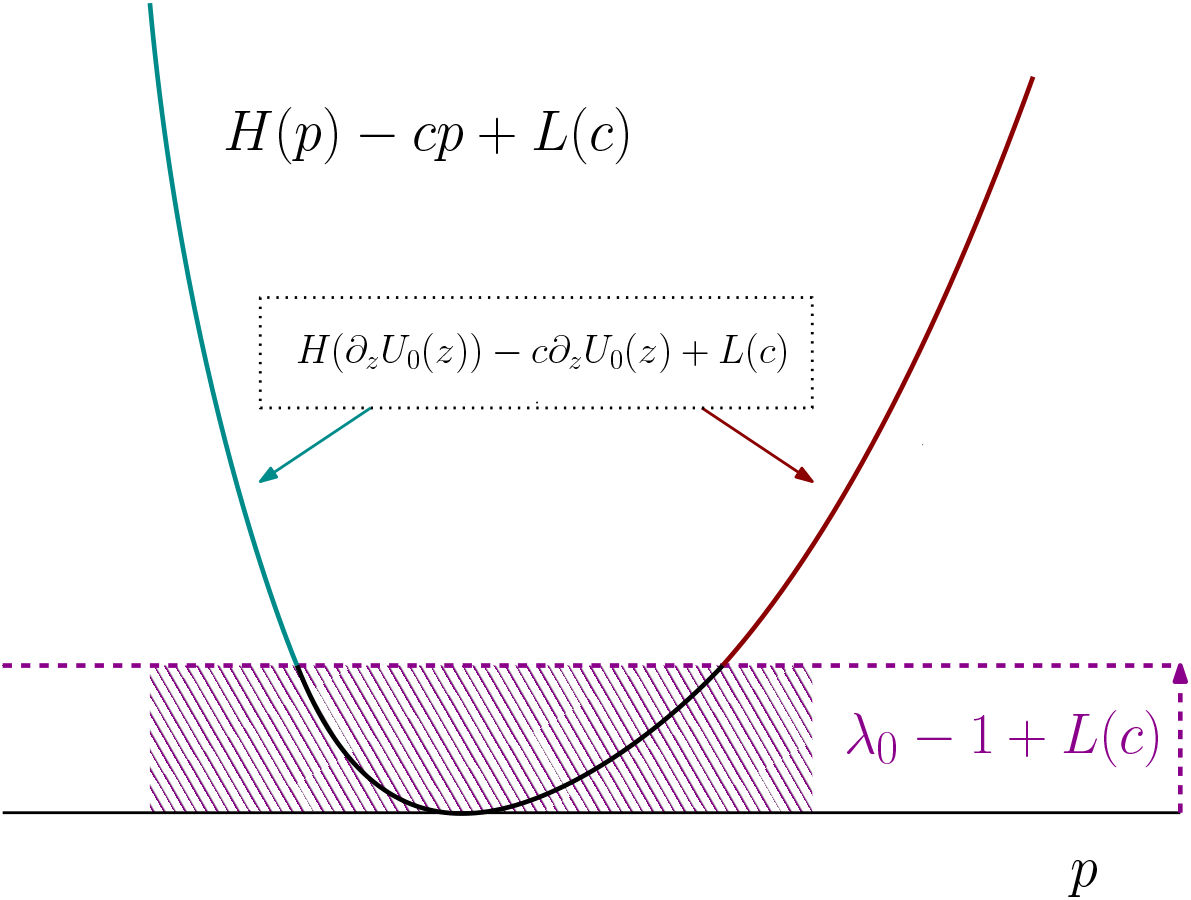
Sketch of the resolution of the main equation (19). The key function *p* ↦ *H*(*p*) − *cp* + *L*(*c*) is plotted. It is 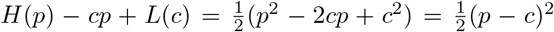 in the case of the diffusion approximation. More generally, it is always a convex function, with minimum value zero. The equation (19) can be reformulated as *H*(*p*_0_) − *cp*_0_ + *L*(*c*) = *m*(*z*) + [*λ*_0_ − 1 + *L*(*c*)], where the derivative *p*_0_ = *∂*_*z*_*U*_0_(*z*) must continuously change values as *z* goes from −∞ to +∞. In particular, it must have opposite signs at *z* = −∞ and *z* = + ∞, otherwise *U*_0_ would correspond to a non-admissible distribution *F* having an infinite limit on one side. A graphical analysis shows that it prescribes a unique value for *λ*_0_, that is *λ*_0_ = 1 − *L*(*c*). On the one side, evaluating at *z* = 0, we find [*λ*_0_ − 1 + *L*(*c*)] = *H*(*p*_0_) − *cp*_0_ + *L*(*c*) ⩾ 0 (by definition of *L*(*c*) which is the completion of the generalized square). On the other hand, we cannot have *λ*_0_ > 1 − *L*(*c*). If so, then we would get *H*(*p*_0_) − *cp*_0_ + *L*(*c*) ⩾ [*λ*_0_ − 1 + *L*(*c*)] > 0 for all values of the derivative *p*_0_. Thus, the solution would lie on one of the two branches of the function *p* ↦ *H*(*p*) − *cp* + *L*(*c*) (left or right), without possible continuous connection between the two. Consequently, the value *p*_0_ could be determined unambiguously by inverting *H*(*p*_0_) − *cp*_0_ + *L*(*c*) = *m*(*z*) + [*λ*_0_ − 1 + *L*(*c*)] on that branch for each *z* ∈ (−∞, +∞). This would induce the same limit for *p*_0_ as *z* → *±*∞, contradiction. Once the value of *λ*_0_ is found, it remains to solve *H*(*p*_0_) − *cp*_0_ + *L*(*c*) = *m*(*z*). This can be done in principle by inverting the function *p* ↦ *H*(*p*) − *cp* + *L*(*c*) for each *z*, with a careful choice of the branch. The switch between the two branches happens at (*z* = 0, *p*_0_ = *∂*_*c*_*L*(*c*)), where both functions *z* → *m*(*z*) and *p* ↦ *H*(*p*) − *cp* + *L*(*c*) reach their minimum value (zero).

The knowledge of the mean fitness enables deriving the lag load, which equilibrates birth and death in the population concentrated at trait 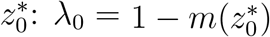, or equivalently

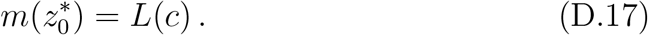

Note that the latter is equivalent to setting 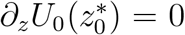 in (D.13) (critical point of the density), which is another characterization of the lag load.

The variance can be completed subsequently by differentiating (D.13) with respect to *z* and evaluating at 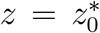. It is found that the variance equilibrates the fitness gradient and the speed of environmental change (*i.e*. the variations in the trait value in the moving frame):

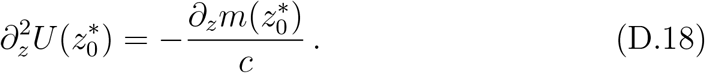

#### SI D.3. Conjugacy: Enlightening heuristics

There exists an alternative way to get some of the previous formula. The idea is to twist the unknown distribution *F* by a well chosen exponential function, in order to remove the transport part −*c∂*_*z*_*F* due to the environmental change. An enlightening example is the case of the diffusive approximation. Suppose the model is

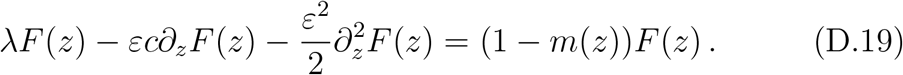

Then, the twisted distribution F(*z*) = *F*(*z*)*e*^*cz/ε*^ satisfies the following equation:

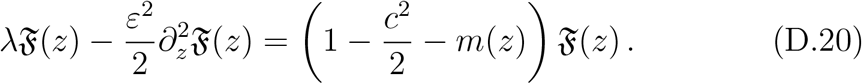

Therefore, we are reduced to a simpler problem without environmental change, at the expense of a global increase of mortality of value *c*^2^/2, consistently with the result of Section SI D.1.1.

However, the general case is based on heuristics rather than formal arguments. Starting from equation (14), or equivalently:

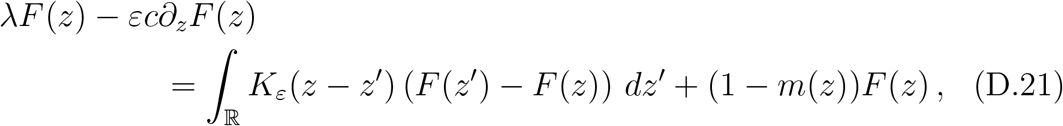

the density *F* is replaced with 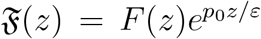, for some *p*_0_ ∈ ℝ to be characterized later on. The equation for 𝔉 is:

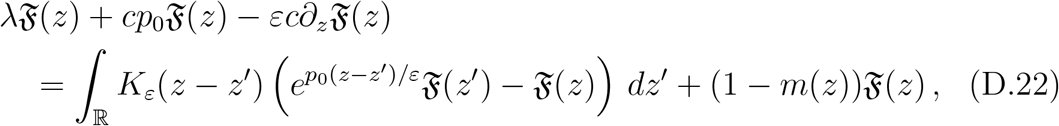

It is useful to rearrange the terms as follows:

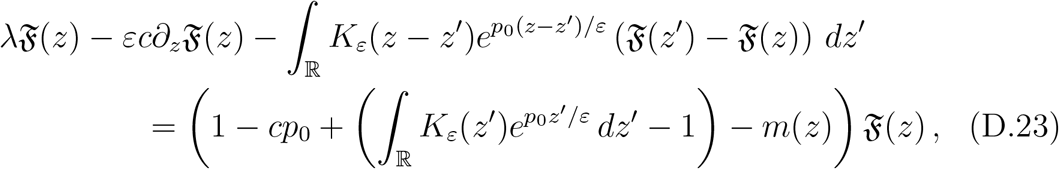

A natural way to choose *p*_0_ is to guarantee that the combination of transport and mutations preserves the center of mass of the distribution. This is a way to remove artificially the asymmetrical transport part. Thus, we propose the following characterization of *p*_0_: for any distribution 𝔉,

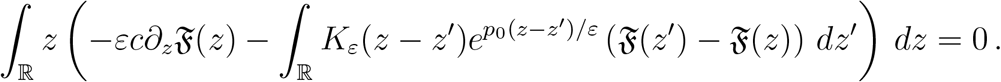

This is equivalent to:

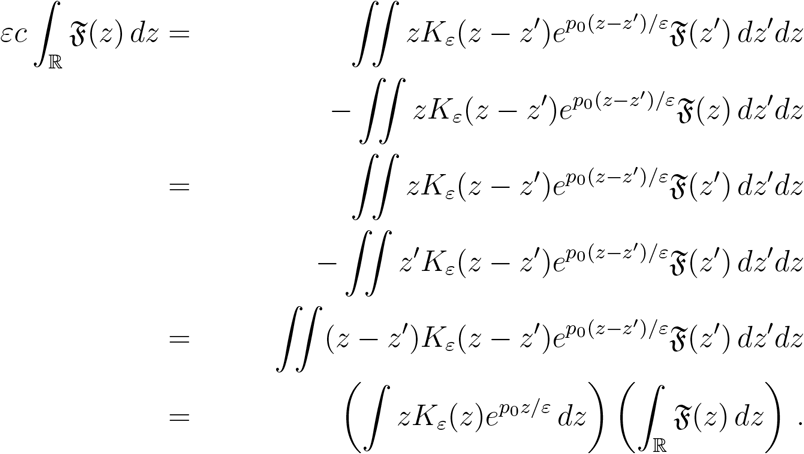

Finally, the required condition is equivalent to the following one, which appears to be independent of *ε* > 0:

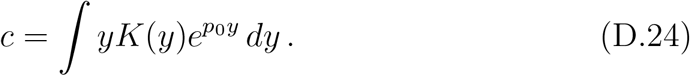

With the notations of Section SI D.1, this is also *c* = *∂*_*p*_*H*(*p*_0_). The right hand side of (D.23) becomes:

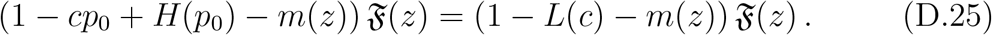

As a conclusion, we have shown that the combination of transport and mutations is equivalent to an operator which preserves the center of mass, up to a global increase of mortality of value *L*(*c*).

#### SI D.4. Some properties of the Hamiltonian and Lagrangian functions

We gather below some classical properties of the special functions that appeared useful in the analysis above.

##### The Hamiltonian and the mutation rate

The function *H* plays a pivotal role in our analysis. It could eventually break down if *H* degenerates. This would be the case, for instance, if the kernel *K* could be decomposed as **K** = (1 −*η*)*δ*_0_ + *η***K**_mut_, with small *η* ≪ 1. Indeed, it could be reformulated as follows

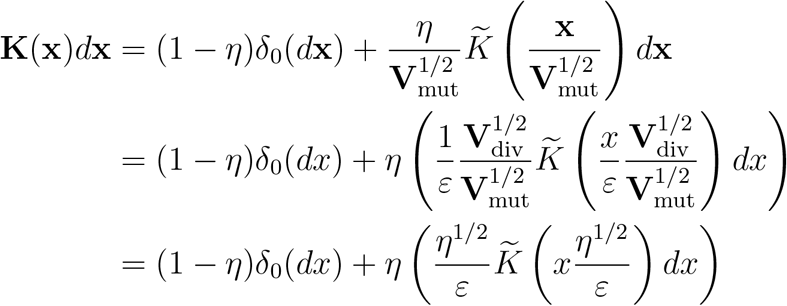

where we have used the relationship **V**_div_ = *η***V**_mut_ in the last line. Hence, the corresponding Hamiltonian function would be

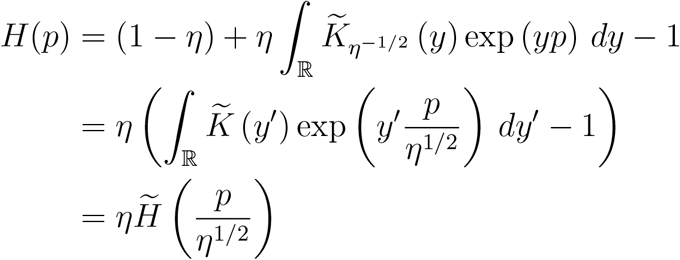

where 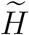 is the Laplace transform of the mutation kernel 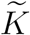. The latter expression would degenerate as *η* → 0, except if 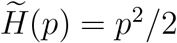.

##### Diffusion approximation as an extremal case of the convolution case

By symmetry of the kernel *K*, and its properties, the Hamiltonian function can be bounded below:

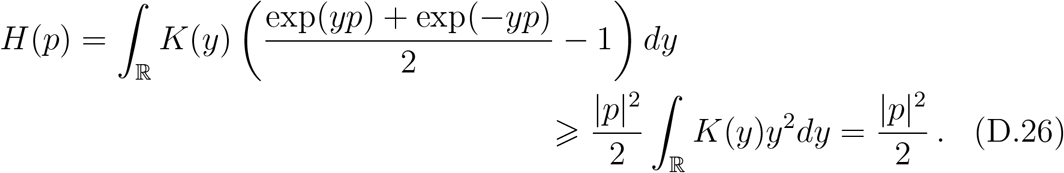

The latter expression is realized by the so-called diffusion approximation, see Section SI D.1.1. Indeed, the Hamiltonian function there was simply the square of the gradient (D.3). It is a direct consequence of the formula *L*(*c*) = max_*p*_ (*pc* − *H*(*p*)) (completion of the generalized square) that the Lagrangian function is bounded above:

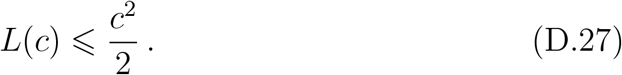

Hence, the maximum of lag load is realized for the diffusion approximation.

##### The Hamiltonian and the Lagrangian functions are dual from each other

The Hamiltonian *H*(*p*) can be recovered from the Lagrangian function *L*(*c*) by the very same formula, simply exchanging the roles of *c* and *p*:

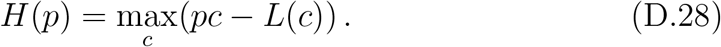

This inversion of the roles can also be seen on the derivatives of the functions, which are reciprocical one from each other. Indeed, at *p* = *p*_0_, we have *∂*_*p*_*H*(*p*_0_) = *c*_0_, where *c*_0_ is the one achieving the maximum value in (D.28), that is, the one satisfying the first order condition *p*_0_ = *∂*_*c*_*L*(*c*_0_). This is exactly the definition of reciprocical functions. This is a natural property from the viewpoint of convex analysis: the two functions *H* and *L* are indeed convex, so they have both monotonic derivative functions. The relationship between *H* and *L* is precisely the reciprocity of their derivatives.

##### The Hamiltonian function contains all the moments of the mutation kernel

By definition of the exponential function we have:

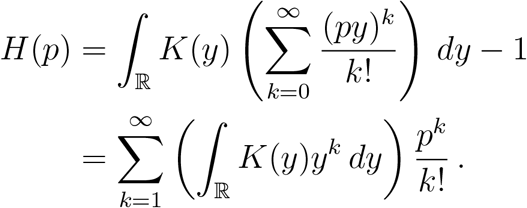

Hence, the moments of *K* are successive derivatives of *H* at the origin.

##### Influence of the kurtosis of the mutation kernel

As an immediate consequence, we see that the mean fitness *λ*_0_ = 1 −*L*(*c*) crucially depends on the full shape of the mutation kernel *K*. Indeed, the Lagrangian function *L* is related to the Laplace transform of the mutation kernel *K* (20) via the Legendre transform (22). To investigate this relationship, we investigate five kernels having the same variance, but different shapes, see Table D.4. We can show from the Taylor expansions that the Hamiltonian functions are ordered from top to bottom as follows:

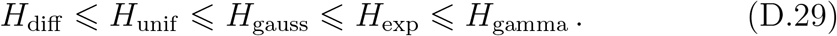

Accordingly, the Lagrangian functions are ordered in the opposite way, and the resulting mean fitnesses are ordered as follows:

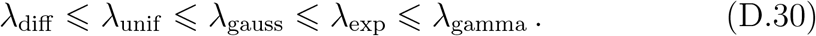

Hence, the lag load is ordered with respect to the kurtosis of the kernel.

#### SI D.5. Consistency of the formula for 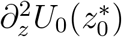 at c = 0

Here, we justify Remark 1, meaning that the formula obtained for 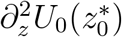 at *c* > 0 (D.18) coincides with the formula at *c* = 0, namely 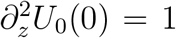. The latter is derived as follows. Firstly, the mean fitness (D.15) is *λ*_0_ = 1, as *L*(0) = 0, and the mean relative phenotype (D.17) is naturally 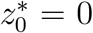 at *c* = 0 by definition of the mortality rate, optimum at the origin. Secondly, the expression of 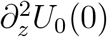 can be obtained by two alternative ways.

By differentiating twice (D.11) with respect to *z*, yields

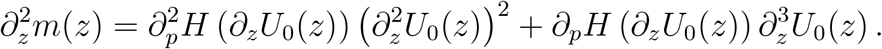

By evaluating this expression at *z* = 0, the last contribution vanishes because *∂*_*p*_*H*(*∂*_*z*_*U*_0_(0)) = *∂*_*p*_*H* (0) = 0. Hence, we get that

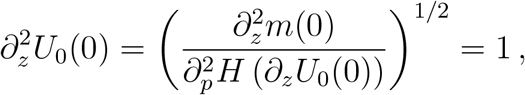

since 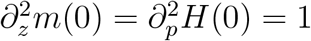.

**Table D.4:**
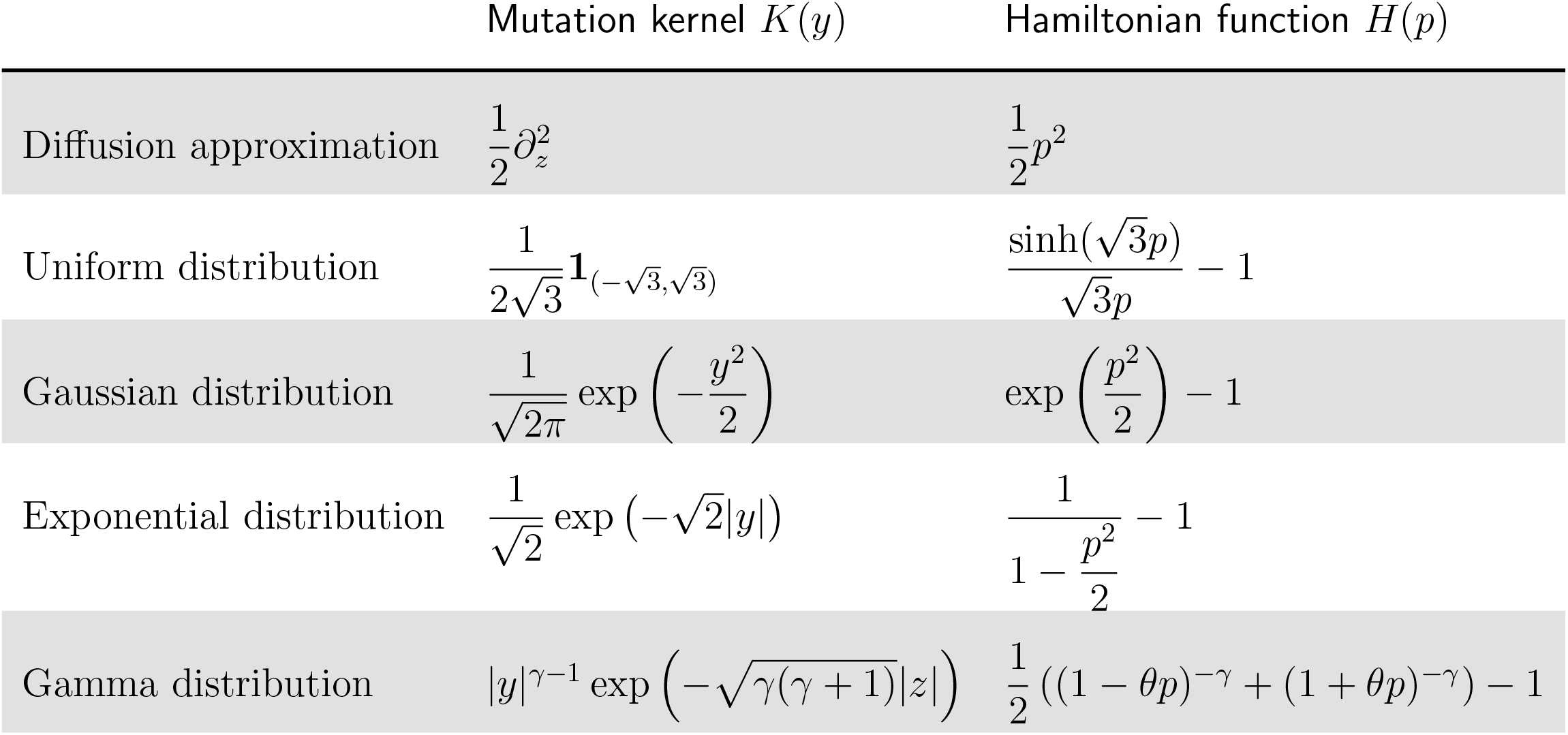
(Left) Five examples of mutation kernels with same (unit) variance, ordered by increasing kurtosis (from top to bottom). (Right) The associated Hamiltonian functions, with analytical formula. The corresponding Lagrangian functions cannot be expressed with classical functions, but the first one, up to our knowledge.

Alternatively, performing suitable Taylor expansions in expressions of, respectively, 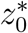 (24) and 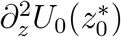 (25), as *c* → 0, yields:

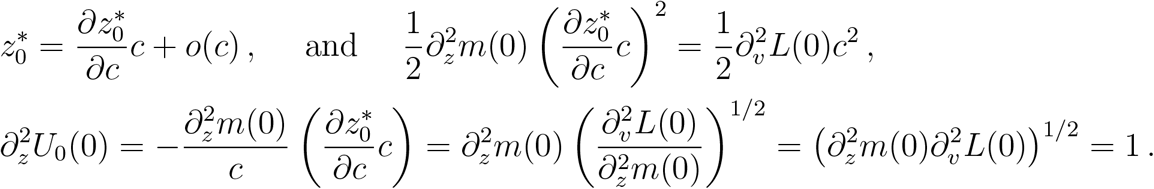

By reciprocity of the derivatives of *H* and *L*, we have 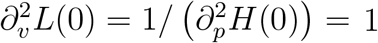. Both calculations coincide.

#### SI D.6. Quantitative description of the first correction (λ_1_, U_1_)

We derive useful informations from the equation (D.12) about the pair (*λ*_1_, *U*_1_). The methodology goes as in Section SI D.1.

We give the formula for the correctors *λ*_1_, 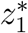, and the local shape around the minimal value: 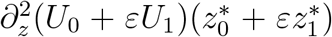. However, only the former one (*λ*_1_) is meant to be used in the main text, as it contains useful information about the mutation load in the population.

The formula are summarized in the following list, which completes those obtained in Section (SI D.2) at the leading order:

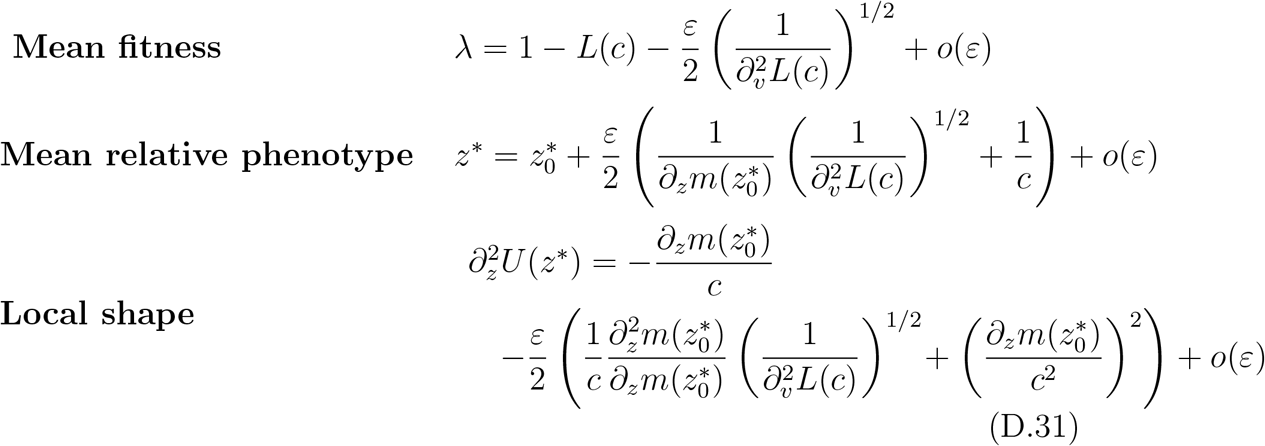

##### Description of the Mean fitness λ_1_

The equation (D.12) evaluated at the optimal trait *z* = 0 yields 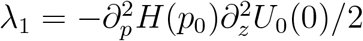, where *p*_0_ = *∂*_*z*_*U*_0_(0). To compute 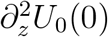, we differentiate (D.11) twice, and evaluate the expression at *z* = 0:

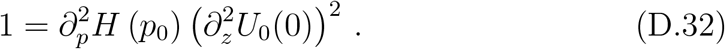

Recall that *p*_0_ = *∂*_*v*_*L*(*c*). Moreover, since *∂*_*p*_*H* and *∂*_*v*_*L* are reciprocal functions, then the second derivatives are inverse from each other. Therefore 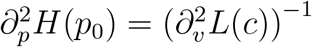. Thus, *λ*_1_ is given by the following expression:

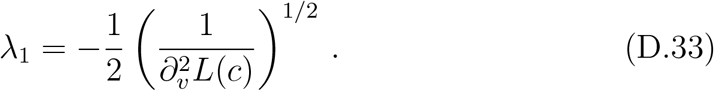

##### Description of the mean relative phenotype 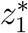

By pushing the computations further, it is also possible to derive the first order correction of the lag 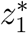. It is defined such that 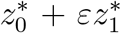 is the critical point of *U*_0_ + *εU*_1_, that is 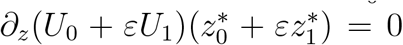. By expanding this relation, but keeping only the first order terms, we obtain 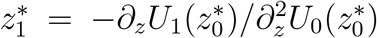. On the other hand, evaluating the equation (D.12) at 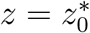 yields −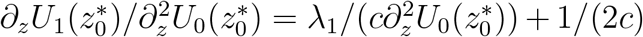. Using the expression (D.18) of 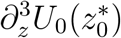, we obtain:

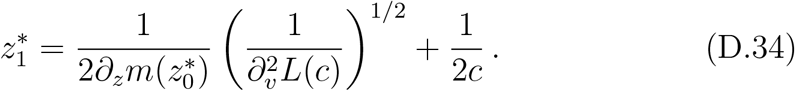

##### Description of the local shape

We expand the second derivative of *U*_0_ + *εU*_1_ at the lag point 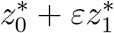 with respect to *ε* and we obtain

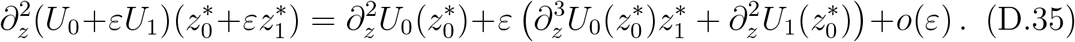

We aim at characterizing the term of order *ε* in this expansion. The first additional contribution 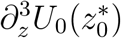 can be deduced from the equation (D.11) by differentiating it twice, and evaluating at 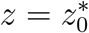:

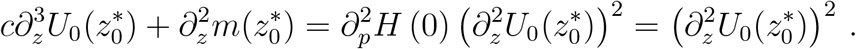

The second additional contribution 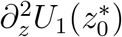 is deduced from the equation (D.12) by differentiating once and evaluating at 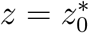:

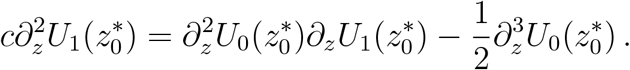

Combining these two expressions with the expression (D.34) of 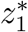, and 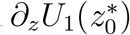, we get

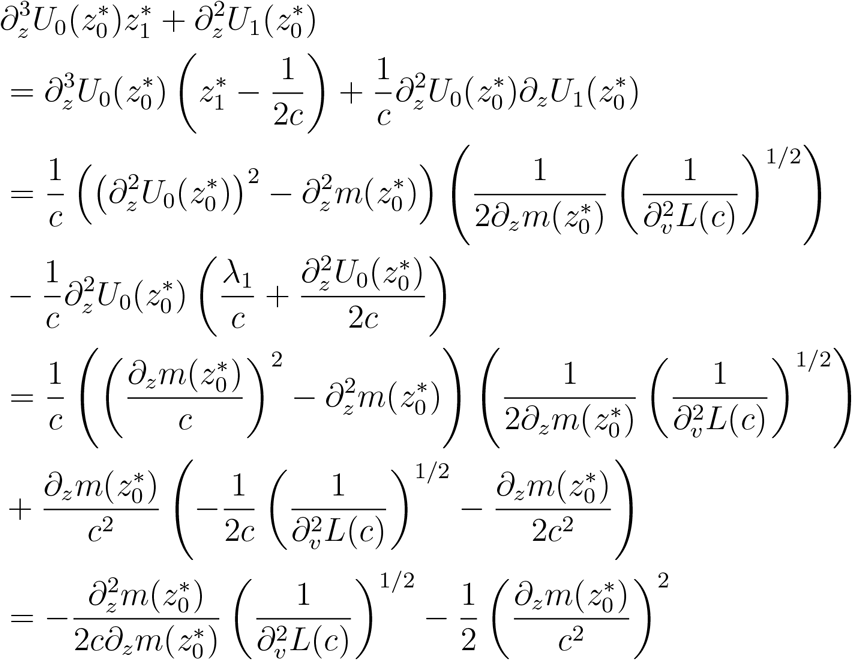

This concludes the analysis of the corrector problem at first order.

#### SI D.7. Numerical computation of the distributions U_0_ and U_1_ in the asexual model

The equation for *U*_0_ (D.11) is a non linear Ordinary Differential Equation (ODE). It has a singular point at *z* = 0, where the function *p* ↦ *cp* −*H*(*p*) cannot be inverted. It was solved numerically in the following way: after differentiation with respect to *z*, equation (D.11) becomes

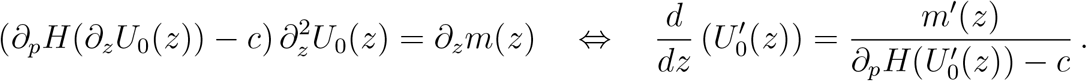

This ODE on 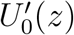 was solved using a classical solver (RK45), separately on the two branches *z* > 0 and *z* < 0. The issue is to initialize appropriately the solver for *z* = 0^+^, and *z* = 0^−^. The correct initialization was deduced from the analytical expressions of 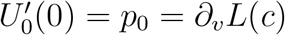.

Next, the linear ODE for *U*_1_ (D.12) was computed along characteristic lines:

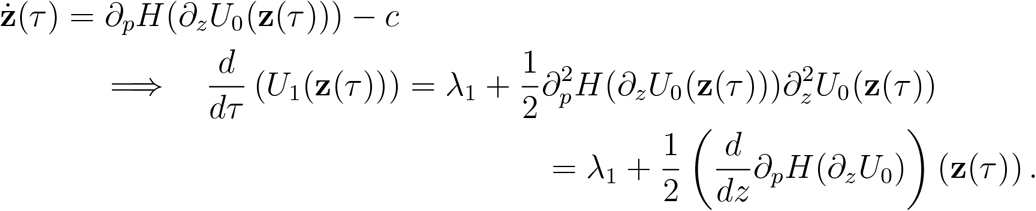

Integrating this formula with respect to time *τ* yields

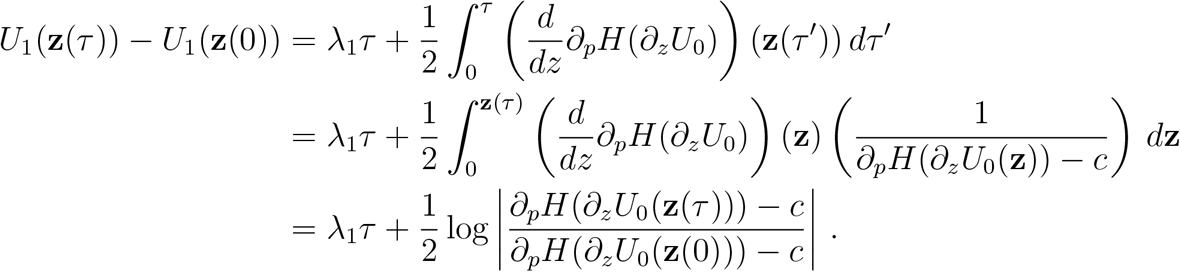

Again, the delicate issue is to evaluate appropriately the value *U*_1_(**z**(0)) for a starting point **z**(0) close to 0 (notice that 0 is an equilibrium point for the ODE: **ż**(*τ*) = *∂*_*p*_*H*(*∂*_*z*_*U*_0_(**z**(*τ*))) − *c*). The correct approximation is given by the analytical expression of *∂*_*z*_*U*_1_(0) obtained by differentiating equation (D.12) with respect to *z* and evaluating it at *z* = 0.

### SI E. Qualitative properties of the phenotypic variance at equilibrium Var(F)

In this section, we discuss in detail the behavior of the phenotypic variance at equilibrium with respect to the speed of change **c** in the scenario of asexual reproduction. Let us remind that in this case the phenotypic variance at equilibrium is well approximated by the following expression at the leading order:

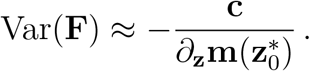

It is convenient to introduce the positive lag 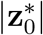, which is the distance to the optimal trait located at **z** = 0, so that

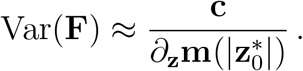

Recall that the lag is deduced from the inversion of the increment of mortality **m**:

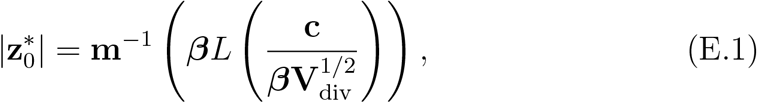

where **m**^−1^ is the inverse of the function **m** on (0, ∞). The differentiation of the lag 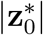 with respect to **c** goes as follows:

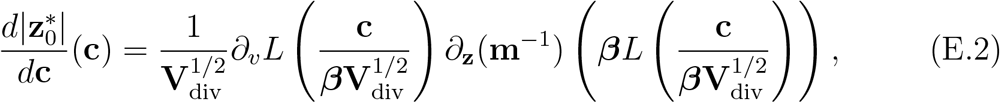

Since *∂*_**z**_(**m**^−1^) = 1*/∂*_**z**_**m**(**m**^−1^), the previous expression becomes

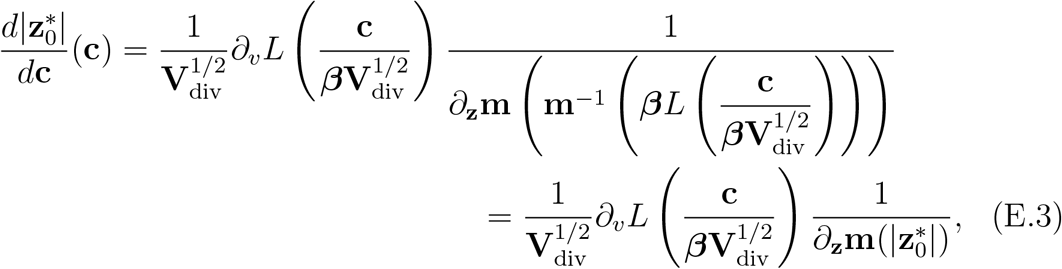

Reformulating this expression, we get an alternative expression for the variance:

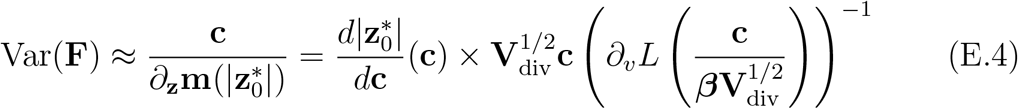

Now let us differentiate the latter expression with respect to **c**:

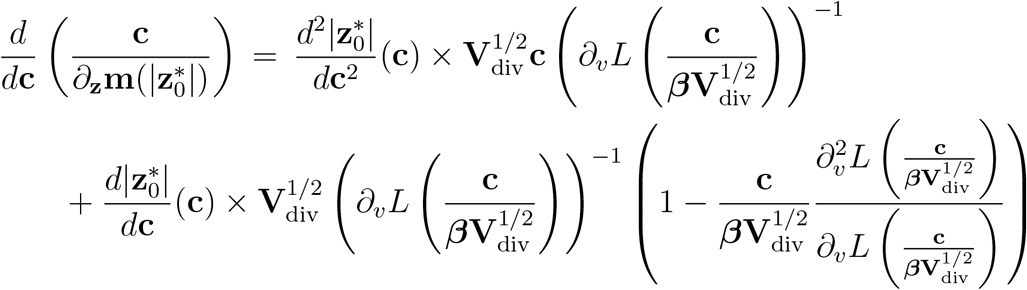

We shall establish that for all **c** > 0, the following inequality holds true:

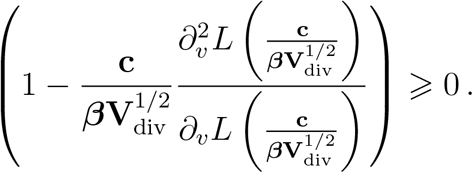

Indeed, it can be reformulated by means of *p* such that 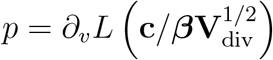, as follows:

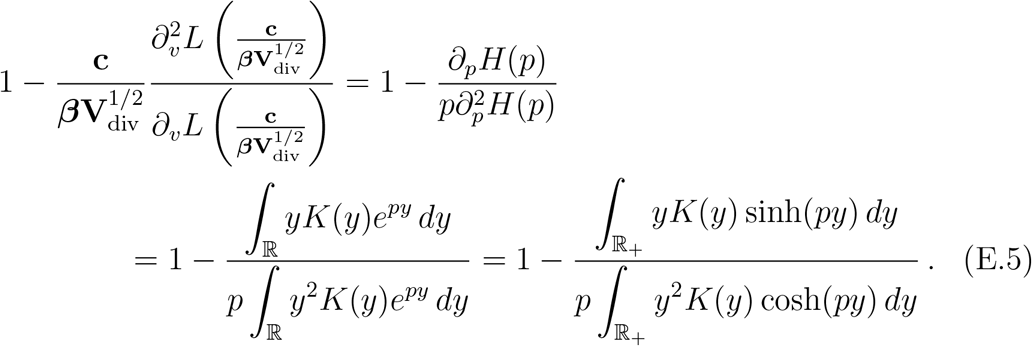

The conclusion follows from the pointwise inequality tanh(*py*) ⩽ *py* for *p, y* ⩾ 0, which is equivalent to sinh(*py*) ⩽ *py* cosh(*py*).

On the other hand, we have shown that the lag increases with respect to the speed of change **c**, thus 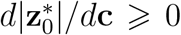. Then, if the lag is convex with respect to the speed of change **c**, that is 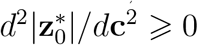, then the phenotypic variance at equilibrium increases with respect to the speed **c**.

However, the convexity of the lag depends on the convexity of the function *c* ↦*m*^−1^(*L*(*c*)). If the selection is quadratic *m*(*z*) = *z*^2^/2, this function is concave for any mutation kernel. However, if the selection function is more than quadratic, we can find mutation kernels such that the lag becomes convex.

In the diffusion approximation *L*(*c*) = *c*^2^/2, we can go further. In this case, we know from equation (33) that the lag accelerates with **c** if *m* is sub-quadratic. Whereas it is concave if *m* is super-quadratic in the sense of (34).

As a result, we have shown that the variance Var(**F**) increases with **c** if the function *c* ↦*m*^−1^(*L*(*c*)) is convex. More precisely, in the diffusion approximation, the variance increases with **c** if *m* is sub-quadratic in the sense of (33).

### SI F. Sexual type of reproduction (details of Section 3.2)

In this section we develop the computations required to describe *U* up to order *ε*^2^, as in (29). We present arguments from convex analysis to characterize *U*_0_. We provide an explicit formula for the first order correction *U*_1_ as an infinite series. Meanwhile, we present tedious computations needed to identify the linear part of *U*_1_, and we derive the first order correction of the mean fitness *λ*_1_ as a by-product.

Our starting point is the following relationship which is equivalent to finding a stationary density in the moving frame, expanded at first order in *ε*^2^:

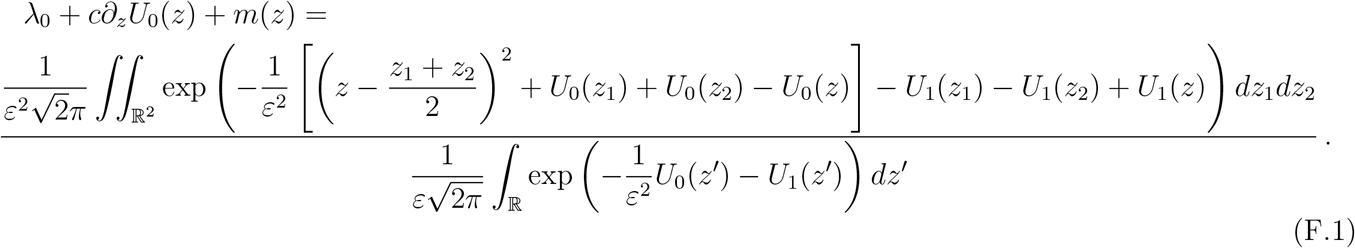

Note that the prefactors (involving *ε, π* have been arranged for the sake of normalizing singular integrals).

The arguments below are formal computations. We refer to (Calvez et al., 2019) for a rigorous analysis of this asymptotic analysis in the case *c* = 0, and to (Patout, 2020) for the time marching problem.

#### SI F.1. The characterization of U_0_ by convex analysis

Recall that the identity satisfied by *U*_0_ is the following one, ensuring that the right hand side of (F.1) does not get trivial as *ε* → 0:

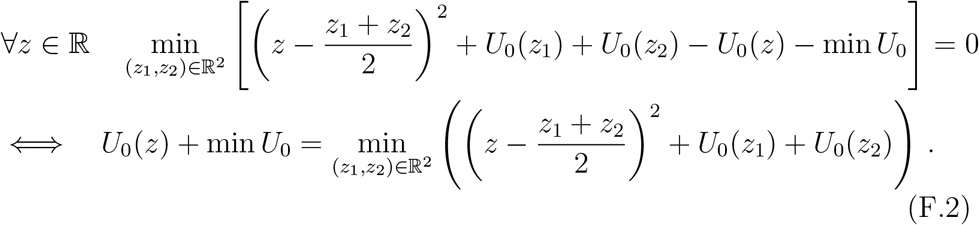

The goal of this section is to prove that any solution of the functional equation (F.2) is given by a member of the three parameters family

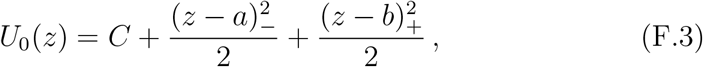

where the parameters *a, b* are such that *a* ⩽ *b* and *C* is an arbitrary constant. We denote by 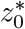 a minimum point of *U*_0_. We can restrict to min *U*_0_ = 0 without loss of generality (so that the additive constant *C* is set to 0). The characterization of *U*_0_ is done in several steps.

##### Regularity and −λ Concavity

Firstly, notice that *U*_0_(*z*) −*z*^2^ is a concave function, as it can be written as

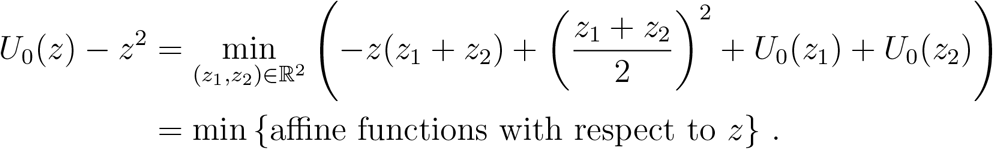

We deduce that *U*_0_ is continuous, and that it admits left and right derivatives everywhere.

##### The convex conjugate

The trick is to introduce the convex conjugate 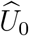 (also called the Legendre transform of *U*_0_):

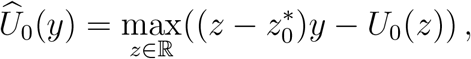

where 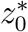 is a minimum point of *U*_0_. The basic properties of 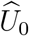 are listed below:

- 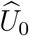 is convex, so it is continuous, and it admits left and right derivatives everywhere,
- 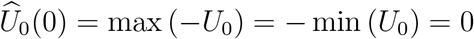,
- for all *y*, 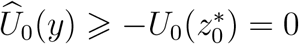, thus min 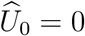.

We deduce from the functional identity (F.2), that

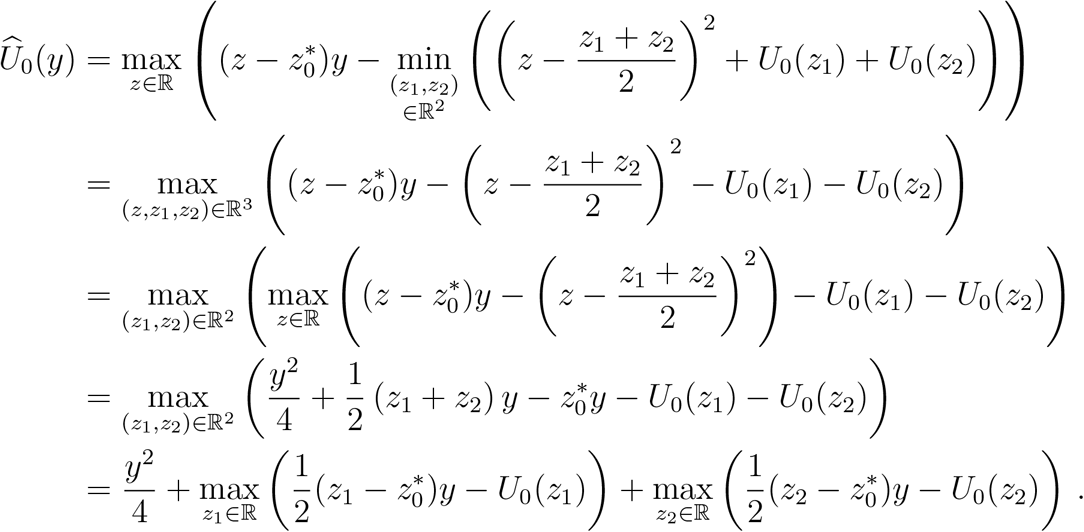

Finally, we end up with the following functional identity,

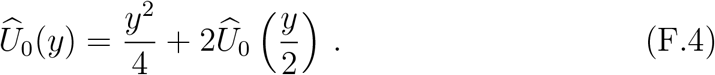

We observe that 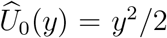 is a solution to the latter identity. However, it is not the only one. More generally, let 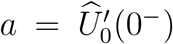 and 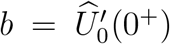 denote the left and the right derivative at *y* = 0, respectively. By convexity, and optimality at the origin *y* = 0 (namely, 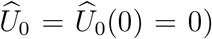), we have *a* ⩽ 0 ⩽ *b*. We deduce recursively from (F.4) the series expansion

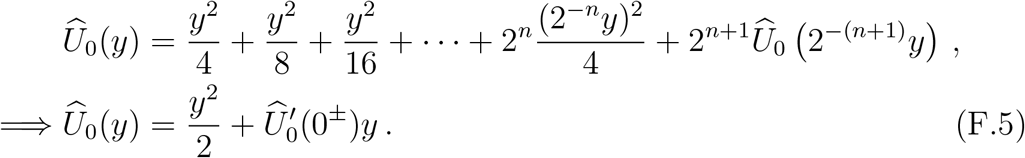

Obviously, the choice of the left or right derivative depends on the sign of *y*.

##### The convex bi-conjugate

Next, we define the convex bi-conjugate

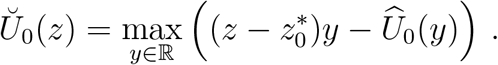

Standard results in convex analysis states that *Ŭ*_0_ and *U*_0_ coincide if *U*_0_ is convex. More generally, *Ŭ*_0_ is the (lower) convex envelope of *U*_0_ (Rockafellar, 1970). This is quite useful, because the characterization (F.5) enables to compute the convex bi-conjugate:

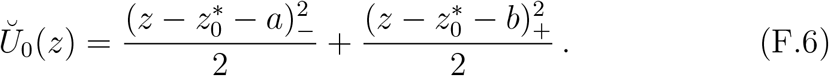

We deduce that the latter function is the (lower) convex envelope of *U*_0_. The last (delicate) step consists in proving that it coincides with *U*_0_.

##### From the convex envelope to the function

The idea is to use the functional identity (F.2) iteratively. As 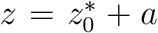 is an extremal point of the graph of *Ŭ*_0_, the values of *U*_0_ and *Ŭ*_0_ must coincide at this point. Hence, we have 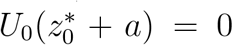, and similarly 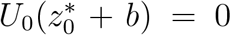. Recall that 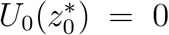 by definition. As a consequence, we have for 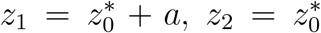, and 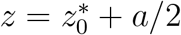 in (F.2):

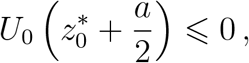

from which we deduce that *U*_0_ vanishes at 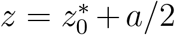 as well, and similarly at 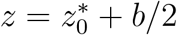. The same argument shows that *U*_0_ vanishes at each middle point between two vanishing points. So, it vanishes on a dense set of points in 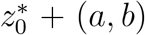. By continuity of *U*_0_, it vanishes everywhere on 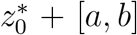. Finally, it coincides with its (lower) convex envelope (F.6) because the latter is strictly convex outside the interval [*a, b*].

Finally, it is necessary that *a* = *b* = 0 in the present context. Otherwise *F* would not correspond to a population density uniformly with respect to vanishing *ε*.

We have proved that *U*_0_ is necessary of the form

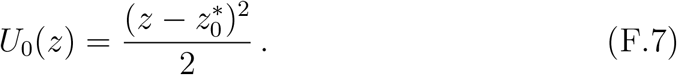

However, we are not able at this point to characterize the mean relative phenotype 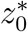. We need to push the analysis beyond the first order and compute the profile *U*_1_, as done in the following sections.

##### Discussion

There is an immediate interpretation of this result. We found that the equation (F.2) satisfied by *U*_0_ does not depend on the selection function *m*. Thus we can say that the main equation (F.1) is dominated by the reproduction term in the regime of small variance. Hence, the stationary distribution at the leading order equilibrium is the Gaussian distribution with prescribed variance (here, renormalized to a unit value), meaning a quadratic polynomial after taking the logarithm. In fact, Gaussian distributions are known to be stationary distributions of the Infinitesimal model in the absence of selection. As selection does not act on reproduction, there is no way to find the mean relative phenotype at equilibrium, and so 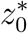 must be unknown at this point of analysis. The situation is quite different from the case of asexual reproduction, where no stationary distribution can be achieved without selection, and the mean relative phenotype is deduced from the knowledge of *U*_0_, accordingly.

#### SI F.2. Description of the corrector U_1_

Next, we can rearrange the right hand side in (F.1) using the characterization of *U*_0_ (F.7). It is instructive to begin with the denominator integral, which is a classical computation:

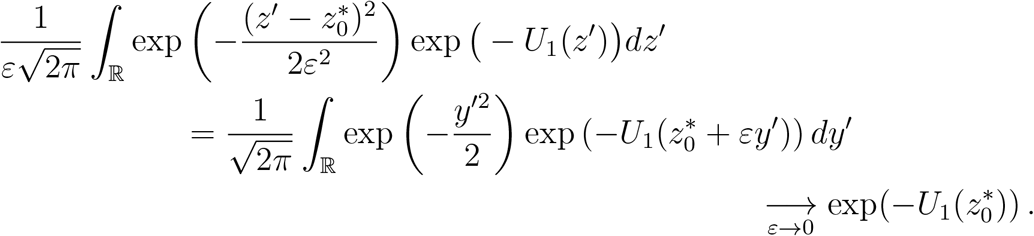

Indeed, the function 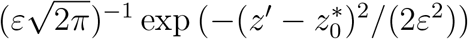 is the approximation of a Dirac mass as *ε* → 0. Hence the integral concentrates on the mean relative phenotype 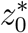: this yields the convergence of the integral towards 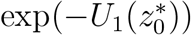. An alternative way to say is that, in the integral *F*(*z*′) *dz*′, most of the contribution comes from those *z*′ which are close to 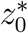.

##### SI F.2.1. What are the most representative parental trait values?

The same kind of computation allows handling the numerator in (F.1). The key point is to understand how the term inside the integral gets concentrated as *ε* → 0. In other words, we shall identify what are the most representative trait values (*z*_1_, *z*_2_) of parents giving birth to an offspring of trait *z*. Those will contribute mainly to the integral in the right hand side. They will enable to derive the equation for *U*_1_.

A preliminary computation is required: the double integral gets concentrated at the minimum points (with respect to variables (*z*_1_, *z*_2_)) of the quadratic form under brackets:

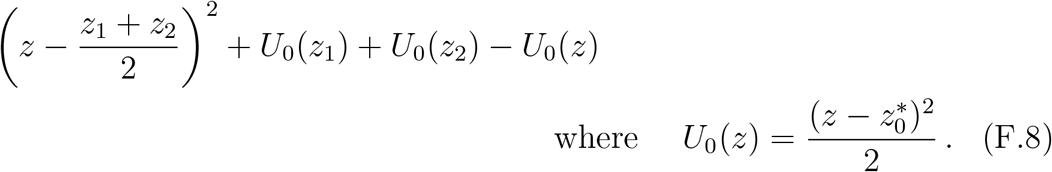

We know already that the minimum value is zero thanks to the characterization (F.2). The values above the minimum will contribute very little to the integral as they will have size of order exp(−*δ/ε*^2^), for *δ* > 0. Indeed, this decays to zero very fast as *ε* → 0.

Direct computation provides the unique minimum 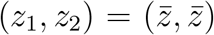, with 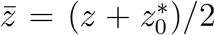. This means that an offspring of trait *z* is very likely to be the combination of equal parental trait values *z*_1_ = *z*_2_, equal to the mid-value between *z* and the mean relative phenotype 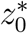. This is the result of an interesting trade-off: on one hand, parents with phenotype close to the mean relative phenotype value 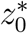 are more frequent; on the other hand, the chance of producing an offspring with phenotype *z* decreases when their own phenotype departs from the latter value. As a compromise, the most likely configuration is when both parents have the mid-point trait 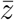, see Figure S2.

We thus define the following change of variable centered around this minimum point:

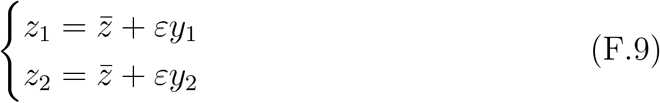

The quadratic form between brackets [⋯] in the numerator of (F.1) is transformed into an expression which does not depend on *ε*:

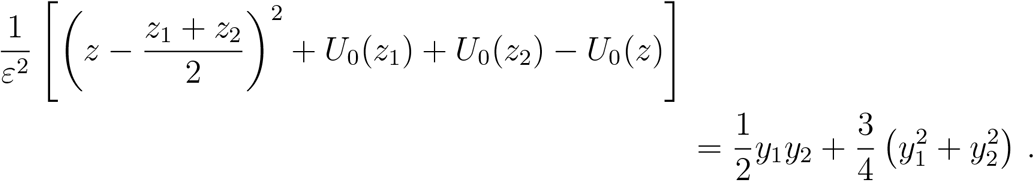

And the numerator finally writes

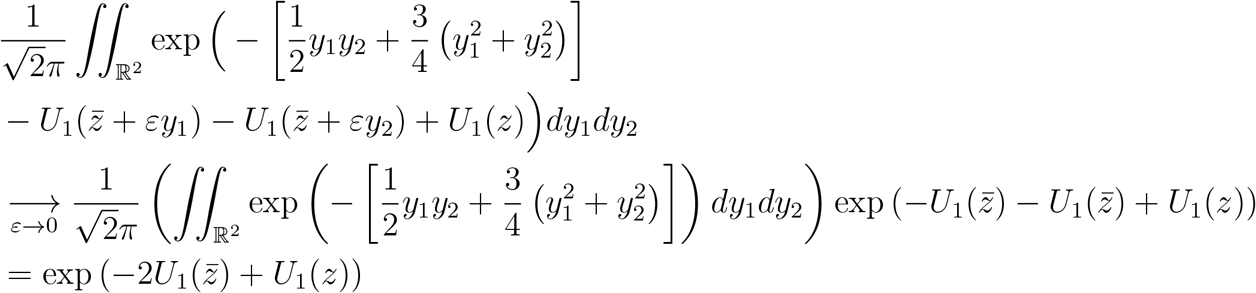

Note that the prefactor 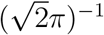 is such that the integral in (*y, y*) has unit value.

##### SI F.2.2. Equation for the corrector U_1_

We conclude that equation (F.1) converges as *ε* → 0 to the following equation on the corrector *U*_1_:

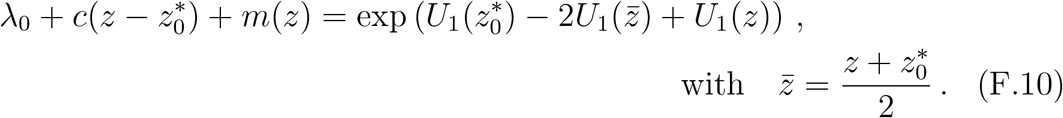

**Figure S2:**
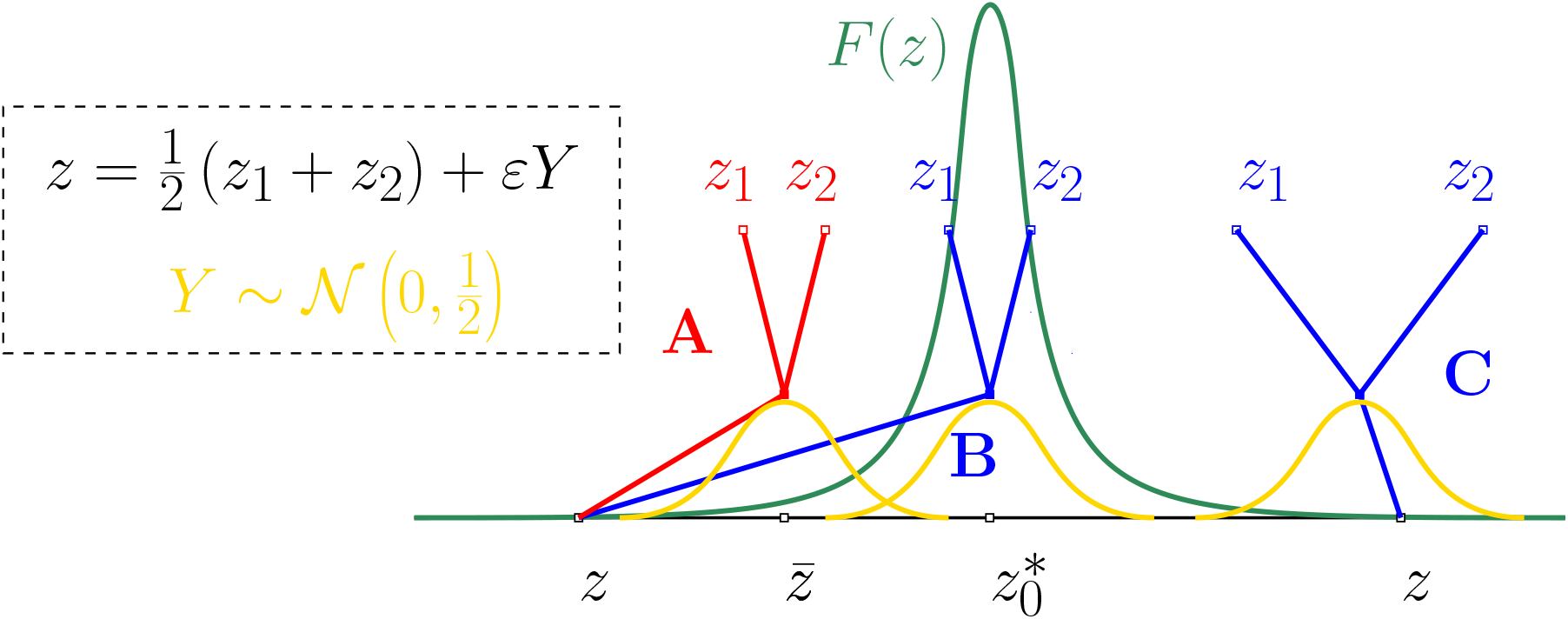
Sketch of the argument that underpins the estimation of the double integral in (F.1). Recall that the infinitesimal model assigns to an offspring the trait *z* which is the mean value of the parental trait values plus a normal random variable with standard deviation 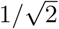 (in dimensionless variables). Among the three scenarios **A, B, C**, the first one is by far the most likely in the regime of small variance *ε*^2^ ≪ 1. In scenario **B**, the parental trait values (*z*_1_, *z*_2_) are close to the mean relative phenotype 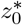: this is a likely event from the point of view of the parental trait distribution. However, it is very unlikely to draw a random number *Y* so large resulting in *z* at the next generation. In scenario **C**, the deviation is small, so that the mean parental trait is close to *z*: this is a likely event from the point of view of the “choice” of the offspring trait. However, it is very unlikely to draw a parent with trait *z*_2_ from the phenotypic distribution *F* : that one is too far from the mean relative phenotype in the tail of the distribution. Scenario **A** is the compromise between these two antagonistic effects.

This equation is simple enough to admit an explicit solution as an infinite series, as shown below.

Note that the values of *λ*_0_ and 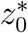 can be deduced readily from (F.11) as explained in the main text (32).

##### SI F.2.3. Analytical expression of U_1_

It is convenient to reformulate equation (F.10) as follows, by using the formula (32) for *λ*_0_ and 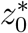,

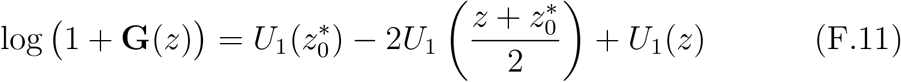

where 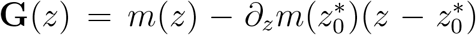 is such that **G**(0) = *∂*_*z*_**G**(0) = 0.

Differentiating this equation with respect to *z*, we obtain

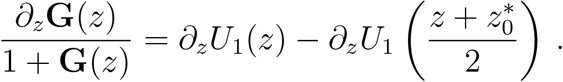

After the change of variable 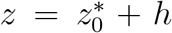, we get eventually the recursive relation where the value at some 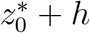 can be computed from the value at 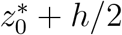,

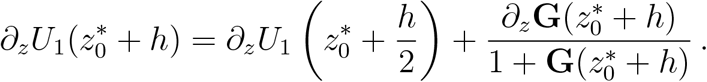

We deduce the following series expansion,

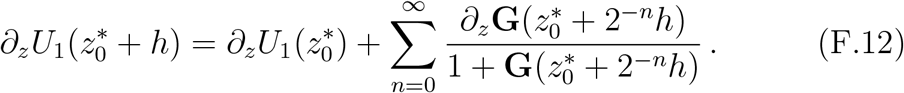

This provides an expression for *U*_1_ after integration with respect to *h*,

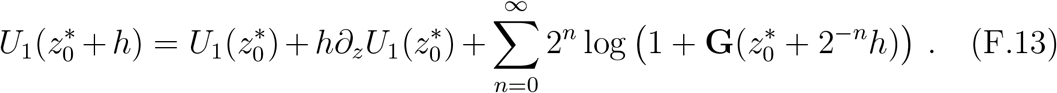

There are two degrees of freedom in the above expression of *U*_1_. First, the constant part 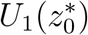 cannot be determined, because *U* is defined up to an additive constant. Thus, we are free to choose any value for 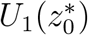, say 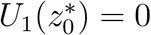 for instance. On the other hand, the value 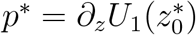 plays a key role in the shape of the distribution, related to the expansion of the mean relative phenotype, see (F.20) below, but its value cannot be elucidated at this stage. We need to push the expansion up to order *ε*^4^ to get the following formula for *p*\*:

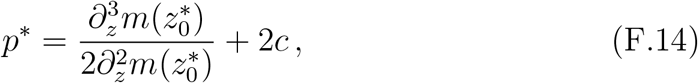

see next section for the complete computation (see also (Calvez et al., 2019) for an alternative path with limited expansions to the next order in the case *c* = 0).

We deduce the following expression for *U*_1_,

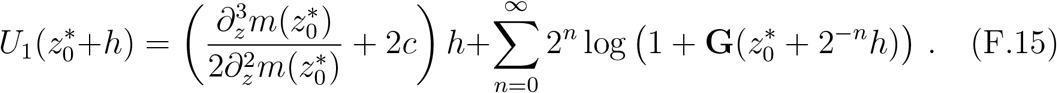

##### SI F.2.4. The missing linear part: calculation of 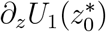

Starting with the equation satisfied by *U* (28), and plugging the ansatz

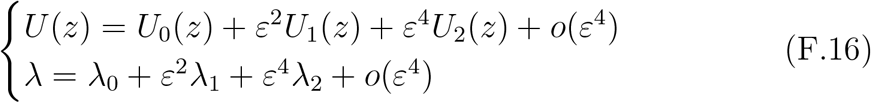

we obtain the following equation up to order *ε*^2^:

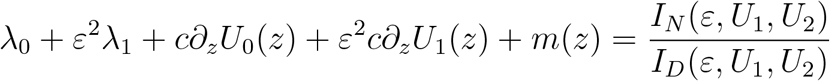

where

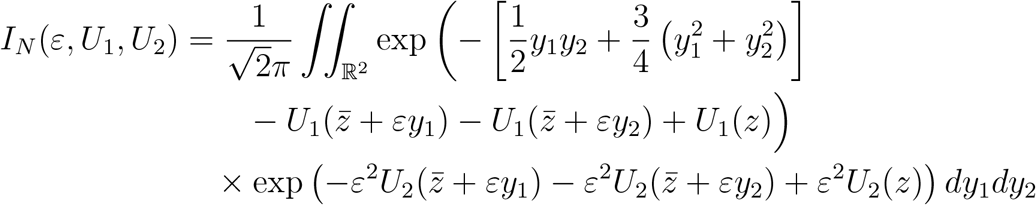

and

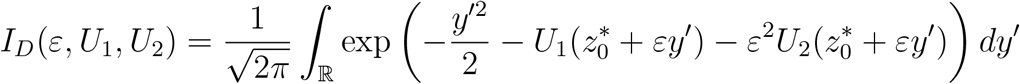

The integrals were subject to the same change of variables as in (F.9). After elimination of higher order contributions, we obtain for the denominator, up to order *ε*^2^:

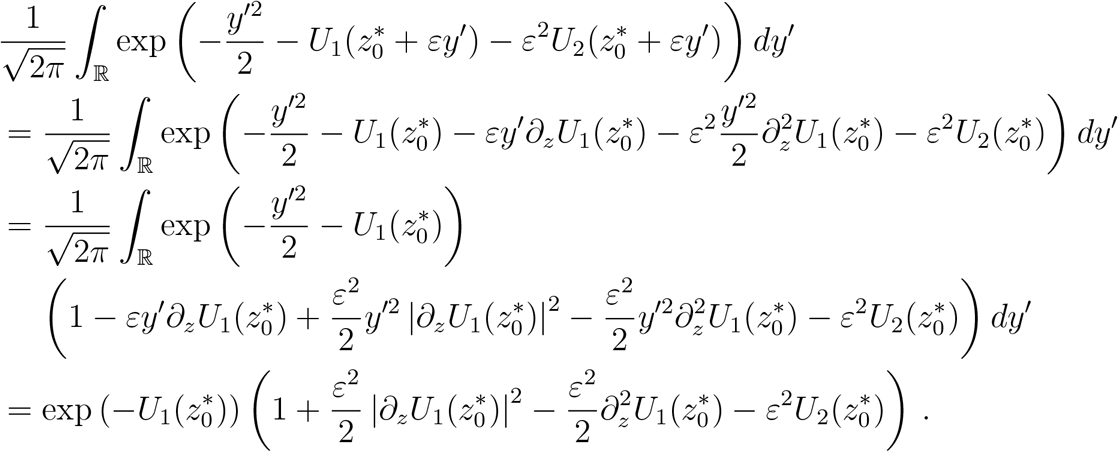

In an analogous way, we obtain for the numerator,

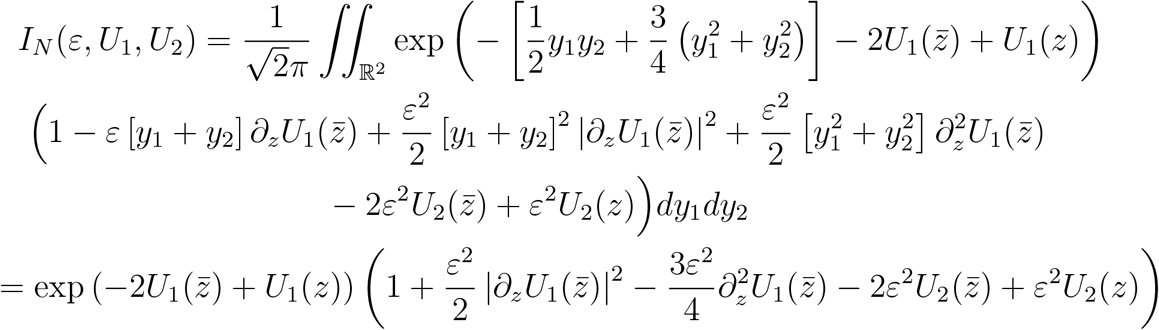

Combining all these expansions, we obtain up to order *ε*^2^:

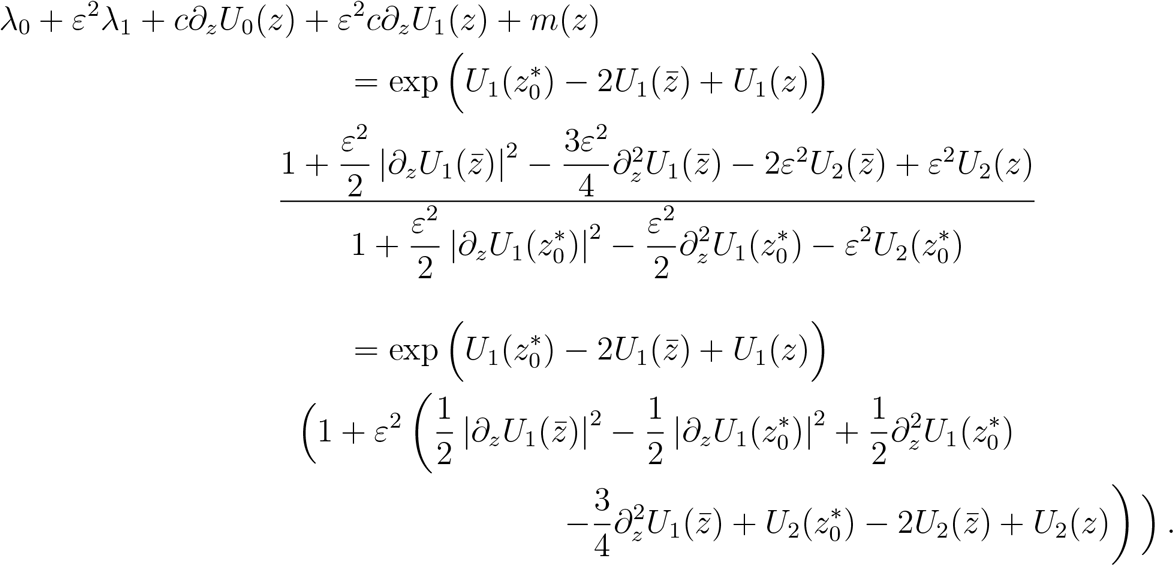

By identifying contributions of order *ε*^2^ on both sides, we deduce the following equation for the next order correction *U*_2_,

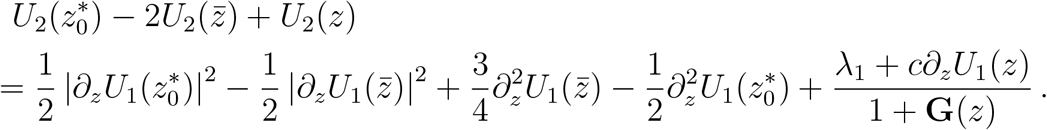

By evaluating, and differentiating at 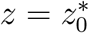, we deduce the following pair of identities,

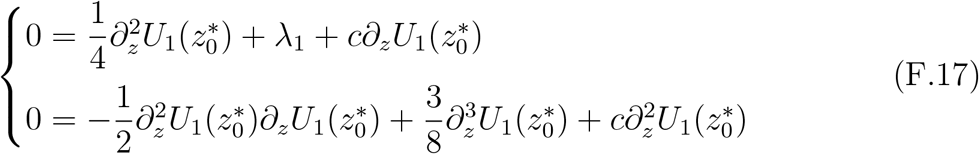

The second identity enables to compute 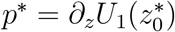:

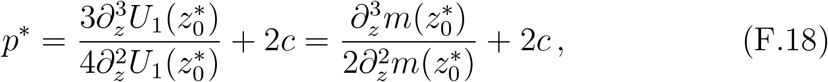

where 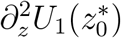 and 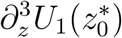 are deduced from equation (F.11) after multiple differentiation, or directly from (F.12). This yields the missing part in (F.15).

##### SI F.2.5. Analytical expressions of the macroscopic corrections terms λ_1_ and 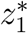

###### Description of Malthus rate λ_1_

The first identity in (F.18) provides 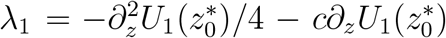. The expression (F.10) differentiated twice and evaluated at 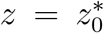, yields 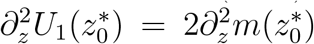. We conclude from the expression of *p*\* that

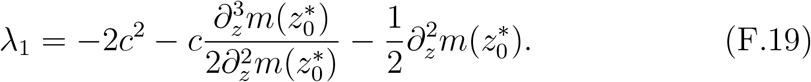

###### Description of the mean relative phenotype correction 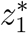

The first order correction of the mean relative phenotype 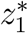 is defined such that 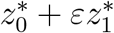 is the critical point of *U*_0_ + *ε*^2^*U*_1_, that is 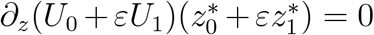. Expanding this relation and keeping only the terms of order *ε*^2^, we obtain using the expression of *p*\*,

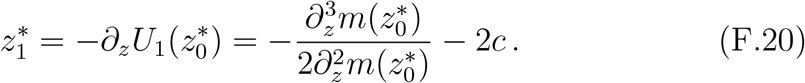

###### Description of the local shape

The second derivative of *U*_0_ + *ε*^2^*U*_1_ at the mean relative phenotype *z*\* is equal to 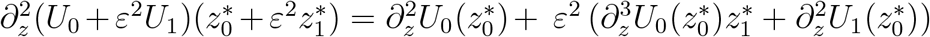, up to the order *ε*^2^. Since 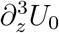 is equal to 0, we can deduce from the expression of *U*_1_ that the local shape around *z*\* is given by

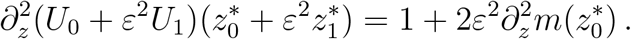

### SI G. Numerical computation of the equilibrium (*λ*, F)

In order to obtain numerical approximations of the pair (***λ*, F**), we get back to the time marching dynamics of the density **f** (**t, z**) which satisfies the following equation:

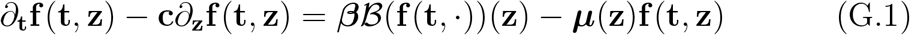

The density **f** (**t, z**) is expected to behave like exp(***λ*t**)**F**(**z**) for large time. It is preferable to introduce the frequency of traits in population: **p**(**t, z**) = **f** (**t, z**)/ ∫ **f** (**t, z**′) *d***z**′. The equation for **p** is:

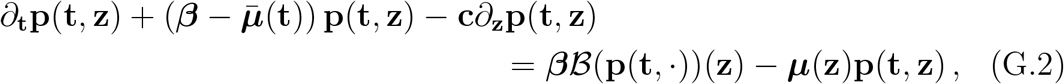

where the additional 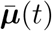 ensures that ∫ **p** remains constant:

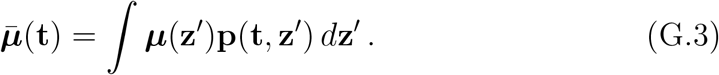

We expect that the pair 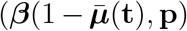 does converge to (***λ*, F**) as **t** →+ ∞.

Classical numerical methods were used to approximate (G.2)-(G.3) for large time, until some error threshold is reached for ∥*∂*_**t**_**p**(**t**, ·)∥_∞_. The transport term −**c***∂*_**z**_**p**(**t, z**) was handled using an upwind scheme. The convolutions involved in operator ℬ were handled using the function conv in MAT-LAB software. The grid mesh was adapted to the scales in SI SI B in order to capture the appropriate phenomena at the correct scale.

### SI H. Comparison with an Individual–based model

In this section we aim to compare our deterministic approximation with the outcome of a stochastic individual based model (IBM model) with a finite population. We first describe briefly the IBM model. Then we compare the equilibrium distribution of the IBM with our approximation distributions described in Fig. 8. Finally, we compare our results on the effect of the speed of environmental change with the outcomes of the IBM model.

#### Stochastic Individual Based Model

We consider a stochastic IBM model where each individual is characterized by its trait *X*_*i*_ and its size 1/**N**. They reproduce at a rate ***β*** and die at a rate that depends on their traits *X*_*i*_, the speed of environmental change **c** and on the number of individuals *N*_*t*_ in the population at time *t*. More precisely, the individuals may die due to their maladaptness in the phenotypic landscape, which happens at a rate ***μ***(*X*_*i*_ − **ct**). Or they may die from density dependence at a rate ***κ****N*_*t*_/**N**, where *N*_*t*_/**N** denotes the size of the population (it is similar to ***ρ***(*t*) in our deterministic model). The density dependence keeps the number of individuals finite, and proportional to **N**. In particular, when **N** tends to infinity, the number of individuals also tends to infinity and the (renormalized) stochastic model converges to the deterministic model (2) (Champagnat et al., 2006).

In the case of a birth event, the trait of the offspring is drawn according to the operator ℬ. In the asexual model, the offspring trait *X*_offspring_ is given by

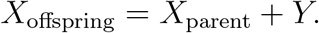

where *Y* is a random variable with probability distribution **K**_div_. In the sexual infinitesimal model, the trait of an offspring *X*_offspring_ with parents traits *X*_parent,1_ and *X*_parent,2_ is given by

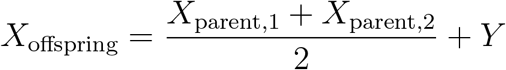

where *Y* is drawn from a centered normal distribution with variance **V**_div_/2.

Numerically, this model has a very high computational cost, especially when the number of individuals is large. As a consequence, we performed the simulations using an approximating model, by first fixing *dt* to a small but deterministic value. Then, for each individual, we draw a time of birth following the exponential law ℰ(***β***) and a time of death following the exponential law ℰ(***μ***(*X*_*i*_ −**ct**) +***κ****N*_*t*_/**N**). Then we simply count which individuals led to a reproduction event and which died on the time-window [**t, t** + *dt*]. This amounts to the supposition that on this time interval, individuals cannot reproduce more than once.

#### Deterministic approximation of the phenotypic distribution

We first compare our approximation of the phenotypic distribution with the empirical distribution of the IBM model for the scenarios described in Fig. 8. When the number of individuals is large (of order **N** = 10^4^), we see that our second order approximations are accurate and fit with the empirical distribution of the stochastic model (see Fig. S3).

#### Deterministic approximation of the effect of the changing speed

Here, we compare our approximation formula described in Table 2, with the outcomes of the stochastic model with a small number of individuals (**N** is equal to 10^2^ or 10^3^) in the various scenarios described in Fig. 3.

When the speed of change is slow compared to the critical speeds, our approximations seem accurate in the sense that the approximation error usually falls on our confidence intervals (see Fig. S4-S6). In the infinitesimal sexual model, our approximation also does well when the speed is close to the critical threshold. In this model, we know that the population adapt thanks to the bulk of the population, which moves forward. Thus, even if the number of individuals decreases, many individuals remain at the dominant trait. The number of individuals does not have a critical influence on the adaptation response.

However for the asexual model, when the speed increases, our approximations become less accurate. In this model, only the individuals near the optimal trait help the population to adapt. Thus when the speed increases, the proportion of individuals near the optimal trait decreases because the lag increases. Moreover, when the number of individuals decreases, the actual number of individuals at the optimal trait may be zero, which may lead to an additional burden, and possibly the extinction of the population before the critical value **c**_*c*_ is reached (Calvez et al., 2023). In particular, we see in Figures S4-S6 (a) that the mean fitness of the population drops below 0 for fifty percent of the simulations when the speed is close to the critical speed. Thus the effect of the population size is stronger for the asexual model than for the infinitesimal sexual model.

**Figure S3:**
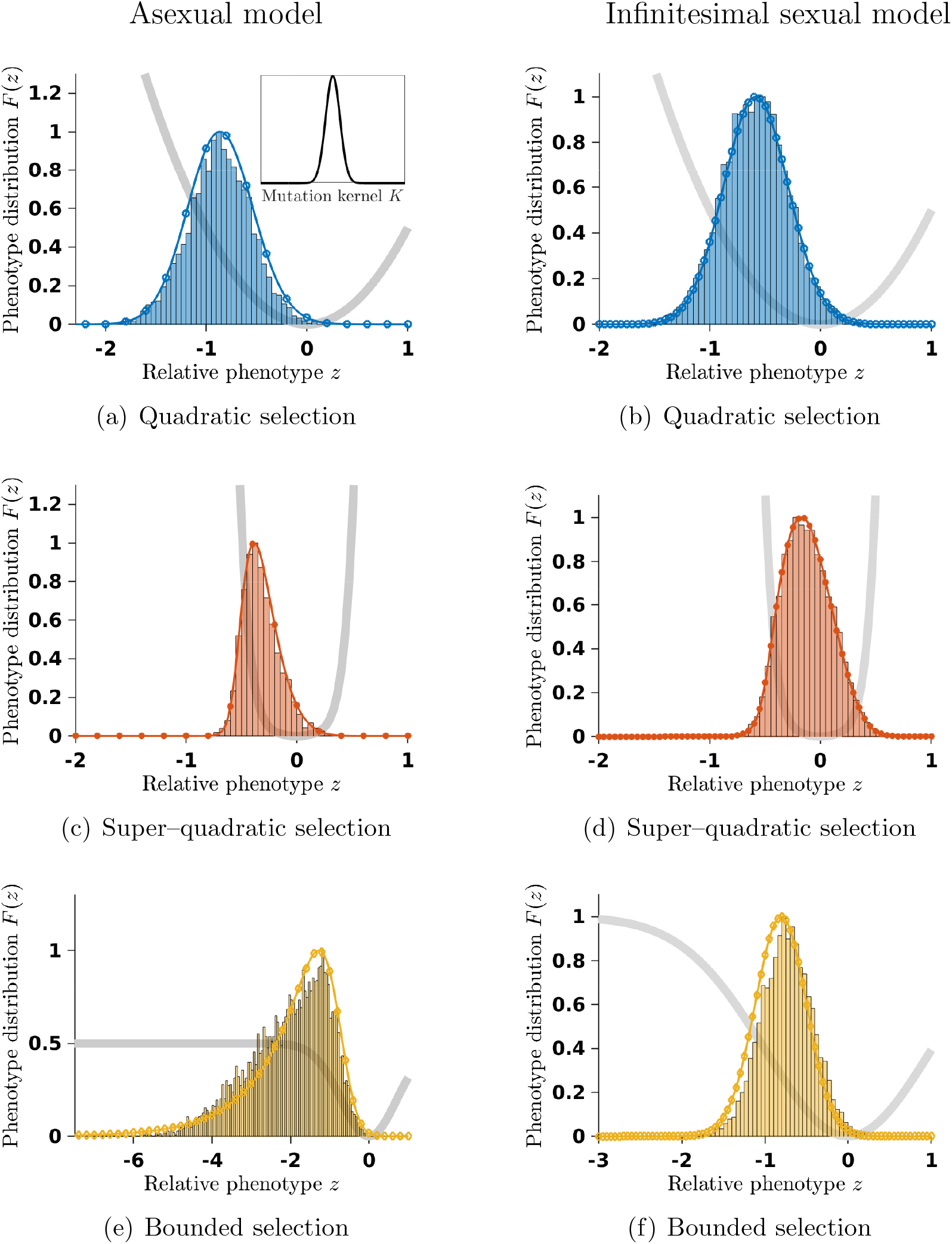
Mutation-selection equilibria **F** in changing environment with three different shapes of selection: (a)-(b) quadratic function *m*(*z*) = *z*^2^/2 (blue circled marked curves); (c)-(d) super-quadratic function *m*(*z*) = *z*^2^/2 + *z*^6^/64 (blue star marked curves); (e)-(f) bounded function *m*(*z*) = *m*_∞_(1 −exp(−*z*^2^/(2*m*_∞_)) (orange diamond marked curves). The speed of environment change is **c** = 0.09 in the asexual model while it is **c** = 0.05 in the infinitesimal sexual model so that it remains below the critical speeds **c**_*c*_ and **c**_tip_ and the distribution deviates significantly from the Gaussian distribution approximation. Other parameters are: ***β*** = 1, **V**_sel_ = 1 and **V**_div_ = 0.01 and *m*_∞_ = 0.5 in the asexual model and **V**_div_ = 0.1 and *m*_∞_ = 1 in the infinitesimal sexual model. We compare our analytical results (second order results plain marked curves) with the histogram of the stochastic model with **N** = 10^4^ individuals. For the asexual scenario, we used the Gaussian kernel.

**Figure S4:**
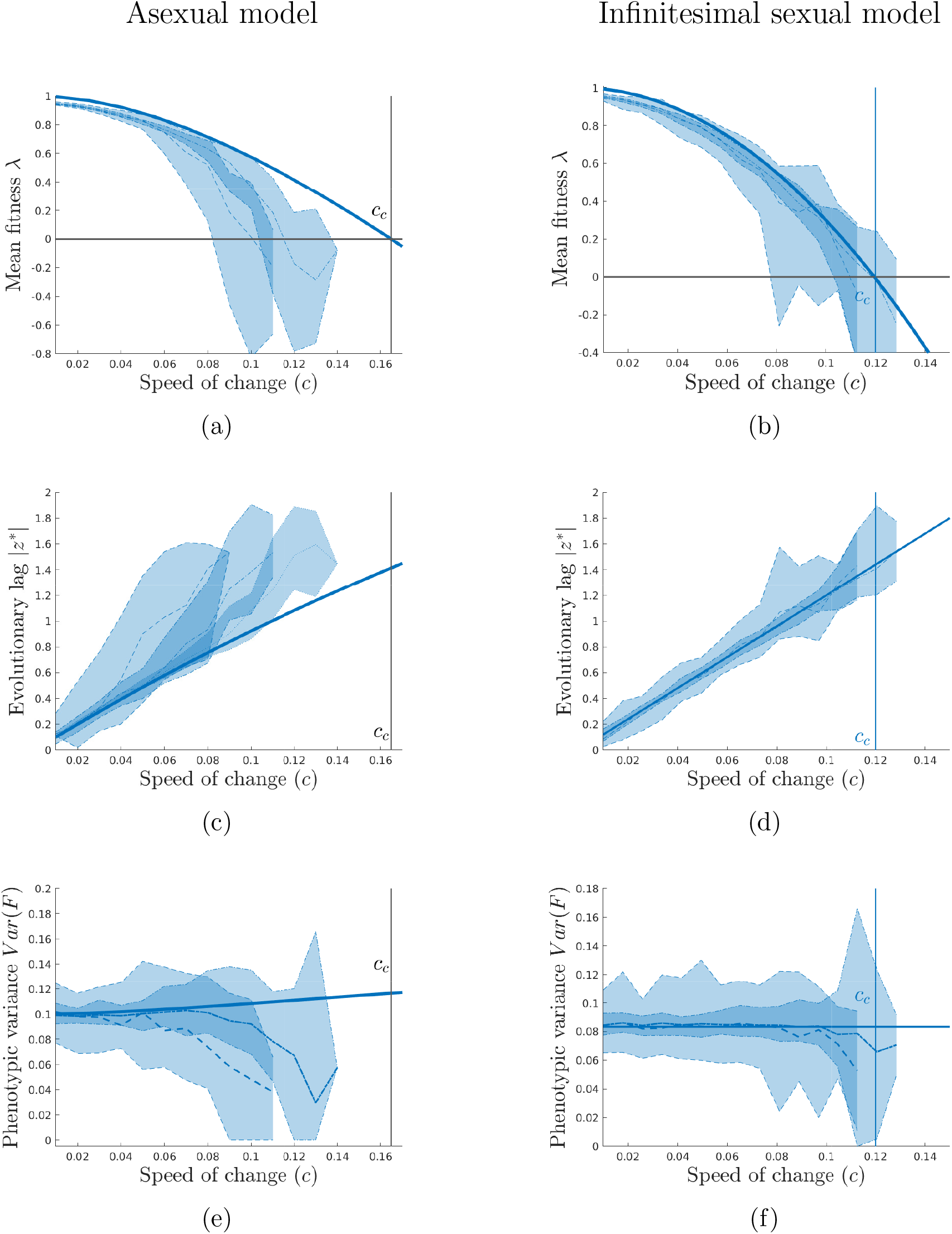
Influence of the speed of environmental change **c** for a population with finite number of individuals (**N** = 10^2^ dashed curves and **N** = 10^3^ dash-dotted curves) under quadratic selection *m*(*z*) = *z*^2^/2. Other parameters are: ***β*** = 1, **V**_sel_ = 1 and **V**_div_ = 0.01 in the asexual model and **V**_div_ = 0.1 in the infinitesimal sexual model. In the asexual model, the mutation kernel is Gaussian. The shade region corresponds to the 95% and 5% confidence intervals around the median. The plain curves correspond to the first order approximation in the asexual model and the second order approximation in the sexual infinitesimal model.

**Figure S5:**
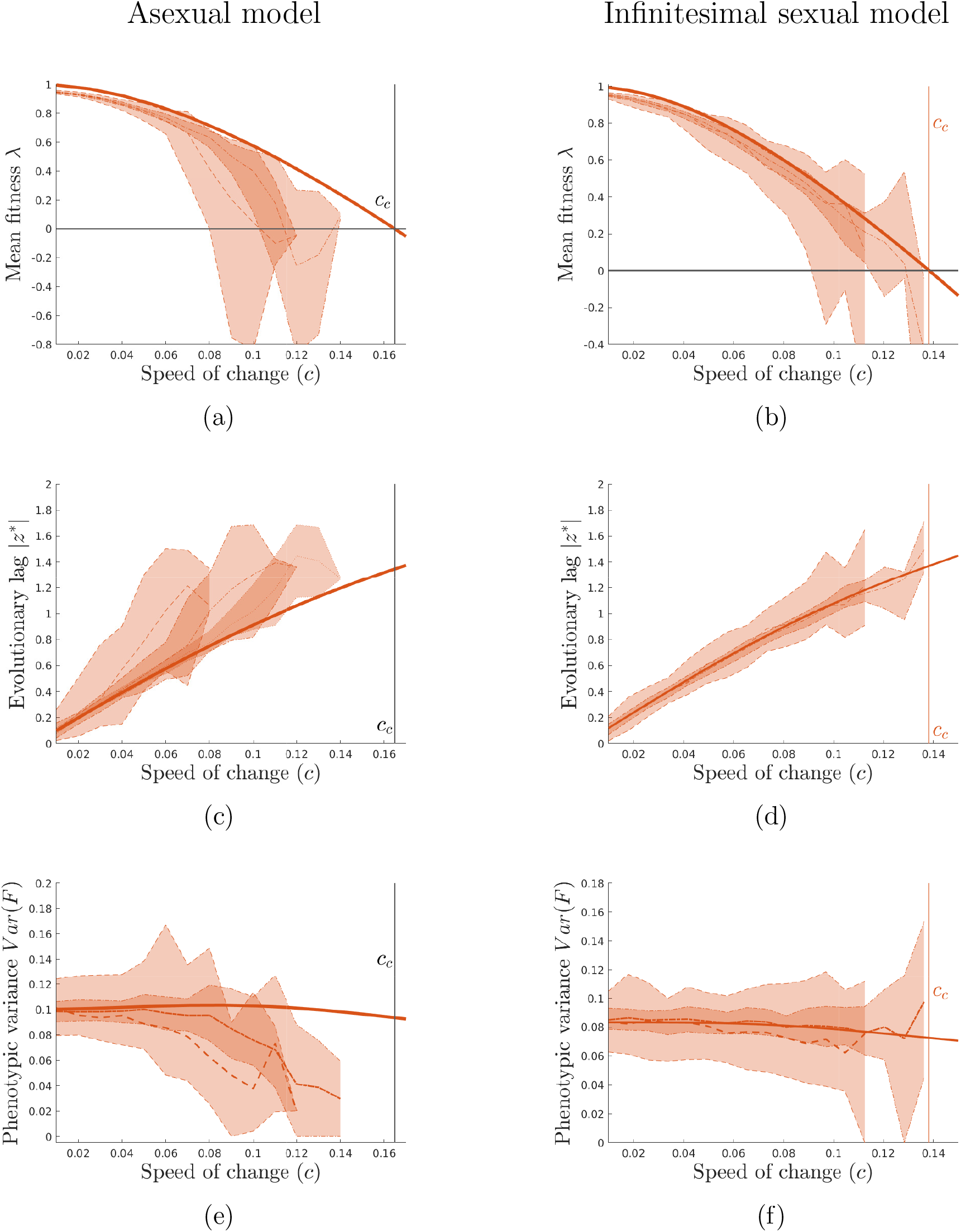
Influence of the speed of environmental change **c** for a population with finite number of individuals (**N** = 10^2^ dashed curves and **N** = 10^3^ dash-dotted curves) under super–quadratic selection *m*(*z*) = *z*^2^/2 + *z*^6^/64. Other parameters are: ***β*** = 1, **V**_sel_ = 1 and **V**_div_ = 0.01 in the asexual model and **V**_div_ = 0.1 in the infinitesimal sexual model. In the asexual model, the mutation kernel is Gaussian. The shade region corresponds to the 95% and 5% confidence intervals around the median. The plain curves correspond to the first order approximation in the asexual model and the second order approximation in the sexual infinitesimal model.

**Figure S6:**
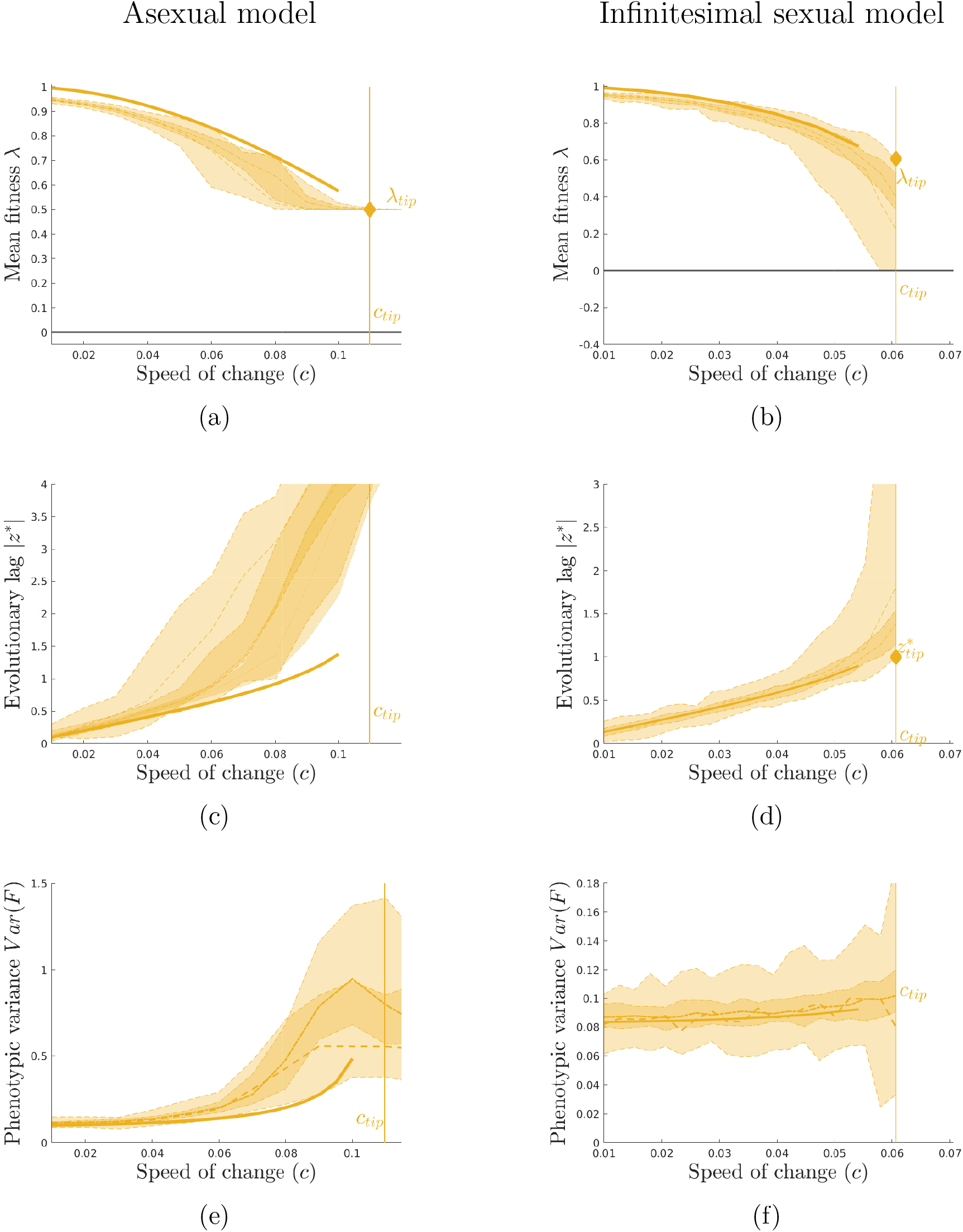
Influence of the speed of environmental change **c** for a population with finite number of individuals (**N** = 10^2^ dashed curves and **N** = 10^3^ dash-dotted curves) under bounded selection function *m*(*z*) = *m*_∞_(1 − exp(−*z*^2^/(2*m*_∞_)). Other parameters are: ***β*** = 1, **V**_sel_ = 1 and **V**_div_ = 0.01 and *m*_∞_ = 0.5 in the asexual model and **V**_div_ = 0.1 and *m*_∞_ = 1 in the infinitesimal sexual model. In the asexual model, the mutation kernel is Gaussian. The shade region corresponds to the 95% and 5% confident intervals around the median. The plain curves correspond to the first order approximation in the asexual model and the second order approximation in the sexual infinitesimal model.

## Notes

Fundings: The author(s) acknowledge support of the Institut Henri Poincaré (UAR 839 CNRS-Sorbonne Université), and LabEx CARMIN (ANR-10-LABX-59-01). JG acknowledges GLOBNETS project (ANR-16-CE02-0009) and ModEcoEvo project funded by the Univ. Savoie Mont-Blanc. This project has received funding from the European Research Council (ERC) under the European Union’s Horizon 2020 research and innovation programme (grant agreement No 865711). OR acknowledges support from the Peter Wall Institute of Advanced Studies and from the France Canada Research Funds.

### Competing Interest Statement

The authors have declared no competing interest.

### Summary of Updates

We clarified some points on the stochastic model and linked ou results with existing results on finte population size.

## References

Agrawal, A.F., Whitlock, M.C., 2010. Environmental duress and epistasis: how does stress affect the strength of selection on new mutations? Trends in Ecology & Evolution 25, 450–458. doi:10.1016/j.tree.2010.05.003.

Alexander, H.K., Martin, G., Martin, O.Y., Bonhoeffer, S., 2014. Evolutionary rescue: linking theory for conservation and medicine. Evolutionary applications 7, 1161–1179.

Barles, G., Mirrahimi, S., Perthame, B., 2009. Concentration in lotka-volterra parabolic or integral equations: A general convergence result. Methods and Applications of Analysis 16, 321–340. doi:10.4310/MAA.2009.v16.n3.a4.

Barton, N.H., Etheridge, A.M., Véber, A., 2017. The infinitesimal model. Theoretical Population Biology 118, 50–73. doi:10.1101/039768.

Barton, N.H., Keightley, P.D., 2002. Understanding quantitative genetic variation. Nature Reviews Genetics 3, 11–21. doi:10.1038/nrg700.

Barton, N.H., Turelli, M., 1987. Adaptive landscapes, genetic distance and the evolution of quantitative characters. Genetical Research 49, 157–173. doi:10.1017/S0016672300026951.

Barton, N.H., Turelli, M., 1989. Evolutionary quantitative genetics: how little do we know? Annual Review of Genetics 23, 337–370. doi:10.1146/annurev.ge.23.120189.002005.

Bulmer, M.G., 1971. The effect of selection on genetic variability. Amer. Nat. 105, 201–211.

Bulmer, M.G., 1974. Linkage disequilibrium and genetic variability. Genet. Res. 23, 281–289.

Bulmer, M.G., 1980. The Mathematical Theory of Quantitative Genetics. Oxford, Clarendon Press.

Burger, R., 1991. Moments, cumulants, and polygenic dynamics. J. Math. Biol. 30, 199–213.

Bürger, R., 2000. The Mathematical Theory of Selection, Recombination, and Mutation. Wiley Series in Mathematical & Computational Biology, Wiley.

Burger, R., Lynch, M., 1995. Evolution and Extinction in a Changing Environment: A Quantitative-Genetic Analysis. Evolution 49, 151–163. doi:10.2307/2410301.

Bürger, R., 1999. Evolution of Genetic Variability and the Advantage of Sex and Recombination in Changing Environments. Genetics 153, 1055–1069. doi:10.1093/genetics/153.2.1055.

Calvez, V., Forien, R., Méléard, S., 2023. in preparation.

Calvez, V., Garnier, J., Patout, F., 2019. Asymptotic analysis of a quantitative genetics model with nonlinear integral operator. J École Polytechnique — Mathématiques 6, 537–579. doi:10.5802/jep.100.

Calvez, V., Henderson, C., Mirrahimi, S., Turanova, O., Dumont, T., 2022a. Non-local competition slows down front acceleration during dispersal evolution. Annales Henri Lebesgue 5, 1–71. doi:10.5802/ahl.117.

Calvez, V., Henry, B., Méléard, S., Tran, V.C., 2022b. Dynamics of lineages in adaptation to a gradual environmental change. Annales Henri Lebesgue 5, 729–777. doi:10.5802/ahl.135.

Calvez, V., Lam, K.Y., 2020. Uniqueness of the viscosity solution of a constrained hamilton–jacobi equation. Calc. Var. 59. doi:10.1007/s00526-020-01819-0.

Champagnat, N., Ferriere, R., Méléard, S., 2006. Unifying evolutionary dynamics: from individual stochastic processes to macroscopic models. Theoretical population biology 69, 297–321.

Charlesworth, B., 1993. Directional selection and the evolution of sex and recombination. Genetical Research 61, 205–224. doi:10.1017/S0016672300031372.

Cloez, B., Gabriel, P., 2020. On an irreducibility type condition for the ergodicity of nonconservative semigroups. Comptes Rendus. Mathématique 358, 733–742.

Collins, S., de Meaux, J., Acquisti, C., 2007. Adaptive walks toward a moving optimum. Genetics 176, 1089–1099. doi:10.1534/genetics.107.072926.

Cotto, O., Ronce, O., 2014. Maladaptation as a source of senescence in habitats variable in space and time. Evolution 68, 2481–2493. doi:10.1111/evo.12462.

Coville, J., Hamel, F., 2019. On generalized principal eigenvalues of nonlocal operators witha drift. Nonlinear Analysis, 111569.

Dekens, L., 2020. Evolutionary dynamics of complex traits in sexual populations in a strongly heterogeneous environment: how normal? 2012.10115.

Dekens, L., Lavigne, F., 2021. Front propagation of a sexual population with evolution of dispersion: a formal analysis. doi:10.48550/ARXIV.2105.02523.

Dekens, L., Mirrahimi, S., 2021. Dynamics of dirac concentrations in the evolution of quantitative alleles with sexual reproduction. doi:10.48550/ARXIV.2111.14814.

Dekens, L., Otto, S.P., Calvez, V., 2021. The best of both worlds: combining population genetic and quantitative genetic models. doi:10.48550/ARXIV.2111.11142.

Diekmann, O., Jabin, P.E., Mischler, S., Perthame, B., 2005. The dynamics of adaptation: An illuminating example and a hamilton–jacobi approach. Theoretical Population Biology 67, 257 – 271. doi:http://dx.doi.org/10.1016/j.tpb.2004.12.003.

Dimassi, M., Sjostrand, J., 1999. Spectral Asymptotics in the Semi-Classical Limit. London Mathematical Society Lecture Note Series, Cambridge University Press.

Evans, L.C., 2010. Partial differential equations. American Mathematical Society, Providence, R.I.

Evans, L.C., Ishii, H., 1985. A PDE approach to some asymptotic problems concerning random differential equations with small noise intensities. Ann. Inst. H. Poincaré Anal. Non Linéaire 2, 1–20.

Feng, J., Kurtz, T.G., 2006. Large Deviations for Stochastic Processes. Mathematical Surveys and Monographs.

Fisher, R.A., 1918. The correlation between relatives on the supposition of Mendelian inheritance. Trans. R. Soc. Edinburgh 52, 399–433.

Fleming, W.H., 1977. Exit probabilities and optimal stochastic control. Appl. Math. Optim. 4, 329–346. doi:10.1007/BF01442148.

Fleming, W.H., 1979. Equilibrium distributions of continuous polygenic traits. SIAM J. Appl. Math. 36, 148–168. doi:10.1137/0136014.

Frank, S.A., Slatkin, M., 1990. The distribution of allelic effects under mutation and selection. Genetical Research 55, 111–117. doi:10.1017/S0016672300025350.

Freidlin, M.I., Wentzell, A.D., 1998. Random perturbations of dynamical systems. Springer.

Gauzere, J., Teuf, B., Davi, H., Chevin, L.M., Caignard, T., Leys, B., Delzon, S., Ronce, O., Chuine, I., 2020. Where is the optimum? predicting the variation of selection along climatic gradients and the adaptive value of plasticity. a case study on tree phenology. Evolution letters 4, 109—123. doi:10.1002/evl3.160.

Gomulkiewicz, R., Houle, D., 2009. Demographic and genetic constraints on evolution. Am. Nat. 174, E218–229.

Halligan, D.L., Keightley, P.D., 2009. Spontaneous mutation accumulation studies in evolutionary genetics. Annual Review of Ecology, Evolution, and Systematics 40, 151–172. doi:10.1146/annurev.ecolsys.39.110707.173437.

Hill, W.G., 2010. Understanding and using quantitative genetic variation. Philosophical Transactions of the Royal Society B: Biological Sciences 365, 73–85. doi:10.1098/rstb.2009.0203.

Iglesias, S.F., Mirrahimi, S., 2021. Selection and mutation in a shifting and fluctuating environment. Communications in Mathematical Sciences.

Johnson, T., Barton, N., 2005. Theoretical models of selection and mutation on quantitative traits. Philosophical Transactions of the Royal Society B: Biological Sciences 360, 1411–1425. doi:10.1098/rstb.2005.1667.

Jones, A.G., Bürger, R., Arnold, S.J., Hohenlohe, P.A., Uyeda, J.C., 2012. The effects of stochastic and episodic movement of the optimum on the evolution of the g-matrix and the response of the trait mean to selection. J. Evol. Biol. 25, 2210–2231. doi:https://doi.org/10.1111/j.1420-9101.2012.02598.x.

Keightley, P.D., Hill, W.G., 1988. Quantitative genetic variability maintained by mutation-stabilizing selection balance in finite populations. Genet. Res. 52, 33–43.

Kimura, M., 1965. A stochastic model concerning the maintenance of genetic variability in quantitative characters. Proc. Natl. Acad. Sci. USA 54, 731– 736. doi:10.1073/pnas.54.3.731.

Klausmeier, C.A., Osmond, M.M., Kremer, C.T., Litchman, E., 2020. Ecological limits to evolutionary rescue. Philosophical Transactions of the Royal Society B: Biological Sciences 375, 20190453. doi:10.1098/rstb.2019.0453.

Kopp, M., Hermisson, J., 2009. The genetic basis of phenotypic adaptation i: Fixation of beneficial mutations in the moving optimum model. Genetics 182, 233–249. doi:10.1534/genetics.108.099820.

Kopp, M., Matuszewski, S., 2014. Rapid evolution of quantitative traits: theoretical perspectives. Evolutionary Applications 7, 169–191.

Lam, K.Y., 2017. Stability of dirac concentrations in an integro-pde model for evolution of dispersal. Calc. Var. 59. doi:10.1007/s00526-017-1157-1.

Lam, K.Y., Lou, Y., 2017. An integro-pde model for evolution of random dispersal. J. Func. Anal. 272, 1755–1790. doi:10.1016/j.jfa.2016.11.017.

Lam, K.Y., Lou, Y., Perthame, B., 2022. A Hamilton-Jacobi Approach to Evolution of Dispersal. URL: http://arxiv.org/abs/2205.05534, xdoi:10.48550/arXiv.2205.05534. arXiv:2205.05534 [math].

Lande, R., 1975. The maintenance of genetic variability by mutation in a polygenic character with linked loci. Genetical Research 26, 221–235. doi:10.1017/S0016672300016037.

Lande, R., Shannon, S., 1996. The role of genetic variation in adaptation and population persistence in a changing environment. Evolution 50, 434–437. doi:10.2307/2410812.

Lange, K.J., 1978. Central limit theorems of pedigrees. J. Math. Biol. 6, 59–66.

Lorz, A., Mirrahimi, S., Perthame, B., 2011. Dirac mass dynamics in multidimensional nonlocal parabolic equations. Commun. Partial Differential Equations 36, 1071–1098.

Lynch, M., Gabriel, W., 1990. Mutation load and the survival of small populations. Evolution 44, 1725–1737. doi:https://doi.org/10.1111/j.1558-5646.1990.tb05244.x.

Lynch, M., Gabriel, W., Wood, A.M., 1991. Adaptive and demographic responses of plankton populations to environmental change. Limnology and Oceanography 36, 1301–1312.

Lynch, M., Lande, R., 1993. Evolution and extinction in response to environmental change. Sinauer Assoc.

Martin, G., Roques, L., 2016. The non-stationary dynamics of fitness distributions: Asexual model with epistasis and standing variation. Genetics 204, 1541–1558. doi:10.1534/genetics.116.187385.

Mirrahimi, S., 2017. A hamilton–jacobi approach to characterize the evolutionary equilibria in heterogeneous environments. Mathematical Models and Methods in Applied Sciences 27, 2425–2460. doi:10.1142/S0218202517500488.

Mirrahimi, S., Raoul, G., 2013. Population structured by a space variable and a phenotypical trait. Theor. Popul. Biol. 84, 87–103.

Mirrahimi, S., Roquejoffre, J.M., 2016. A class of hamilton–jacobi equations with constraint: Uniqueness and constructive approach. J. Diff. Equ. 260, 4717–4738. doi:https://doi.org/10.1016/j.jde.2015.11.027.

Nei, M., 2014. Mutation-Driven Evolution. Oxford University Press.

van Nes, E.H., Arani, B.M.S., Staal, A., van der Bolt, B., Flores, B.M., Bathiany, S., Scheffer, M., 2016. What Do You Mean, ‘Tipping Point’? Trends in Ecology & Evolution 31, 902–904. doi:10.1016/j.tree.2016.09.011.

Osmond, M.M., Klausmeier, C.A., 2017. An evolutionary tipping point in a changing environment. Evolution 71, 2930–2941. doi:10.1111/evo.13374.

Patout, F., 2019. Analyse asymptotique d’équations intégro-différentielles : modèles d’évolution et de dynamique des populations. Theses. Université de Lyon.

Patout, F., 2020. The cauchy problem for the infinitesimal model in the regime of small variance. ANalysis PDE doi:2001.04682.

Patout, F., Forien, R., Garnier, J., 2020. Ancestral lineages in mutationselection equilibria with moving optimum. arXiv:2011.05192.

Pease, C.P., Lande, R., Bull, J.J., 1989. A model of population growth, dispersal and evolution in a changing environment. Ecology 70, 1657–1664.

Perthame, B., 2007. Transport equations in biology. Frontiers in Mathematics, Birkhäuser Verlag.

Perthame, B., Barles, G., 2008. Dirac concentrations in Lotka-Volterra parabloci PDEs. Indiana Univ. Math. J. 457, 3275–3301.

Perthame, B., Souganidis, P.E., 2016. Rare mutations limit of a steady state dispersal evolution model. Math. Model. Nat. Phenom. 11, 154–166. doi:10.1051/mmnp/201611411.

Raoul, G., 2021. Exponential convergence to a steady-state for a population genetics model with sexual reproduction and selection. 2104.06089.

Rauch, J., 2012. Hyperbolic partial differential equations and geometric optics.volume 133. American Mathematical Society Providence, RI.

Rockafellar, R.T., 1970. Convex Analysis. Princeton landmarks in mathematics and physics, Princeton University Press.

Roques, L., Patout, F., Bonnefon, O., Martin, G., 2020. Adaptation in general temporally changing environments. SIAM J. Appl. Math. 80, 2420–2447. doi:10.1137/20M1322893.

Santiago, E., 1998. Linkage and the maintenance of variation for quantitative traits by mutation–selection balance: an infinitesimal model. Genet Res 71, 161–170. doi:10.1017/S0016672398003231.

Tufto, J., 2000. Quantitative genetic models for the balance between migration and stabilizing selection. Genetical research 76, 285–293.

Turelli, M., 1984. Heritable genetic variation via mutation-selection balance: Lerch’s zeta meets the abdominal bristle. Theor. Popul. Biol. 25, 138–193. doi:10.1016/0040-5809(84)90017-0.

Turelli, M., 2017. Commentary: Fisher’s infinitesimal model: A story for the ages. Theoretical Population Biology 118, 46 – 49. doi:https://doi.org/10.1016/j.tpb.2017.09.003.

Turelli, M., Barton, N.H., 1990. Dynamics of polygenic characters under selection. Theor. Popul. Biol. 38, 1–57. doi:https://doi.org/10.1016/0040-5809(90)90002-D.

Turelli, M., Barton, N.H., 1994. Genetic and statistical analyses of strong selection on polygenic traits: what, me normal? Genetics 138, 913–941.

W Hao, K-Y Lam Y.L., 2021. Ecological and evolutionary dynamics in advective environments: Critical domain size and boundary conditions. Disc. Conti. Dyn. Syst. - B 26, 367–400.

Walters, R.J., Blanckenhorn, W.U., Berger, D., 2012. Forecasting extinction risk of ectotherms under climate warming: an evolutionary perspective. Functional Ecology 26, 1324–1338.

Waxman, D., Peck, J.R., 1999. Sex and Adaptation in a Changing Environment. Genetics 153, 1041–1053. doi:10.1093/genetics/153.2.1041.

Zworski, M., 2012. Semiclassical analysis. volume 138. American Mathematical Society Providence, RI.

